# A synthetic cell phage cycle

**DOI:** 10.1101/2025.06.14.659709

**Authors:** Antoine Levrier, Paul Soudier, David Garenne, Ziane Izri, Steven Bowden, Ariel B. Lindner, Vincent Noireaux

## Abstract

Viral infection of living cells, exemplified by bacteriophage interaction with bacteria, is fundamental to biology and universal across living systems. Here, we establish an all-cell-free viral cycle where T7 phages infect synthetic cells, equipped with lipopolysaccharides on the outer leaflet of the lipid membrane while encapsulating a cell-free gene expression system. We track each cycle step to demonstrate T7 phage-specific adsorption onto the liposomes, genome ejection, replication, expression, and assembly of new infectious virions within the synthetic cells. We quantify key characteristics of the cycle, including the multiplicity of infection, replication efficiency, liposome size constraints, and phage rebinding dynamics. This work establishes a versatile, fully defined in vitro platform for reconstructing and investigating viral infections from individual molecular components.

## Main text

Building biological systems from their molecular constituents enables isolating, quantifying, and characterizing biochemical mechanisms often entangled and thus difficult to probe *in vivo*. It is now possible to reconstitute complex biological processes in cell-free systems across multiple scales^1–3^. It offers a testbed for validating and enriching our understanding of life’s fundamental principles and constructing synthetic life systems^4,5^. Among these challenges, establishing a phage cycle is particularly relevant due to the critical role of viral processes in biology. As Arthur Kornberg noted fifty years ago, “Despite the extraordinary complexity of the viral life cycle, its dissection and eventual reconstruction with chemically defined components in a cell-free system from start to finish remains an attractive prospect”^6^. Strikingly, despite our extensive knowledge of phage evolution^7–9^, ecology^10–12^, and molecular mechanisms^13–15^, a complete cell-free phage infection cycle based on a synthetic cell system has not been achieved, underscoring both conceptual and technical frontiers.

Previous studies have provided isolated elements toward this goal. First, phage φX174 genome injection was achieved *in vitro* in the presence of a soluble receptor^16^. Later, the synthesis of infectious φX174 phages from purified components was demonstrated outside cells^17^, and its genome was synthetically assembled^18^. Further, phage T5 genome ejection into liposomes displaying the FhuA receptor^19–21^ showed that phages can be interfaced with single lipid bilayers via their receptor. Phage T7 infects *Escherichia coli* and is among the most characterized phages^22,23^. Every step of its infection cycle has been studied^24^, making it a relevant phage for assembling a synthetic, cell-free viral cycle. T7 is particularly adequate as a model system due to its obligate lytic cycle, minimal reliance on host factors^25^, and ease of genome engineering^26,27^. Previous studies have demonstrated the synthesis of infectious T7 phages in cell-free gene expression (CFE) systems^28^ and genome ejection in the presence of purified LPS (lipopolysaccharide)^29^. However, in contrast to phage T5, T7 genome ejection into liposomes via its LPS receptor has yet to be achieved. Furthermore, no study has integrated all the steps of a phage infection cycle into a single, self-contained framework.

Here, we set out to build a cell-free phage infection cycle (fig. 1A) composed of the phage T7 and synthetic cells (SCs) (fig. 1B). Despite the remarkable simplification compared to the *in vivo* framework, this synthetic platform reconstitutes all the steps of a phage cycle.

**Fig. 1.**
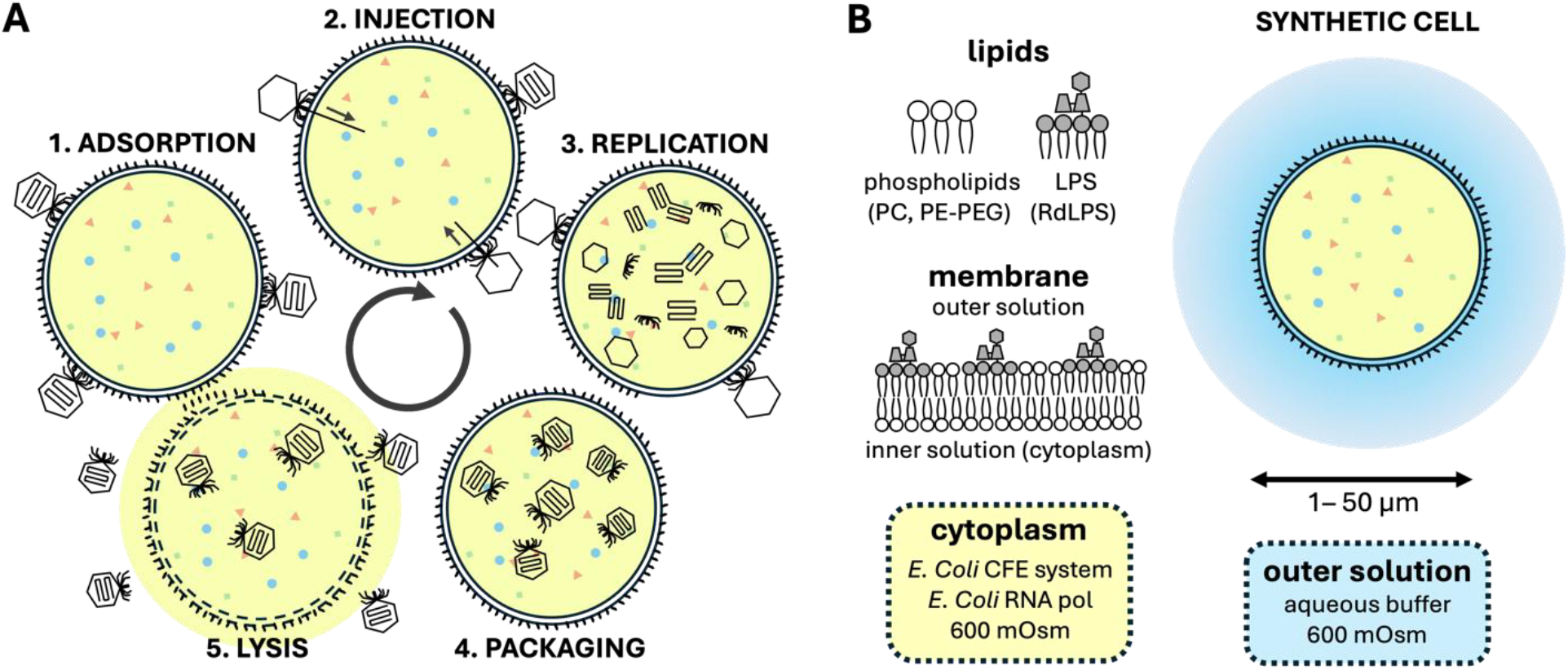
Schematic representation of the phage infection cycle in SCs. **(A)** The goal is to recapitulate key infection steps: phage adsorption, DNA ejection across the liposome bilayer, genome expression and replication, virion assembly, and SC lysis, releasing progeny phages into the external solution. (**B)** Schematic representation of RdLPS SC composition. The SCs consist of liposomes made from phospholipids (POPC and PE-PEG5000) and RdLPS. The RdLPS are exposed on the outer leaflet to enable phage T7 binding and subsequent infection. A cell-free expression (CFE) system devoid of DNA, RNA, and phage DNA and proteins is encapsulated into the liposome to allow for phage genome expression upon DNA ejection, initiating replication and virion production.

### Phage T7 binds specifically to SCs displaying rough LPS

We hypothesized that cell-sized single-bilayer liposomes presenting rough LPS on the outer leaflet and encapsulating a CFE system would enable phage T7 binding, genome ejection, genome expression, and subsequent synthesis of phage particles (fig. 1A). This previously untested hypothesis assumes that rough LPS embedded within a single lipid membrane is a sufficient T7 receptor for genome ejection into the SCs. However, creating CFE SCs incorporating rough LPS is challenging. LPS have high molecular weights and poor solubility compatibility with phospholipids, and they must remain stable in the membrane without compromising CFE inside the SCs^30,31^. Moreover, LPS must be incorporated into the outer leaflet while minimizing their presence in the inner leaflet to prevent unintended genome ejection inside the SCs. Accordingly, we opted for the RdLPS (fig. 1A), a short rough LPS type (fig. S1), and a cognate T7 phage tail fiber mutant that can, unlike wild-type T7 phage (T7 WT), interact with RdLPS *in vitro* (fig. S1). We assumed that our choice of RdLPS, one of the smallest LPS variants of *E. coli*, would enable stable integration into SCs, due to its greater hydrophobicity compared to larger LPS targets^32,33^.

Cell-sized liposomes loaded with a CFE reaction were prepared by emulsion transfer^34,35^ of a ternary lipid mixture comprised of phosphatidylcholine (POPC), phosphatidylethanolamine polyethylene glycol (PE-PEG), and RdLPS (fig. 2A, fig. S2). The presence of PE-PEG was critical for preventing aggregation and facilitating the formation of stable liposomes. We explored the membrane composition while maintaining a total lipid concentration of 100 μM (fig. 2B, fig. S3). A 55/15/30 mol% PC/PE-PEG/LPS mixture was chosen as it yielded a homogeneous and stable population of RdLPS-liposomes with diameters ranging from 1 μm to 50 μm. Notably, this ratio is consistent with reported LPS molar ratios in the *E. coli* outer membrane^36^.

**Fig. 2.**
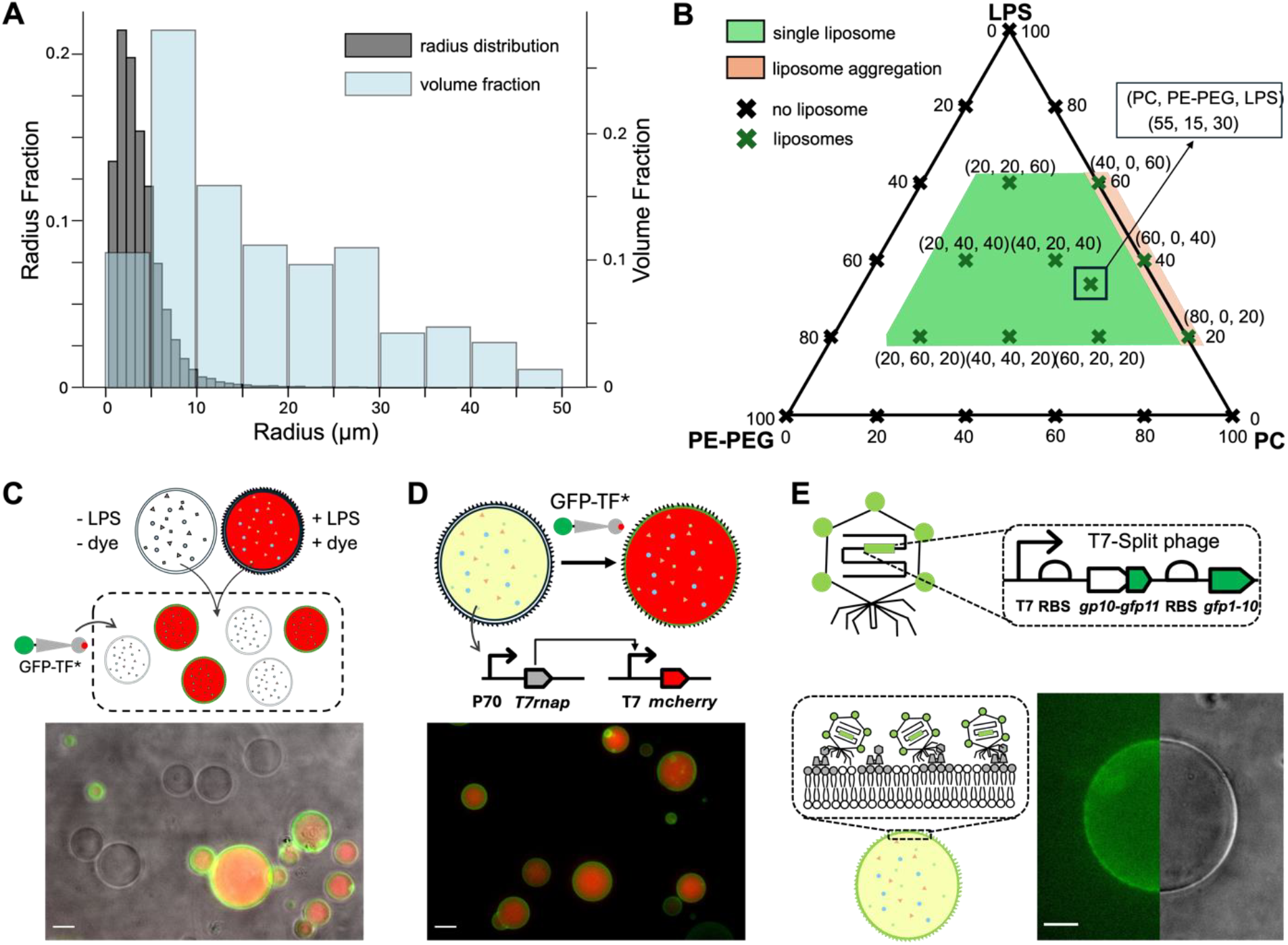
Prototyping synthetic cells with functional RdLPS for phage T7 infection. **(A)** Size and volume distribution of RdLPS SCs generated in this study. The SCs exhibit polydispersity, with an average radius of 3.7 μm. Liposomes smaller than 20 μm in radius account for 65% of the total encapsulated CFE reactions. **(B)** Ternary phase diagram of SC lipid composition. The green region indicates the lipid mixtures that successfully form SCs at a total lipid concentration of 100 μM. Liposomes do not form in the absence of RdLPS and POPC, while liposomes lacking PE-PEG tend to aggregate. **(C)** Selective detection of RdLPS-containing liposomes. Two populations of liposomes, with and without RdLPS, were prepared separately and subsequently mixed in an outer solution containing GFP-TF*, a fluorescent RdLPS-binding probe derived from the tip of the tail fiber of a T7 phage mutant. Microscopy images confirm that GFP-TF* selectively binds to RdLPS liposomes. Scale bar: 20 μm. **(D)** A linear DNA circuit encoding a *mcherry* reporter gene under a T7 promoter and a transcriptional activation cascade (P70a-*T7rnap*) is mixed with the CFE reaction and encapsulated into RdLPS SCs. Microscopy images show red fluorescence inside the liposomes, confirming active CFE, while GFP-TF* fluorescence at the membrane confirms the presence of functional RdLPS on the SC surface. Scale bar: 20 μm. **(E)** Adsorption of fluorescent T7-Split-S* phages onto RdLPS SCs. Top: Schematic of the T7-Split-S* system, in which the *gfp11* gene fragment is fused to the N-terminus of the T7 capsid gene (gene *gp10B*, supplementary text). The complementary *gfp1-10* gene fragment is expressed downstream of gene *10B-gfp11* under a ribosome binding site, allowing fluorescence upon phage production. Bottom: Microscopy image of RdLPS SCs incubated with T7-Split-S* in the outer solution. Green fluorescence localized at the membrane indicates the specific binding of T7-Split-S* to RdLPS-rich SCs. Liposome image is split between fluorescence (left) and phase contrast (right). Scale bar: 20 μm. Experimental details are presented in methods and tables S1-4.

Incorporation of the RdLPS into the outer leaflet of the lipid membrane and its expected function were assayed with a chimeric protein (GFP-TF*) consisting of superfolder GFP fused to the N-terminal of the tip of the T7 RdLPS-specific tail fiber mutant (TF*)^37^ (fig. S4). Two liposome populations (PC/PE-PEG and PC/PE-PEG/RdLPS) encapsulating CFE reactions were prepared separately. A red fluorescent dextran marker was added to the CFE reaction encapsulated within the RdLPS liposomes. Upon addition of GFP-TF* to a mixture of both populations, the fluorescence localized exclusively to RdLPS-containing membranes, confirming specific binding and stable LPS incorporation (fig. 2C, fig. S5-8). We confirmed the asymmetrical localization of the RdLPS on the outer leaflet of the lipid membrane (fig. S9-11, supplementary text). The presence of RdLPS in the membrane did not inhibit CFE gene expression activity inside the liposomes (fig. 2D, fig. S12-13).

To further validate this system, the T7-Split-S* phage was engineered from T7 WT using PHEIGES ^27^ to encode RdLPS-specific tail fibers and to tag capsid proteins fluorescently (fig. 2E, fig. S14, supplementary text). A T7-Split-S* titer of 10^11^ PFU/mL added to the outer solution of the RdLPS SCs resulted in a clear fluorescent phage localization at the membrane. The adsorption rate of an RdLPS-specific phage onto RdLPS SCs was estimated to be 5.10^−6^ min^-1^ (fig. S14), faster than reported values for bacterial cells (3.10^−9^ min^-138^). These findings confirmed RdLPS-specific phage adsorption onto RdLPS SCs.

### T7 genome is ejected, expressed, and replicated into SCs

Bacteriophages eject their genomes into bacterial hosts through diverse mechanisms that have been explored theoretically and experimentally^39–41^. The genome ejection process, partially driven by internal capsid pressure, has been characterized *in vivo* for several bacteriophages^42,43^. For instance, phage T5 can eject its genome into single-bilayer liposomes displaying the FhuA protein receptor via the tip of the long non-contractile tail that crosses the lipid membrane^19^. In contrast, T7’s short tail prevents T7 from crossing the cell membranes. Inner core proteins are ejected and assemble a trans-membrane channel to translocate the genome into the cell^44^. Additionally, T7 requires its RNA polymerase to fully translocate its genome^45^. In a buffer solution, T7 spontaneously releases its genome upon incubation with purified rough LPS due to conformational changes in the binding region^29^. To track both genome internalization and subsequent gene expression inside the SCs, we engineered two T7 phage variants, either the WT or RdLPS-specific tail fibers (T7-mC-WT, T7-mC-S*), each carrying a *mcherry* reporter gene downstream of the major capsid gene under the control of a T7 promoter (fig. 3A). We hypothesized that, as observed in bacterial infections, genome ejection into SCs would lead to the expression of the T7 RNA polymerase, thereby initiating a cascade of *mcherry* gene expression. As designed, the addition of T7-mC-S* phages to the external solution of the RdLPS SCs, encapsulating a DNA-free CFE reaction, resulted in a red fluorescence signal accumulation inside the liposomes (fig. 3B, S15-16, movie S1), indicating successful genome ejection and the expression of T7 genes. The addition of GFP-TF* to the infected RdLPS SCs revealed that while LPS were homogeneously distributed across liposomes, not all liposomes were infected (fig. 3C). Control experiments using T7-mC-WT, T7-WT, and T7-S* did not yield fluorescence above background levels, confirming that genome internalization was RdLPS-dependent (fig. 3D, fig. S17).

**Fig. 3.**
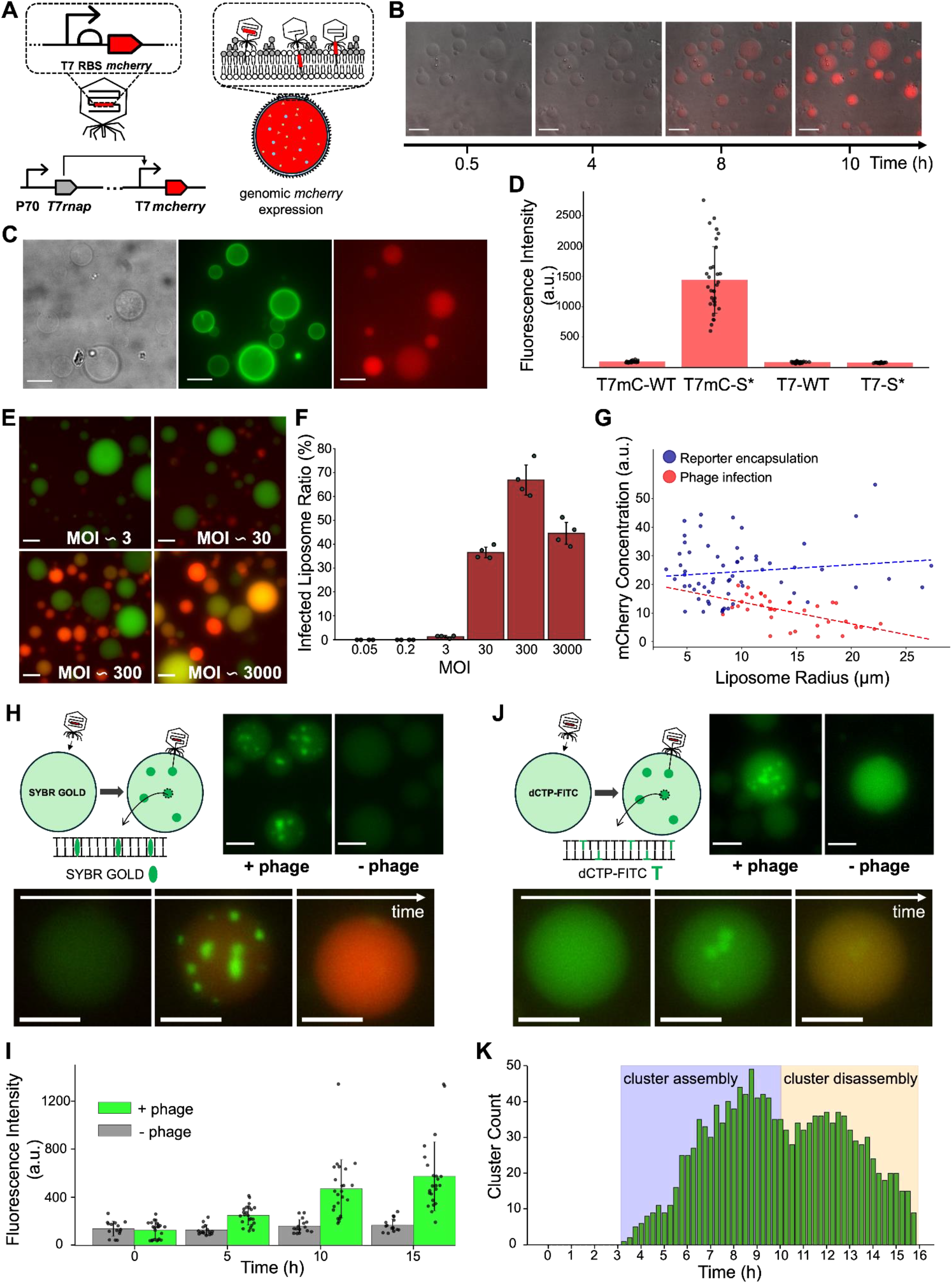
Genome Ejection and Expression of T7-mC-S in RdLPS SCs. **(A)** Schematic representation of the T7-mC-S* phage genome ejection and subsequent gene expression within RdLPS SCs. The T7-mC-S* phage carries a *mcherry* reporter gene inserted downstream of the major capsid protein gene (*gp10*) under a T7 promoter. Following adsorption onto RdLPS SCs, the phage ejects its genome across the lipid bilayer into the inner CFE reaction, where transcription and translation occur. The encapsulated CFE reaction is initially devoid of phage DNA, RNA, or proteins. **(B)** Time-lapse fluorescence microscopy images of RdLPS SCs exposed to T7-mC-S* phages (MOI = 300) in the outer solution at 30°C. The emergence of red fluorescence over time indicates successful phage genome ejection, transcription, and translation of mCherry inside the SCs. Scale bar: 40 μm. **C**. 100x magnification fluorescence microscopy image of an RdLPS SC infected by T7-mC-S*. GFP-TF* (green) marks the RdLPS membrane, while mCherry (red) indicates phage genome expression. Scale bar: 10 μm. **(D)** Quantification of mCherry fluorescence intensity in RdLPS SCs incubated with different phage variants: T7-mC-WT, T7-mC-S*, T7 WT, and T7-WT-S*. T7-mC-WT and T7-WT cannot bind RdLPS, while T7-WT-S* lacks the *mcherry* reporter gene. The results confirm that T7-mC-S* specifically infects RdLPS SCs and drives *mcherry* expression. **(E)** Representative fluorescence microscopy images showing RdLPS SCs infection at different MOIs. Images were taken after 15 hours of incubation at 30°C. Dextran-FITC (green) was used as a volume marker to facilitate liposome quantification. Scale bar: 25 μm. **(F)** Proportion of infected SCs at different MOIs, calculated from four replicate experiments. **(G)** Scatter plot of mCherry concentration as a function of liposome radius. Red points represent liposomes infected by T7-mC-S*, while blue points correspond to liposomes expressing *mcherry* from an encapsulated genetic circuit (P70a-*T7rnap* and T7-*mcherry* at 0.1 nM and 1 nM). Linear fits indicate that mCherry concentration remains constant in SCs with pre-encapsulated circuits but decreases with increasing liposome radius upon T7-mC-S* infection. **(H)** Detection of phage genome ejection using SYBR Gold, a DNA-intercalating dye. Left: Schematic of an RdLPS SC encapsulating CFE and SYBR Gold. Right: Microscopy images of SCs containing TXTL and SYBR Gold incubated with (MOI 100) or without T7-S* phages. The green fluorescence of SYBR Gold indicates the presence of phage DNA inside SCs. Bottom: Time-lapse images of the same liposome at 4, 6.5, and 16 hours after incubation with T7-mC-S*, showing progressive genome ejection (green, SYBR Gold fluorescence) followed by *mcherry* expression (red). Scale bar: 20 μm. **(I)** Bar graph showing the evolution of green fluorescence intensity over time in SCs encapsulating SYBR Gold with (green) or without (gray) T7-S* phages (MOI 100) in the outer solution. **(J)** Detection of phage genome replication using FITC-dCTP, a fluorescent nucleotide incorporated into newly synthesized DNA. Left: Schematic of an RdLPS SC encapsulating CFE and FITC-dCTP. Right: Microscopy images of SCs containing TXTL and FITC-dCTP incubated with (MOI 100) or without T7-S* phages. Green fluorescence indicates newly synthesized phage DNA. Bottom: Time-lapse images of the same liposome over 6, 10, and 16 hours after incubation with T7-mC-S*, showing FITC-dCTP fluorescence (green) and subsequent *mcherry* expression (red). Scale bar: 20 μm. **(K)** Quantification of FITC-dCTP fluorescence clusters inside RdLPS SCs over time in four replicate experiments. Cluster counts represent the total number of observed replication foci per frame across replicates. Experimental details are presented in methods and tables S1-2.

The rate of infection depended on the multiplicity of infection (MOI), defined as the ratio of plaque-forming units (PFU) to SCs (fig. 3E, fig. S18-19, movie S2). At an MOI of approximately 3, fewer than 2% of SCs exhibited fluorescence. In contrast, nearly 70% of SCs were infected at an MOI of 300 (fig. 3F). Quantitative analysis showed that, at constant phage concentrations in the outer solution, smaller SCs (<10 μm in diameter) exhibited greater fluorescence intensity than larger ones (>15 μm in diameter) (fig. 3G). This suggests that the efficacy of genome ejection depends on the surface-to-volume ratio, where smaller SCs, despite fewer genome ejections, maintain a greater genomic concentration relative to their volume.

We visualized genome ejection into the SCs using a DNA intercalating fluorescent dye that does not inhibit CFE (fig. 3H, fig. S20-22, movie S3). Upon the addition of T7-S* (MOI ≈ 100) in the outer solution, an increase in fluorescence intensity inside the SCs was observed along with the formation of fluorescent clusters (fig. 3I, fig. S22), indicating genome ejection. Over time, these clusters dissipated as mCherry fluorescence increased, suggesting genome replication and subsequent packaging^46,47^. The observed cluster formation and disappearance likely reflect concatemer formation and processing during phage packaging. In addition, T7 genome replication was visualized with a fluorescent nucleotide encapsulated within the SCs to further understand the cluster dynamics. Consistent with the DNA intercalating agent results, green fluorescent clusters formed over time, followed by *mcherry* expression (fig. 3J, fig. S23-26, movie S4). Cluster formation and subsequent dissipation were quantified, indicating genome replication and phage packaging, leading to the transition from heterogeneous to homogeneous fluorescence in SCs (fig. 3K, fig. S25-27).

### Phage T7 infection and synthesis can be kinetically tracked in SCs

The dynamics of phage T7 infection and synthesis in SCs were captured by following the kinetics of the three fluorescent probes (fig. 4A, fig. S28). Signals from the DNA intercalating dye and mCherry appeared within the first two hours. Fluorescence of double-stranded DNA clusters started between 4–5 hours, followed by cluster disassembly between 9–10 hours. From these fluorescence kinetics, we infer that phage adsorption and genome ejection occur within the first two hours, with genome expression initiating around the same time and reaching a characteristic duration of approximately 8–9 hours. DNA replication initiates at around 4–5 hours, consistent with the expected expression timing of T7 replication proteins. Synthesis of infectious phage starts within 4–9 hours.

**Fig. 4.**
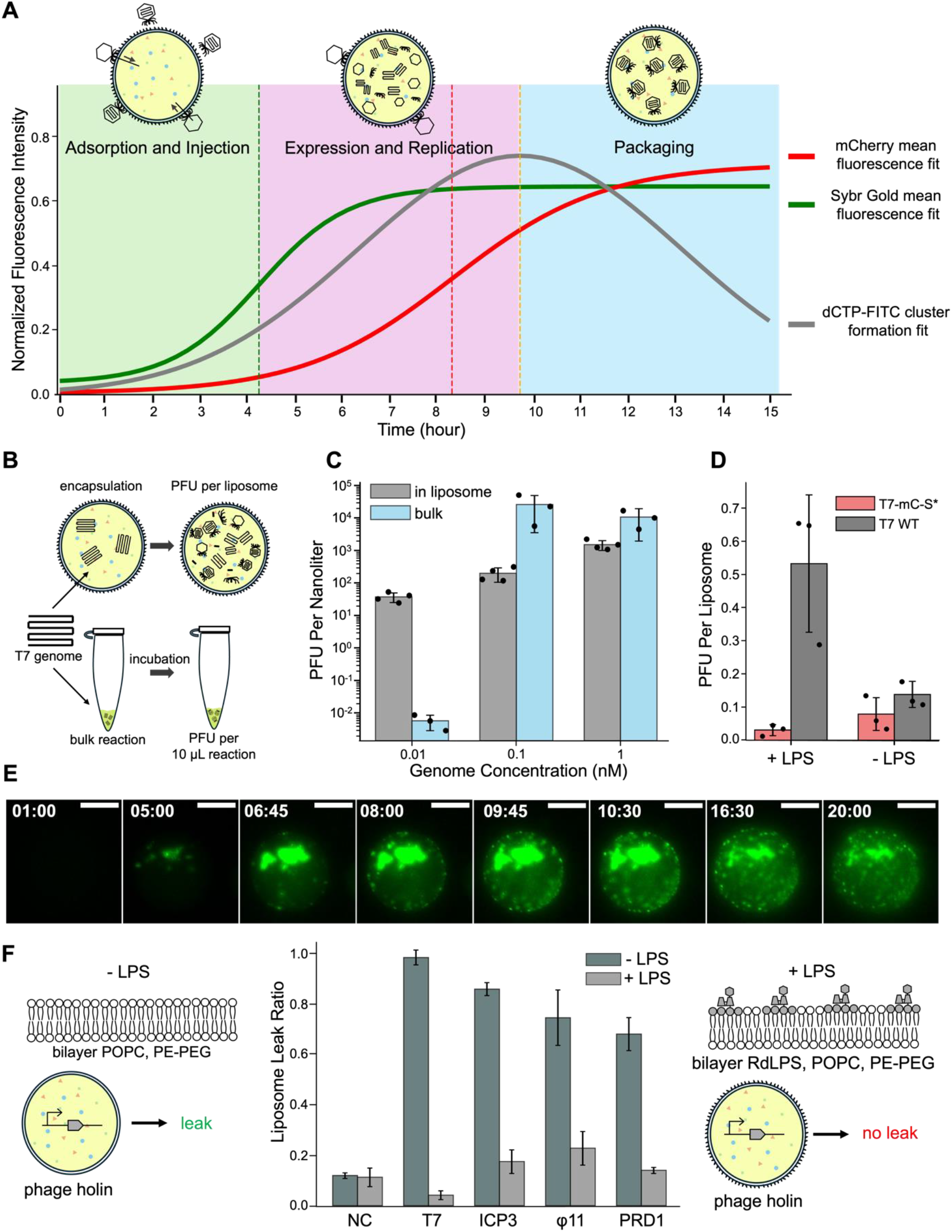
Production and Release of Phages in RdLPS SCs. **(A)** Kinetic analysis of genome ejection, replication, and expression following T7-mC-S infection. The Green sigmoidal fit represents the normalized average SYBR Gold fluorescence intensity within liposomes over time. Blue bars with a Gaussian fit indicate the timeframe during which initial dCTP cluster formation is observed. Red data points with a sigmoidal fit show the normalized mCherry fluorescence intensity inside liposomes. This multi-channel plot provides insights into the temporal dynamics and characteristic times of T7-mC-S* infection in RdLPS SCs. **(B)** Bulk vs. liposome-encapsulated T7 phage assembly. T7 phage DNA is mixed into the CFE reaction and either incubated in bulk (test tube) or encapsulated within an RdLPS SCs. After incubation, the bulk reaction is diluted, and the liposomes are lysed. PFU quantification is performed using a plaque assay. **(C)** Comparison of T7 WT phage assembly efficiency in bulk vs. liposome-encapsulated CFE reactions. Phage assembly efficiency is evaluated at three different initial DNA concentrations. The total TXTL volume in SCs is estimated from the liposome size distribution (Methods). **(D)** Impact of RdLPS on T7 WT and T7-mC-S phage assembly. Genomes are mixed at 0.01 nM in the CFE reaction. PFU per liposome is estimated based on microscopy liposome counts and total PFU quantified via plating. Comparison is made between liposomes with and without RdLPS. **(E)** Time-lapse microscopy images of RdLPS SCs infected with T7-Split-S*. Green fluorescence indicates the production and localization of the fluorescent capsid protein (gp10b-GFP11 + GFP1-10). Fluorescent capsids initially cluster near the site of genome replication and later disperse throughout the SCs as packaging into new phage particles progresses. Scale: 25 μm; time in hours. **(F)** Holin-induced membrane leak in RdLPS *vs*. LPS-less liposomes. Red rhodamine-dextran dye (3 kDa) is mixed with CFE along with a gene circuit expressing a bacteriophage holin gene. Upon holin expression, pores form in LPS-less liposome bilayers, causing dye leakage. In contrast, LPS-rich membranes exhibit limited dye leakage, suggesting a protective effect of RdLPS on membrane integrity. The bar plot quantifies the leak ratio of rhodamine-dextran dye after 3 hours of holin gene expression from different phages (T7, phi11, ICP3, and PRD1) at 30°C. Experimental details are presented in methods and tables S1-4.

To investigate phage assembly within the SCs, we first encapsulated the T7 WT genome at various concentrations into RdLPS-coated SCs and compared, after incubation and lysis by osmotic shock, the phage titers to those obtained from batch 10-μl CFE reactions (fig. 4B, fig. S29-30). Infectious phages were detected in both batch and SCs across all tested genome concentrations. At the lowest DNA concentration (10 pM), phage synthesis was more prevalent inside the SCs than in batch mode, likely due to favorable colocalization of gene expression machinery under confinement as compared to batch reactions. However, at higher concentrations, phage titers were comparable between the two systems, with slightly lower titers observed in SCs (fig. 4C). Next, to assess the effect of RdLPS incorporation on phage titer, we encapsulated T7-WT or T7-mC-S* genomes (10 pM) inside SCs with or without RdLPS into the membrane. After incubation and lysis by osmotic shock, the PFU titers per SC were estimated. While similar PFU per SC were obtained for liposomes lacking RdLPS, a 94 ± 4% reduction of infectious RdLPS-susceptible T7-mC-S* phages was observed in RdLPS-containing SCs (fig. 4D). These findings confirm that infectious phage particles are assembled inside LPS SCs. However, phage titers are reduced for RdLPS-susceptible phages, likely attributed to rapid rebinding to RdLPS and inactivation by genome ejection.

To track phage assembly inside RdLPS SCs, we incubated SCs with T7-Split-S* at an MOI of 100. At this MOI, no detectable fluorescence was observed at the membrane during adsorption. However, once the T7-Split-S* genome entered the SCs, the capsid protein fused to GFP11, and the complementary GFP1-10 were concurrently expressed to form fluorescent capsid proteins as a result of split-GFP complementation. We observed large fluorescent clusters, consistent with capsids self-assembly and binding to the genomic concatemer replicons to package the genomes. Further, this approach enabled visualizing genome concatemer packaging maturation into fluorescent phage capsids. Initially, green fluorescence clusters appeared inside the liposomes within the characteristic time of DNA ejection and replication (fig. 4A, movie S5), indicating fluorescent capsids localized near DNA concatemers (fig. 4E, fig. S31). Over time, the clusters dispersed into smaller fluorescence foci, suggesting concatemer processing into progeny phage particles.

### Phage Release via Osmotic Shock

At the final step of the lytic cycle, phages induce lysis of the host once progeny phages are assembled. In bacterial hosts, lysis is first mediated by a lysis cassette, which is typically expressed late in infection and consists of endolysins (which degrade the cell wall), holins (which form pores into the inner membrane), and spanins (which fuse the inner and outer membranes)^48,49^. This process results in spheroplast formation that spontaneously lyse due to osmotic shock^50^. Unlike bacteria, which experience lysis in a hypotonic environment, our SCs are incubated in isotonic conditions, making the phage lysis cassette insufficient, by construction, to induce liposome rupture. Indeed, no significant lysis was observed when phage genomes were expressed inside the RdLPS SCs. We investigated LPS-carrying SCs as a model for the bacterial outer membrane, and SCs devoid of LPS as a model for the bacterial inner membrane. CFE of the T7 holin into PC/PE-PEG SCs led to significant membrane disruption, as evidenced by leakage of a co-encapsulated dextran dye (fig. 4F). However, in PC/PE-PEG/RdLPS SCs, no dye leakage was observed, indicating that RdLPS inhibits holin-mediated membrane disruption (fig. 4F, movie S6). Similar results were obtained with holins from other bacteriophages, supporting that LPS prevents holin activity in this system, which is consistent with its natural function in forming pores in the LPS-less inner membrane of bacteria ^50^. To mimic natural phage release conditions, a simple osmotic shock can induce liposome disruption and phage release (fig. 4B, C), replicating the final step of the lytic cycle.

## Discussion

For half a century, achieving a cell-free SC phage infection cycle seemed an ambitious undertaking, considering the complexity of the infection mechanism and molecular architecture. Yet, using a systematic and quantitative approach, we demonstrated that phage T7 can effectively infect CFE-loaded single-bilayer SCs and produce progeny. Most tailed phages initiate infection by reversibly binding to a primary receptor, often a surface glycan like LPS, followed by irreversible attachment to a secondary receptor, typically a core LPS sugar or an outer membrane protein, which stimulates genome injection. This study demonstrates that LPS alone is a sufficient membrane component for T7 genome internalization into SCs, providing insights into the minimal requirements for phage infection. A distinction between T7 and T5 lies in their structural differences and mechanisms of genome ejection. T5 utilizes FhuA as a docking site but does not use it as a direct genome channel. Instead, its long non-contractile tail crosses the bilayer at the periphery of FhuA, facilitating DNA internalization. In contrast, T7 possesses a short non-contractile tail, preventing it from penetrating the bacterial membrane. Unlike previous liposome studies with T5, which lacked cytoplasmic CFE activity, our system incorporates a CFE reaction inside the liposomes, thereby enabling all the steps of a phage infection process in an SC system.

This work demonstrates the extent to which biological systems can function far from their natural conditions and in far simpler molecular settings. Unlike bacteria, the LPS-coated SCs lack a second lipid bilayer and a cell wall, which in natural conditions provides mechanical resistance to osmotic stress. *E. coli* maintains an internal osmolarity of approximately 300 mOsm greater than its surrounding broth, allowing the difference in osmotic pressure to burst the cells and release phage progeny. In contrast, in the cell-free phage cycle presented in this work, T7 infection proceeds but bursting does not occur spontaneously because of the isotonic conditions (around 600 mOsm inside and outside the SCs). Lysis is achieved by transferring the SCs from isosmotic to hypoosmotic milieus (fig. 4B, C). We found that while LPS enables T7 DNA internalization, it inhibits holin pore formation preventing lysis and release. The efficacy of the phage replication cycle inside SCs is limited by several factors. The low multiplicity of infection (MOI) constrains the number of productive infections, while DNA replication in CFE achieves 2–3-fold amplification^28^ compared to the 200–300-fold observed in bacterial hosts due to finite resources in CFE. Newly synthesized phages often rebind to inactive SCs via persistent LPS receptors, reducing the effective phage titer and hindering further propagation.

Despite substantial challenges, this work offers a tractable bottom-up approach to studying viral infection in a controlled environment, decoupling phage biology from the complexities of living bacteria. This approach allows the screening of individual infection variables, free from the complex effects of bacterial metabolism, growth, and division. It also provides a modular platform for investigating intracellular phage-host interactions, including phage defense mechanisms. As a proof of concept, we showed that T7 infection can be aborted by encapsulating CRISPR-Cas and guide RNAs targeting the phage genome within SCs (fig. S32), illustrating the potential of this system for studying antiviral strategies.

Large integrative projects meant for building complex biological systems from scratch are taking shape around the globe. Building SCs from the ground up has become major research in the USA, in Europe and in Asia^5,51–53^. The bottom-up construction of synthetic viral cycles has been recently discussed. The phage cycle built in this work captures these two emerging trends observed in cell-free constructive biology. As bioengineering capabilities are rapidly developing, safety concerns arise too. Biocontainment solutions must be anticipated and imagined. In that respect, the cell-free phage infection mechanism devised in this work delivers fundamental information for developing one type of biocontainment solution to control SC systems.

## Material

### Synthetic cell experiments

Lipopolysaccharides from *Escherichia coli* F583 (Rd mutant) (L6893-5MG, Sigma), 16:0-18:1 PC (POPC) (850457, Avanti Polar Lipids), and 16:0 PEG5000 PE (880200, Avanti Polar Lipids) were used. Additional materials included 4 mL glass vials with Teflon-lined caps (600460, Avanti), Posi-Click tubes from Thomas Scientific (1138W14), liquid paraffin (128-04375, Wako), Optical-Quality Imaging Plates (384-well, black-walled, clear bottom, A58941, Thermo Scientific), dNTP set (100 mM, 10297018, Thermo Fisher), SYBR Gold Nucleic Acid Gel Stain (S11494, Thermo Fisher), Fluorescein-12-dCTP (NU-809-FAMX-S, Jena Bioscience), Tetramethylrhodamine-Dextran (3000 MW, Anionic, Lysine Fixable, Thermo Fisher), and Fluorescein isothiocyanate-dextran (46945-100MG-F, Sigma).

### Phage spotting and lysate purification

0.22 μm centrifuge filter tubes were from (8160, Costar), v-bottom 96-well plates (701201, NEST), and square petri dishes (688102, Greiner Bio-One). Difco agar (214010, BD), and LB broth (BP1426-500, Fisher). 100 kDa membrane Amicon Ultra-15 centrifugal filter units (UFC910024, Millipore). Centrifuge tubes, culture tubes, and 25 mL reservoirs were generic.

### Phage DNA extraction

DNase I recombinant RNAse-free (M0303S, NEB), RNase A (T3018L, NEB), Proteinase K (P8111S, NEB). SDS (20% wt/vol, AM9820, Sigma), Phenol:Chloroform Alcohol (25:24:1, 15593049, Sigma), 3 M sodium acetate solution (pH 5.2, AM9740, Sigma). Absolute ethanol and chloroform were generic, as available.

### Assembly and PCR

The T7 genome was obtained from Boca Scientific (310025). KOD1 polymerase was sourced from Diagnocin (TYB-KMM-101), and a PCR clean-up kit from Invitrogen (K310001). PCR tubes (8 strips) were generic, as available. 10X TAE buffer was purchased from Invitrogen (AM9869), and a 1kb+ DNA ladder from Invitrogen (10-787-018) and Greenglo (NC1620342, Thomas Scientific). For assembly, exonuclease III (M0206S, NEB), PEG8000 from Sigma (P2139), and Tris-HCl 1M (15568-025, Invitrogen). All buffers were prepared freshly, pH adjusted, autoclaved and filter sterilized. Large bore low binding tips were used when handling phage genomic DNA and liposomes.

## Methods

### Emulsion transfer method for Synthesis of RdLPS SCs (Fig. S2)

The lipid composition to incorporate RdLPS in liposomes is composed of POPC (760.076 g/mol), PEG-PE 5000 (5745.03 g/mol) and RdLPS (considered here 2700 g/mol^54^). The final concentration in the oil should be in the range: POPC: 25 μM – 125 μM, PEG-PE: 5 μM – 35 μM RdLPS: 20 μM – 80 μM. In this work, the composition used was POPC/RdLPS/PEGPE 55/25/20 (molar ratio) at a total lipid concentration of 100 μM. This corresponds to 55 μM of POPC (42 μg/mL), 15 μM PEG-PE (85 μg/mL) and 30 μM of RdLPS (80 μg/mL).

### Lipid stock preparation

Lipids (POPC, PEGPE, RdLPS) were dissolved in chloroform at appropriate concentrations (5 mg/mL for POPC and RdLPS; 10 mg/mL for PEGPE) and stored in airtight glass vials with Teflon-lined caps at -20°C. Lipid stocks are stable for up to one month at -20°C in the dark.

### Lipid-in-Oil Mixture Preparation

In a 7 mL glass vial, lipids are mixed in 200 μL hexadecane. Typically, 10 μL of POPC, 20 μL RdLPS and 12 μL PEG-PE were added and vortexed for 15 s. 1 mL long-chain mineral oil (Wako liquid paraffin) is added, and the mixture is heated for 1-2 h at 70°C, vortexed briefly to ensure homogeneity, and used directly.

### CFE reaction Emulsification

In a microcentrifuge tube, 200 μL of the lipid-in-oil mixture was combined with 10 μL of the CFE reaction containing the desired components. The solution was emulsified by vortexing or rubbing on a rack, producing CFE droplets encapsulated in lipid micelles. The emulsion was transferred onto a new tube containing 250 μL of cationic solution and centrifuged at 5000 x g for 10 min to form liposomes.

### RdLPS SCs Transfer and Phage Infection

Transfer of LPS liposomes: After centrifugation, the oil phase is removed. 10 μL of liposome pellet is transferred in a new 1.5 ml tube containing 90 μL of outer solution.

Phage Addition and Imaging: Prepare 100 μL outer solution to which phage is added. Typically, 10^7^ -10^8^ PFU/mL were added. In a 384-well plate, 90 μL of outer solution with phages was combined with 10 μL of liposomes. The plate was centrifuged at 600 x g for 5 min, followed by fluorescence microscopy imaging. Acquisition was done with an inverted microscope IX81 (Olympus) equipped with a camera and proper sets of filters. A 20x air objective or 40x and 100x oil objectives were used for observation. Image analysis was conducted with ImageJ and custom-made programs on Python.

### Buffer Compositions

– Cationic solution: 270 mM K-glutamate, 25.5 mM Mg-glutamate, 45 mM Tris-HCl, autoclaved and adjusted to pH 8 and filter sterilized.
– Outer solution: 30 mM Maltodextrin, 90 mM K-glutamate, 4 mM Mg-glutamate, 1.5% PEG 8000, 4 mM amino acids, and an energy mix identical to the CFE reaction. The buffer is adjusted to pH 8 and filter sterilized.

### TXTL reaction and buffers

The TXTL reactions used in this work were based on the TXTL toolbox 2.0, as previously described ^55^. The *E. coli* strain BL21-ΔrecBCD Rosetta2, with a recBCD knockout to prevent the degradation of linear DNA^56^, was used for lysate preparation. Briefly, Cells were grown in 2xYT medium supplemented with phosphates, pelleted, washed, and lysed using a cell press. The lysate was centrifuged, and the supernatant was incubated at 37°C for 80 minutes. After a second centrifugation, the supernatant was dialyzed at 4°C for 3 h. Following a final centrifugation step, the lysate was aliquoted and stored at -80°C. The CFE reaction consisted of the cell extract, an energy mix, and amino acids. The reaction buffer included 50 mM Hepes (pH 8), 1.5 mM ATP and GTP, 0.9 mM CTP and UTP, 0.26 mM coenzyme A, 0.33 mM NAD, 0.75 mM cAMP, 0.068 mM folinic acid, 1 mM spermidine, 30 mM 3-phosphoglyceric acid (3-PGA), 1 mM dithiothreitol (DTT), 1.5% PEG8000, and 20-40 mM maltodextrin. Amino acids were added at concentrations between 1.5 mM and 3 mM for each of the 20 amino acids. Magnesium (2–5 mM) and potassium (50-100 mM) concentrations were calibrated for deGFP synthesis and T7 phage production to obtain reproducible batches. Reactions were incubated at 30°C in either 1.5 mL tubes, 96-well plates, or encapsulated within liposomes in 384 well-plates.

### Microscopy data analysis

Microscopy images were acquired using an Olympus IX81 inverted microscope controlled by Metamorph software. Liposomes were imaged in 384-well plates, mounted on a custom-made microplate stage. Timelapse acquisitions across multiple wells were performed simultaneously. Temperature was maintained at 30°C by adjusting the microscope room temperature. Bright-field, green, and red fluorescence channels were recorded using phase contrast, GFP, and Texas Red filter sets. Image processing including segmentation and particle analysis was performed with ImageJ. exported data were plotted using Python scripts.

**Figure 2.A**: Fluorescent images of RdLPS liposome populations encapsulating a dye were acquired at 20x magnification. Liposome segmentation and particle analysis were performed with ImageJ. Radius and volume measurements were used to plot size and volume distributions. The dataset includes 37,663 liposomes analyzed from three replicates. The mean liposome radius was 3.7 μm, with liposomes smaller than 20 μm in radius accounting for over 65% of the total encapsulated CFE reaction volume.

**Figure 3.D**: RdLPS synthetic cells were infected by T7-mC-WT, T7-WT, T7-S* and T7-mC-S* phages, and their red fluorescence intensity was recorded after 13 hours of incubation with a microscope at a 20x magnification. For each condition and each replicate, the maximum fluorescence intensity inside each liposome characterized is measured. For the four conditions, a total of N = 28, 27, 28 and 31 liposomes respectively were characterized from three replicate.

**Figure 3.F**: RdLPS synthetic cells encapsulating FITC-dextran were incubated with a serial dilution of T7-mC-S*. Liposome count was estimated by segmenting the green fluorescence channel using consistent thresholding across all datasets. Infected liposome count was determined by segmenting the red fluorescence channel after 15 h of incubation, using the same calibrated thresholding from a negative control (no phage). The multiplicity of infection (MOI) was calculated as the ratio of PFU added per well to the total liposome count. The infected liposome ratio was determined by dividing the red-channel liposome count by the total green-channel liposome count. Ratios were calculated from four replicate.

**Figure 3.G:** RdLPS SCs either infected by T7-mC-S* phages or encapsulating *mcherry* expression gene cascade were incubated over 15 hours. Red fluorescence intensity was recorded with a microscope at 20x magnification. Using the procedure described in Supplementary text, the radius and concentration of in-situ synthesized mCherry were measured for N = 62 (*mcherry* gene encapsulation) and N = 36 (T7-mC-S* infection) liposomes. Liposomes were randomly selected from three replicate experiments in each condition. The linear fit slopes, intercept and R2 are (−0.76, 21.3, 0.35) for T7-mC-S* infection and (0.23, 22.1, 0.02) for *mcherry* gene encapsulation.

**Figure 3.I**: RdLPS SCs either infected by T7-mC-S* phages or left uninfected had their green fluorescence intensity recorded for 15 hours every 15 minutes with a microscope at a 20x magnification. For each condition and each replicate, the timelapse images were cropped around isolated liposomes so that only one liposome would be present in each cropping series. The maximum intensity of each cropped image was then taken as the measurement of the green fluorescence intensity inside the liposome. For the two conditions, a total of N = 27 and 17 liposomes from three replicates respectively were characterized.

**Figure 3.K**: Cluster count over time was analyzed using the green fluorescence channel. To isolate clusters, images from the timelapse green channel were processed by subtracting duplicates blurred time series (with different Gaussian sigma factors), effectively removing non-cluster fluorescence. The resulting cluster images were converted into masks, and clusters were counted over time using particle analysis. Identical parameters were applied to all conditions. Data represent clusters from 4 replicates. No clusters were detected in the absence of phage. Data were exported as CSV files and plotted using Python.

**Figure 4.A and figure S28**: Kinetic plots were generated in Python. mCherry fluorescence (T7-mC-S* infection) and Sybr Gold fluorescence data (from Figure 3I and Supplementary Fig. S22) were analyzed, along with cluster formation data from Figure 3K. Sigmoidal fits were applied to normalized mean Sybr Gold and mCherry fluorescence, while a Gaussian fit was used for normalized dCTP-FITC cluster counts. Min-max normalization was applied, and individual data points were included in Supplementary Fig. S28.

**Figures 4C and 4D**: In these experiments, liposomes encapsulated rhodamine-dextran (red fluorescence dye) and the T7 genome. Liposome count and radius were obtained through red-channel image segmentation, and liposome volume was calculated assuming spherical geometry. The total encapsulated CFE volume was estimated as the sum of all liposome volumes in a well. PFU production in liposomes was determined by liposome lysis and plating on an *E. coli* B lawn. PFU per nanoliter was estimated by dividing the recovered PFU by the total estimated liposome volume. Data represents 4 replicates for 4.C and triplicates for 4.D.

**Figure 4F:** A red fluorescent dextran dye and a phage holin gene circuit were co-encapsulated in liposomes with or without RdLPS. Red fluorescence intensity was recorded with a microscope at 20x magnification every 5 minutes for three hours. Leak ratios were obtained by superposing initial (average of the first 10 images colorized in red) and final (average of the last ten images colorized green) images. Using the superpositions the liposomes were sorted in two groups red liposomes (leaked liposomes) and orange liposomes (no leak). Ratio was calculated as the number of leaked liposomes (red) on total liposomes (red +orange). A total of N = 207, 144, 55, 102, 353 liposomes were processed in liposomes without RdLPS (no holin, T7, phi11, ICP3 and PRD1) and N = 91, 73, 142, 85, 110 liposomes with RdLPS liposomes (no holin, T7, phi11, ICP3 and PRD1) from two replicate experiments.

## Supporting information

SI_A Synthetic Phage Cycle

## Acknowledgments

The authors would like to thank Aset Khakimzhan and Seth Thompson for their help in the preparation of the CFE system used in this work, Lucienne Letellier, Jerome Bonnet, Peter Voyvodic, Petra Schwille, Damien Baigl and David Bikard for the insightful scientific discussions.

## Funding

National Science Foundation (CBET FMRG 2228971) (AL, PS, DG, SB, VN)

École Doctorale Frontières de l’Innovation en Recherche et Éducation—Programme Bettencourt (AL, ABL)

Bettencourt Schueller Foundation support to Engaged Life Science Department of the Learning Planet Institute (ABL)

Human Frontier Science Program (LT0007/2024-L) (PS)

Fulbright France (AL)

## Author contributions

Conceptualization: AL, ABL, VN

Methodology: AL, DG, PS

Investigation: AL, PS, DG

Visualization: AL, ZI

Funding acquisition: AL, ABL, VN, PS

Project administration: AL, ABL, VN

Supervision: VN, ABL

Writing – original draft: AL, ABL, VN

Writing – review & editing: AL, ABL, VN, SB

## Data and materials availability

All data supporting the findings of this study are available within the article and the supplementary materials. Raw microscopy data are deposited on a public repository.

