## Supplementary material for "A synthetic cell phage cycle": SI_A Synthetic Phage Cycle

**The PDF file includes:**

Supplementary Text
Figs. S1 to S33
Tables S1 to S4
Supplementary references 1-5

**Other Supplementary Materials for this manuscript include the following:**

Movies S1 to S6
Data S1

### Supplementary Text

Determination of in-liposome concentration, interfacial localization and asymmetry of fluorophores from fluorescence microscopy

#### Determination of the concentration of fluorescent molecules in liposomes

We consider a liposome that contains identical fluorescent molecules that can partition unevenly between the bulk and the surface of the liposome. We assume the concentration of those fluorescent molecules uniform in the outer solution and equal to $C_0$. We also assume the liposome to be spherical with a radius $R$, and to have therefore a distribution of the concentration of the fluorescent molecules with a radial symmetry $c(r)$ where $r$ is the distance from the center of the liposome. The fluorescence intensity $I(x, y)$ inside the liposome measured with a microscope at the location $(x, y)$ at the observation plane is the sum of the light emitted by all the points M at the coordinates $(x, y, z_M)$ across the thickness $H$ of the sample collected by the microscope:

$$I(x,y) = \alpha\left[\int_{-h(x,y)}^{h(x,y)} c\left(\sqrt{x^2+y^2+z_M^2}\right)dz_M + (H-2h(x,y))C_0\right] + \eta$$

where $h(x, y)$ is the vertical coordinate of the point at the interface of the liposome at the coordinates $(x, y)$, $\alpha$ is the proportionality coefficient between the fluorescence intensity captured by the microscope and the sum of the concentrations of the fluorescent molecules collected along the light beam, and $\eta$ the electronic noise of the camera. Microscopy images are focused on the equatorial plane of the dextran-rich droplets; therefore, we can express this equality in cylindrical-polar coordinates$(r', z_M)$ where $r' = \sqrt{x^2 + y^2}$:

$$\begin{aligned} I(r') &= \alpha\left[2\int_0^{h(r')} c\left(\sqrt{r'^2+z_M^2}\right)dz_M + \left(H-2h(r')\right)C_0\right] + \eta \\ &= \alpha\left\{2\int_0^{h(r')} \left[c\left(\sqrt{r'^2+z_M^2}\right) - C_0\right]dz_M + C_0H\right\} + \eta \end{aligned}$$

With

$$h(r') = R\sqrt{1-\left(\frac{r'}{R}\right)^2}$$

The fluorescence intensity measured in the outer solution is:

$$I_0 = \alpha C_0 H + \eta$$

The difference of fluorescence intensities measured between the inside and the outside of the liposome is then directly proportional to the difference of concentration of the fluorescent molecule between the inside and the outside of the liposome:

$$I(r') - I_0 = 2\alpha\int_0^{h(r')} \left[c\left(\sqrt{r'^2+z_M^2}\right) - C_0\right]dz_M$$

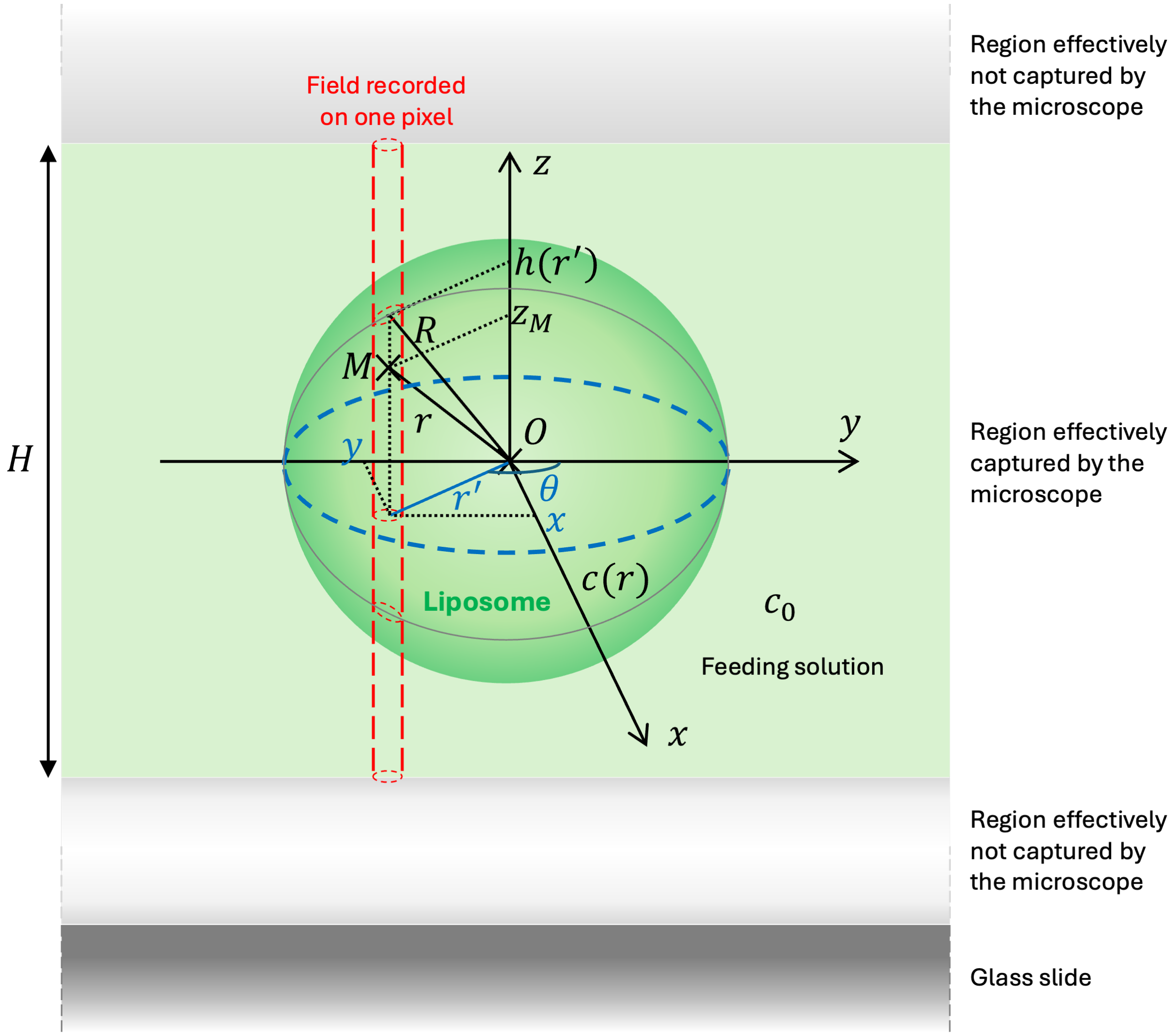


*Parametrization of the fluorescence microscopy imaging of a liposome. The coordinates in the observation plane are written in blue. The gray line runs along the surface of the liposome.*

$I(r') - I_0$ grows with the section of liposome captured by the microscope. To obtain an estimation of the difference of concentration of fluorescent molecules between the inside and the outside of the liposome, that difference of fluorescence intensity must be divided by the height of liposome captured by the microscope:

$$\frac{I(r') - I_0}{2h(r')} = \frac{\alpha}{h(r')} \int_0^{h(r')} \left[ c\left(\sqrt{r'^2 + z_M^2}\right) - C_0 \right] dz_M$$
$$= \alpha \left[ \frac{1}{h(r')} \int_0^{h(r')} c\left(\sqrt{r'^2 + z_M^2}\right) dz_M - C_0 \right]$$
$$= \alpha \left[ < c\left(\sqrt{r'^2 + z_M^2}\right) >_{z_M} - C_0 \right]$$

where $< c\left(\sqrt{r'^2 + z_M^2}\right) >_{z_M}$ is the average concentration of fluorescent molecules in the column of liposome at the coordinates $(x, y)$ and now noted $\tilde{c}(r')$:

$$\tilde{c}(r') =< c\left(\sqrt{r'^2 + z_M^2}\right) >_{z_M} = \frac{1}{h(r')}\int_0^{h(r')} c\left(\sqrt{r'^2 + z_M^2}\right) dz_M$$

We can therefore measure experimentally the quantity $\alpha\tilde{c}(r')$ as:

$$\alpha\tilde{c}(r') = \frac{I(r') - I_0}{2h(r')} + \frac{I_0 - \eta}{H}$$

$\eta$ is measured in the absence of illumination and is found as $\eta = 500$ .
$H$ was measured in a previous study done with the same microscope and using a method similar to the one currently described[1] and was found as $H = 59\ \mu m$ . The value of $H$ is larger than the sizes of all the liposomes screened, therefore the assumption that the liposomes are perceived as spherical under the microscope is justified.

Because:

$$\tilde{c}(r') = \frac{1}{h(r')}\int_0^{h(r')} c\left(\sqrt{r'^2 + z_M^2}\right) dz_M$$
$$= \frac{1}{h(r')}\int_{r'}^{R} c(u)\frac{udu}{\sqrt{u^2 - r'^2}}$$

we cannot extract $c(r)$ from the intensity profile $I(r')$. However, we have

$$\tilde{c}(r') < c(r) >_r$$

where $< c(r) >_r$ is the average of the distribution $c(r)$, different from the average concentration $C$ inside the liposome. Since the interface contains significantly fewer fluorescent molecules than the rest of the liposome, we make the additional assumption that $< c(r) >_r$ is close to the concentration of the fluorescent molecules away from the interface of the liposome, that we assume equal to its value at the center of the liposome. Therefore:

$$\tilde{c}(r') \approx_{r' \sim 0} c(0) \approx C$$

Therefore, we can measure the bulk concentration of the fluorescent molecules inside the liposome up to a multiplicative constant. To increase the accuracy of that estimation, we use the average of $\frac{I(r\prime)-I_0}{2h(r')}$ over the 20% of the radius closer to the center of the liposome while accounting for its spherical symmetry, defined as:

$$\langle \frac{I(r') - I_0}{2h(r')} \rangle_{r' \sim 0} = \frac{\int_0^{0.2R} \frac{I(r') - I_0}{2h(r')} r' dr'}{\int_0^{0.2R} r' dr'}$$

This leads to:

$$\alpha c(0) = \frac{\int_0^{0.2R} \frac{I(r') - I_0}{2h(r')} r' dr'}{\int_0^{0.2R} r' dr'} + \frac{I_0 - \eta}{H}$$

**Determination of the coefficient of interfacial localization of fluorescent molecules at the interface of liposomes.**

We also have:

$$\tilde{c}(r') \approx_{r' \sim R} c(R)$$

Following the same reasoning as previously, we can measure $c(R)$ up to a multiplicative constant experimentally by considering the 20% of the radius closer to the interface of the liposome:

$$\alpha c(R) = \frac{\int_{0.8R}^{R} \frac{I(r') - I_0}{2h(r')} r' dr'}{\int_{0.8R}^{R} r' dr'} + \frac{I_0 - \eta}{H}$$

The coefficient of interfacial localization $\gamma$ of fluorescent molecules near the interface, defined as the ratio of the concentration of fluorescent molecules near the interface over their concentration at the center of the liposome can be obtained directly from fluorescence microscopy imaging as:

$$\gamma = \frac{c(R)}{c(0)} = \frac{\alpha c(R)}{\alpha c(0)} = \frac{\frac{\int_{0.8R}^{R} \frac{I(r') - I_0}{2h(r')} r' dr'}{\int_{0.8R}^{R} r' dr'} + \frac{I_0 - \eta}{H}}{\frac{\int_0^{0.2R} \frac{I(r') - I_0}{2h(r')} r' dr'}{\int_0^{0.2R} r' dr'} + \frac{I_0 - \eta}{H}}$$

A coefficient of interfacial localization of 1 means that the fluorescent molecules are uniformly distributed in the liposome and do not have any particularly favorable interaction with the interface of the liposome. The larger the coefficient, the more fluorescent molecules accumulate at the interface.

**Determination of the coefficient of asymmetry of GFP-fused tail fiber proteins at the interface of liposomes.**

To evaluate the degree of asymmetry of the adsorption of GFP-fused tail fiber proteins (GFP-TF*) at the interface of liposomes, we measured their coefficients of interfacial localizations $\gamma_i$ and $\gamma_{io}$ when GFP-TF* were present only inside or both inside and outside of the liposomes, respectively.
$\gamma_i$ and $\gamma_{io}$ were obtained from the same population of liposomes but incubated in different outer solutions: one devoid of GFP-TF* and one that contains GFP-TF*. Therefore, $\gamma_i$ and $\gamma_{io}$ are obtained with the same concentration $C$ of GFP-TF* inside the liposomes:

$$\gamma_i = \frac{c_i(R)}{C} \; and \; \gamma_{io} = \frac{c_{io}(R)}{C}$$

The excess of GFP-TF* at the interface would simply be $\gamma_i - 1$ and $\gamma_{io} - 1$ respectively. We also assume that at the concentrations of GFP-TF* used inside and outside the liposomes, the interface is saturated in GFP-TF* and that the adsorption of GFP-TF* on either of the sides of the liposome interface does not influence GFP-TF* adsorption on the other side of the liposome interface, therefore,

$$\gamma_{io} = \gamma_i + \frac{c_o(R)}{C} \geq \gamma_i$$

where $c_o(R)$ is the contribution to the interfacial concentration of GFP-TF* coming from the outer solution. For a symmetric adsorption of GFP-TF* at the interface, one would expect twice as much excess GFP-TF* when GFP-TF* are present in the outer solution than when they are absent:

$$symmetric\ interface \rightarrow \gamma_{io} - 1 = 2(\gamma_i - 1)$$

Comparing the excess concentrations of GFP-TF* at the interface, its rescaled degree of asymmetry $\sigma$ is then defined as:

$$\sigma = \frac{(\gamma_{io} - 1) - 2(\gamma_i - 1)}{\gamma_i - 1} = \frac{\gamma_{io} - 2\gamma_i + 1}{\gamma_i - 1}$$

When $\sigma = 0$ the adsorption of GFP-TF* at the interface is the same on either side, while the larger $\sigma$ is, the more favorable the outer surface of the interface is to GFP-TF* adsorption.

Engineering and phenotyping phage T7-mC-S* and T7-Split-S*

**Phage engineering**

Phage assembly was performed following the PHEIGES workflow previously described method[2]. Preparation of the 2X assembly buffer should be done in advance, aliquoted into 50 µL tubes, and stored at -20°C. 2X Assembly Buffer: 20 mM Tris-HCl (pH 7.9), 100 mM NaCl, 20 mM MgCl2, 10% (w/v) PEG 8000, 2 mM Dithiothreitol (DTT) and 1 U/µL Exonuclease III. Genome Assembly: 2X assembly buffer, DNA fragments, and CFE reaction are thawed on ice. Equal volumes of DNA fragments (1-5 nM) are mixed to create an equimolar DNA mix.

T7-mC-S* and T7-Split-S* DNA mixes were prepared following assembly maps in Tables S2-3 at 1 nM for each fragment. 2.5 µL of DNA mix is mixed to 2.5 µL of 2X assembly buffer in a 1.5 mL tube on ice. The mixture is homogenized, spin down and incubated 1 min at room temperature. The tube is then transferred on a heat block prewarmed at 75°C for 1 min, then at 25°C for 5 min. 7.5 µL of 75% CFE reaction is added to 2.5 µL of the assembly mix. The CFE reaction is gently mix, spined down and incubate at 29°C for at least 4 h.

**Phage phenotyping**

CFE reactions containing the synthetized T7-mC-S* or T7-Split-S* were then spotted on an *E. coli rfaC* lawn (RdLPS sensitive tail fiber mutants). Isolated plaques are picked and mixed to a liquid culture of *E. coli rfaC* at OD=0.05. 200uL of this culture containing the phage is incubated at 37C in a shaking plate reader to follow optical density, GFP and mCherry fluorescence over time. The obtained signals from the plate reader are presented below. The data presented shows OD, red and green fluorescence from 4 wells containing a T7-mC-S* isolated plaque, a T7-Split-S* isolated plaque, only the LB medium and the *E. coli rfaC* culture without phage. The plate reader was incubated at 37C, with orbital shaking and signals were recorded every 5 minutes for 6 hours.

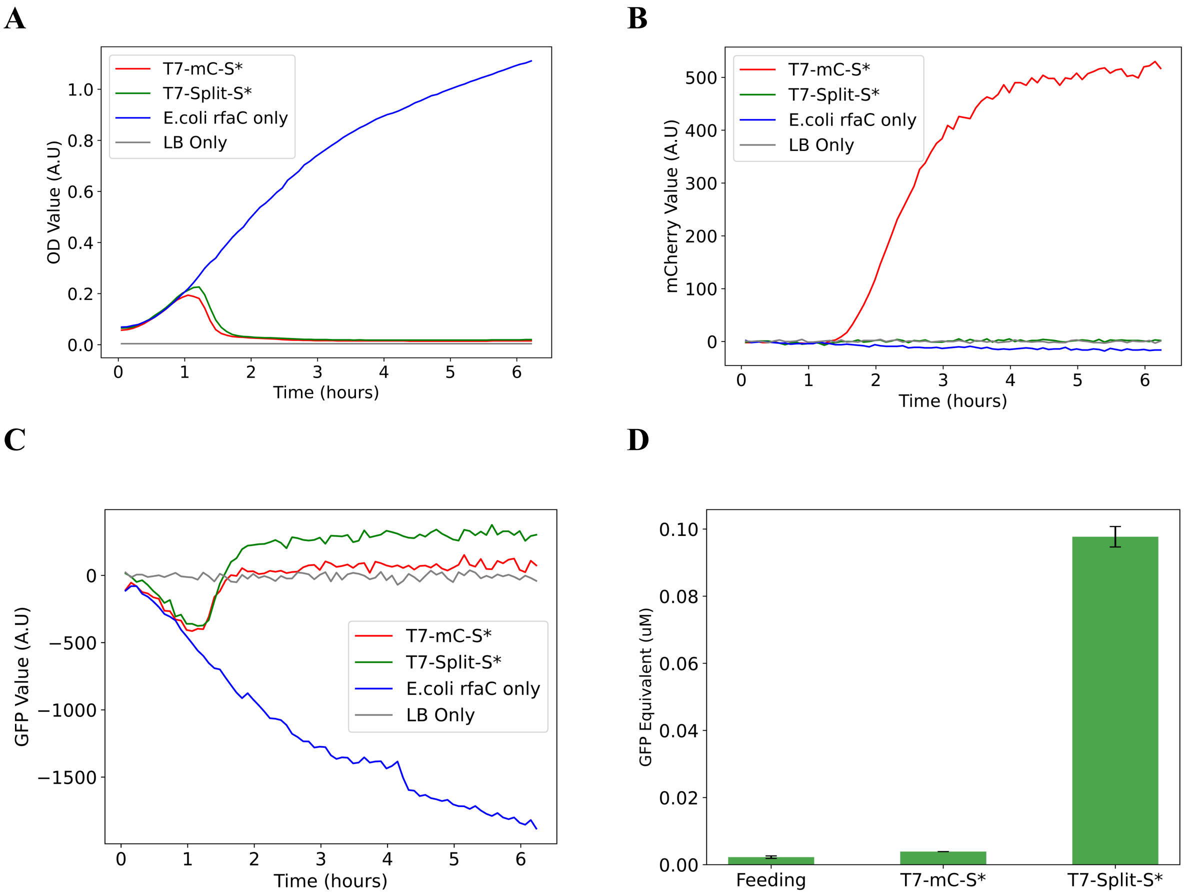


**CFE produced T7-mC-S* and T7-Split-S* Phenotyping. A.** Optical density (OD) kinetics obtained from the 4 wells. In blue bacterial OD increases due to bacterial growth without phage. Red (phage T7-mC-S*) and green (T7-Split-S*) OD drop due to phage induced lysis confirming

RdLPS phenotype. **B.** Red fluorescence signal from the 4 wells over time. Only T7-mC-S* containing well produced a red signal over time confirming successful *mcherry* cassette integration in the phage. **C.** Green fluorescence signal of the 4 wells over time. Data was blanked on the LB signal (medium autofluorescence). The bacterial growth induces a decrease in GFP signal compared to LB. Upon lysis, T7-mC-S* green intensity return to LB level while T7-Split-S* produced a small signal over the LB and T7-mC-S* levels. This indicates production of Split-GFP in the T7-Split-S* containing well. **D.** GFP fluorescence measured in GFP equivalent concentration of concentrated and buffer exchange lysates from T7-mC-S*, T7-Split-S* and the exchange buffer alone. Almost 0.1uM of GFP equivalent are detected in the T7-Split-S* solution indicating successful production of Split-GFP decorated phage particles.

**Phage stock preparation for SCs experiments**

Engineered phage lysates are amplified in 2-5 liter of *E. coli rfaC* as the host until lysis. Phage lysates are treated with 0.5% chloroform to clear remaining bacteria. The lysates are then filter-sterilized through 0.22 µm filters. Lysates are then washed and purified using 100 kDa ultrafiltration columns in phage buffer (10 mM K-glutamate, 1 mM Mg-glutamate, 10 mM Tris-HCl). Phage lysates are washed until no more fluorescence is detected in the pass-through solution (generally 3-5 10mL wash). Phage lysates are recovered in 0.5 mL -1 mL, phage titers are recorded (typically reaching $10^{10}$ - $10^{11}$ PFU/mL) and the phage stock stored at 4°C (phages stocks are used for SCs experiments within 2-3 weeks).

**Phage DNA purification and sequencing**

Phage DNA Purification T7 phage genomic DNA was extracted using the phenol-chloroform method[3]. Briefly, 10 mL of pre-treated lysate ($10^{11}$ PFU/mL) and 10 mL of phenol-chloroform-isoamyl alcohol (PCI) to a separatory funnel. Invert gently, release pressure via the stopcock, and allow layers to separate (20-30 min). Remove the phenol phase (bottom layer) and interphase. Repeat PCI extraction twice more, then perform a final wash with pure chloroform. Recover the aqueous phase, add 1 mL sodium acetate, and 20 mL pre-chilled ethanol. Incubate at -80°C for 1 h, centrifuge at 4000 x g for 20 min, and wash the DNA pellet with 70% ethanol. Resuspend the DNA pellet in 100 µL deionized water (typically 20 nM of T7 phage is obtained). Measure DNA concentration by Nanodrop. Phage genomes were sequenced using long-read nanopore sequencing.

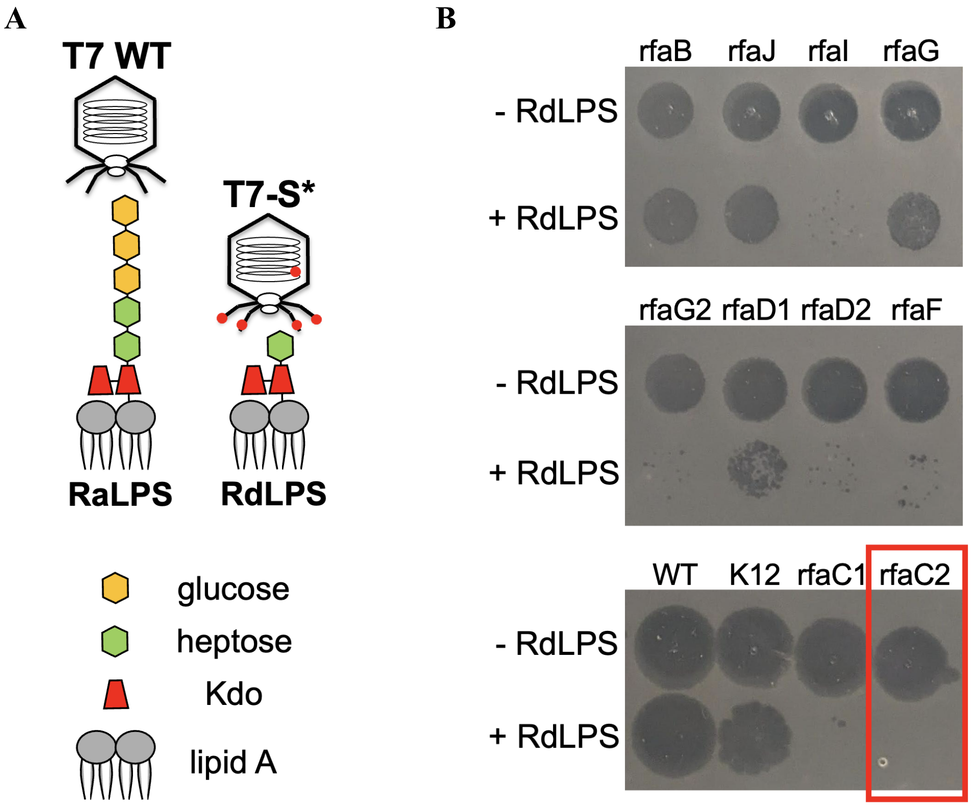


**Fig. S1. A. RaLPS versus RdLPS.** Schematic of T7 wild-type (T7 WT) and T7-S* phages with RaLPS or RdLPS. **A.** T7 WT carries wild-type tail fibers, while T7-S* harbors the S541R mutation in the *gp17* tail fiber gene specific to the RdLPS. **B.** Screening of T7 LPS mutants obtained by tail fiber mutagenesis and selected on *E. coli* strains with different LPS genotypes[2]. Phage lysates ($10^8$ PFU/mL) were incubated with purified RdLPS (100 µg/mL, 3 h at 30°C) or water (control) and spotted. Rd2 LPS (Sigma Cat. No. L6893) from *E. coli* F583 (Rd mutant) contains a KDO-heptose core with a single heptose residue[4]. The T7-S* (phage rfaC2 mutant[2]) was fully inactivated by RdLPS. This mutant tail fiber was used to construct T7-S*, T7-mC-S*, and T7-Split-S* phages in this work.

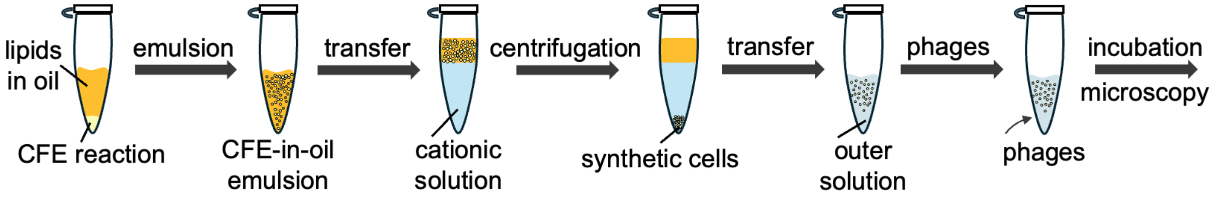


**Fig. S2. Preparation of RdLPS synthetic cells (SCs).** Lipids (POPC/PEGPE/ RdLPS, 55:15:30 molar ratio) were dissolved in mineral oil, heated at 70°C for 1-2 h, and mixed with the CFE reaction to form CFE-in-oil emulsions. The emulsion was layered onto a cationic solution and centrifuged at 5000 × g for 10 min to assemble single-bilayer liposomes. After removing the oil phase, liposomes were transferred to the outer solution with or without phages into a 384 well-plate. Samples were centrifuged at 600 × g for 5 min and imaged on an Olympus IX81 inverted fluorescence microscope.

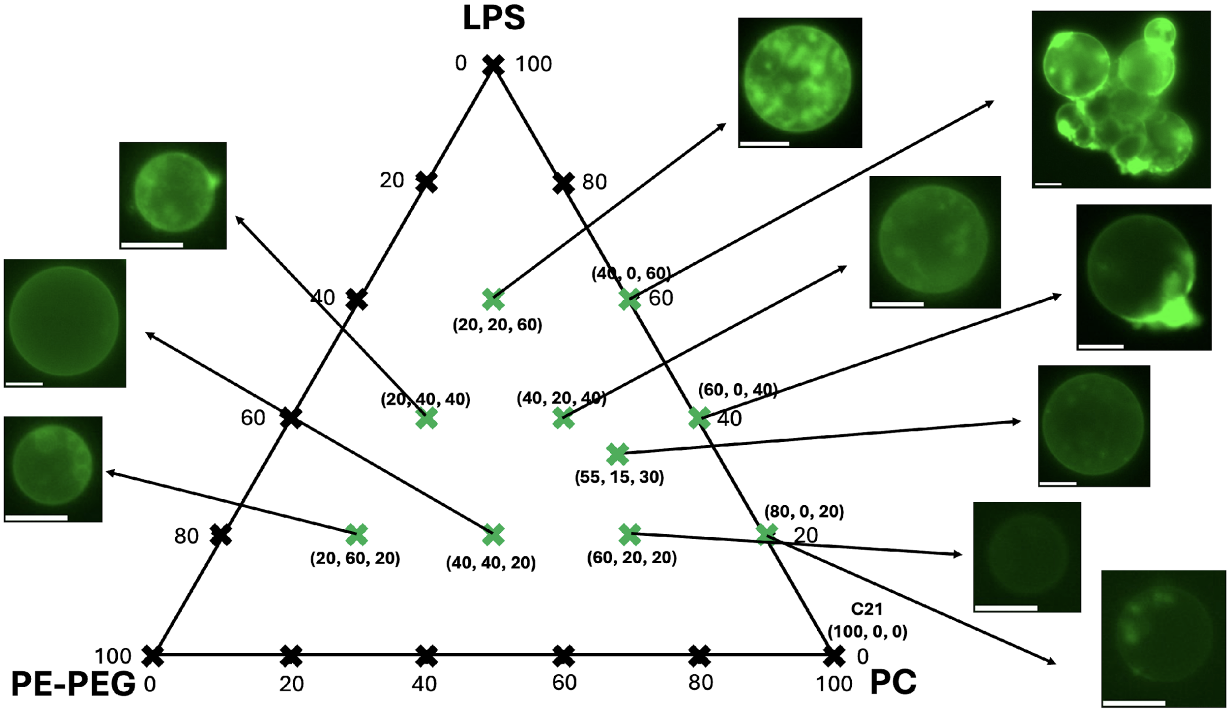


**Fig. S3. Ternary diagram of POPC/PEGPE/RdLPS at a total lipid concentration of 100 µM in oil.** Twenty-two compositions were tested, including 21 combinations in 20% increments for each component and one corresponding to the reported RdLPS content in *E. coli* outer membranes (55:15:30 molar ratio). Green crosses indicate compositions for which liposomes formed, while black crosses indicate no liposome formation, defining a domain of liposome existence. At compositions for which liposomes were formed, GFP-TF* added to the outer solution localized to the membrane, confirming the presence of RdLPS in synthetic cells (SCs). A representative fluorescence microscopy images (green channel) are shown. Reduced RdLPS ratios led to decreased GFP-TF* membrane signals, indicating lower RdLPS incorporation. The absence of PE-PEG resulted in liposome aggregation. Scale bar is 25 µm.

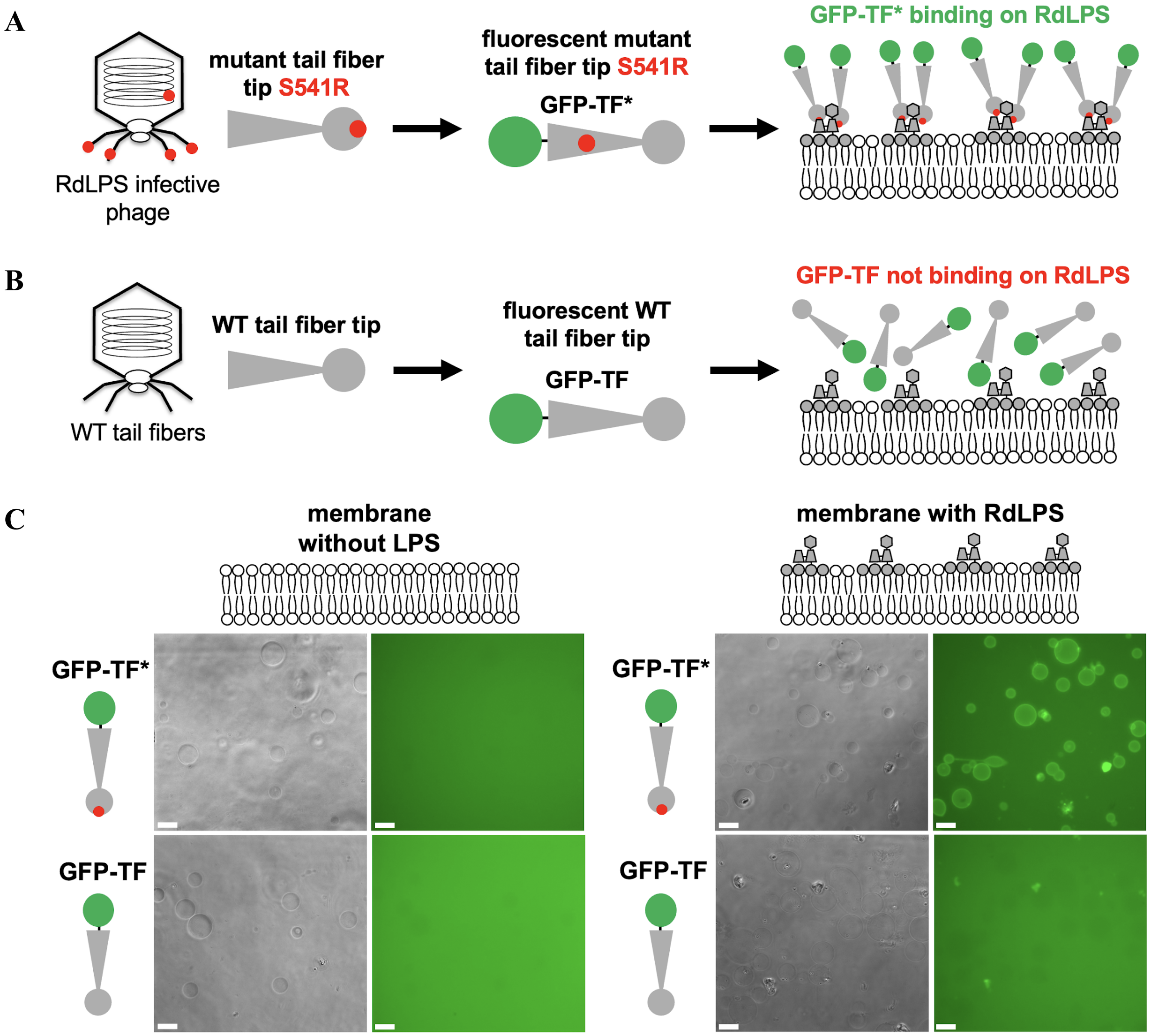


**Fig. S4. RdLPS binding assay. A.** Schematic representation of the design and usage of chimeric fluorescent tail fibers. The tip of the T7-S* tail fiber gene was fused to superfolder GFP (sfGFP) to create a fluorescent LPS-binding construct (GFP-TF*). This construct consists of sfGFP fused to the N-terminal 300 residues of a T7 tail fiber mutant specific for RdLPS. GFP-TF* was cell-free expressed, dialyzed, and added to the outer solution to assess RdLPS presence and activity. Since T7-S* phage is susceptible to RdLPS, GFP-TF* is expected to bind RdLPS, indicating its presence in membranes. **B.** The same method was applied to wild-type T7 tail fibers (GFP-TF). Unlike GFP-TF*, the GFP-TF construct is not expected to bind RdLPS, as wild-type T7 phage does not interact with RdLPS. **C.** Fluorescence localization following the addition of GFP-TF* or GFP-TF to liposomes with or without RdLPS. No fluorescence localization at the surface of the liposomes is observed in the absence of RdLPS. As expected, after 15 min of incubation at room temperature, GFP-TF* shows clear fluorescence localization at the surface of RdLPS-containing liposomes, while GFP-TF does not exhibit any membrane binding. Scale bar is 25 μm.

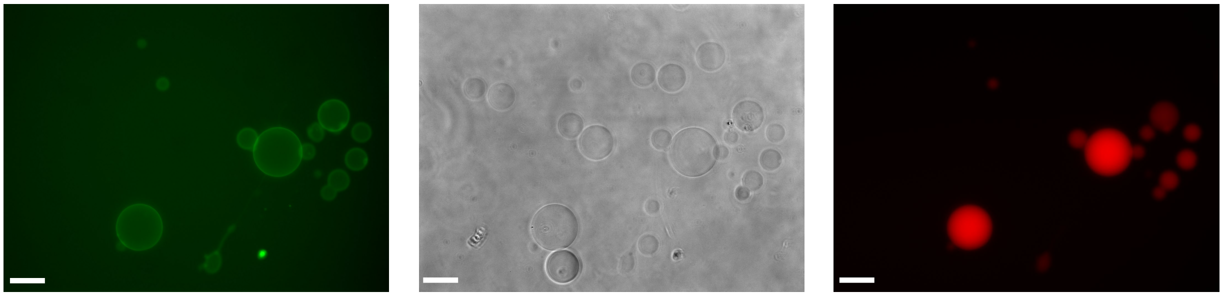

**Fig. S5. Uncropped images on separated channels from Figure 2C.** Picture correspond to a mixture of LPS SCs (red dye) and no LPS liposomes incubated in an outer solution that contains GFP-TF*. The bright field images show the two liposome populations. The green channel corresponds to GFP-TF* fluorescence. Red corresponds to rhodamine-dextran (3 kDa) fluorescence. Pictures were taken after 15 min of incubation; scale bar is 25 µm.

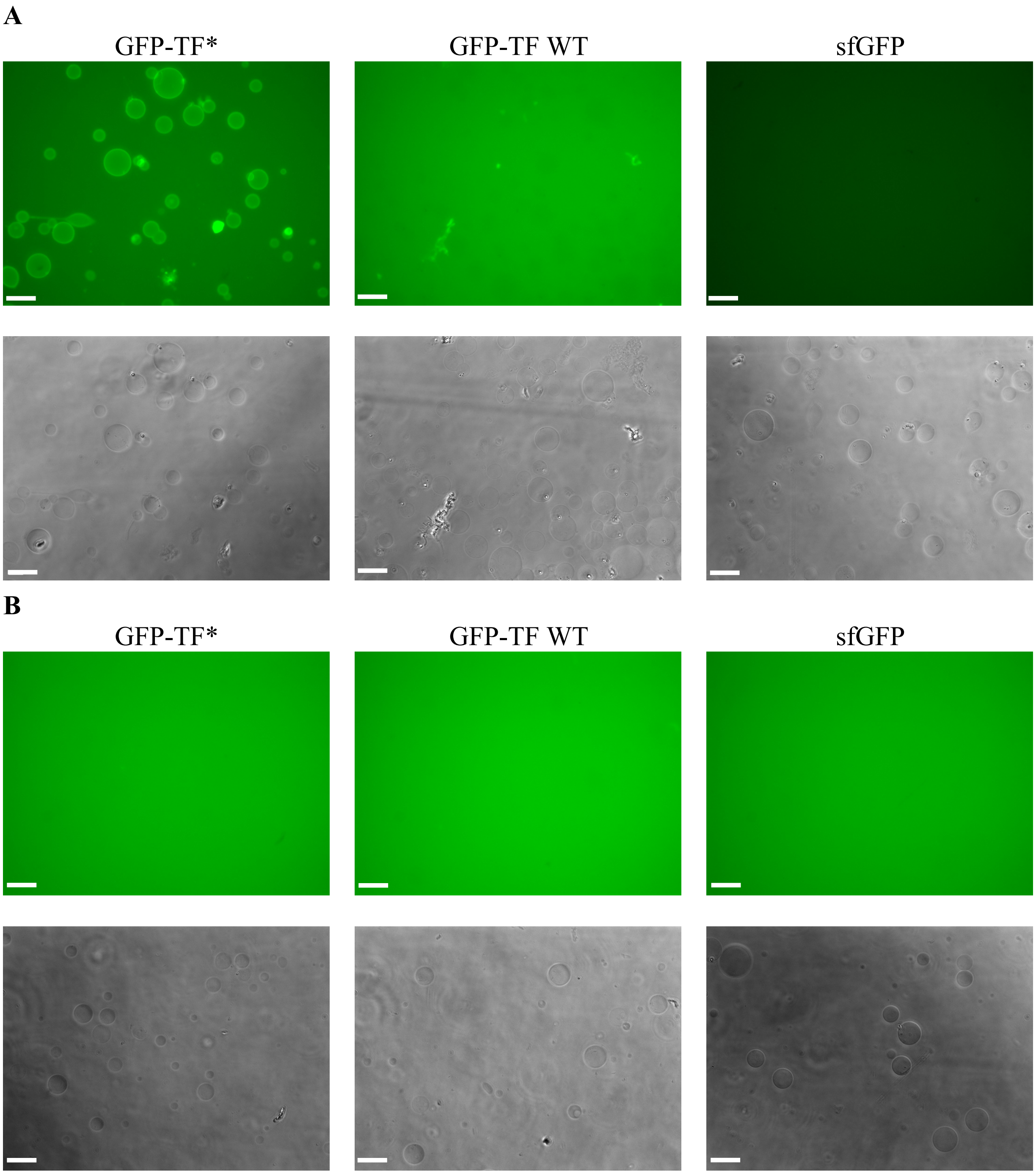


**Fig. S6. Controls GFP-TF*, GFP-TF WT and sfGFP incubated with RdLPS SCs and LPS-free SCs. A.** RdLPS SCs incubated in presence of GFP-TF* (left), GFP-TF WT (middle) or sfGFP (right). As expected, only GFP-TF* bind to RdLPS SCs. **B.** LPS-free SCs are incubated in presence of GFP-TF* (left), GFP-TF WT (middle) or sfGFP. As expected, tail fibers do not localize at the SC membranes. Scale bars are 25 µm, all green channels were acquired with the same microscope settings and are displayed at the same intensity levels.

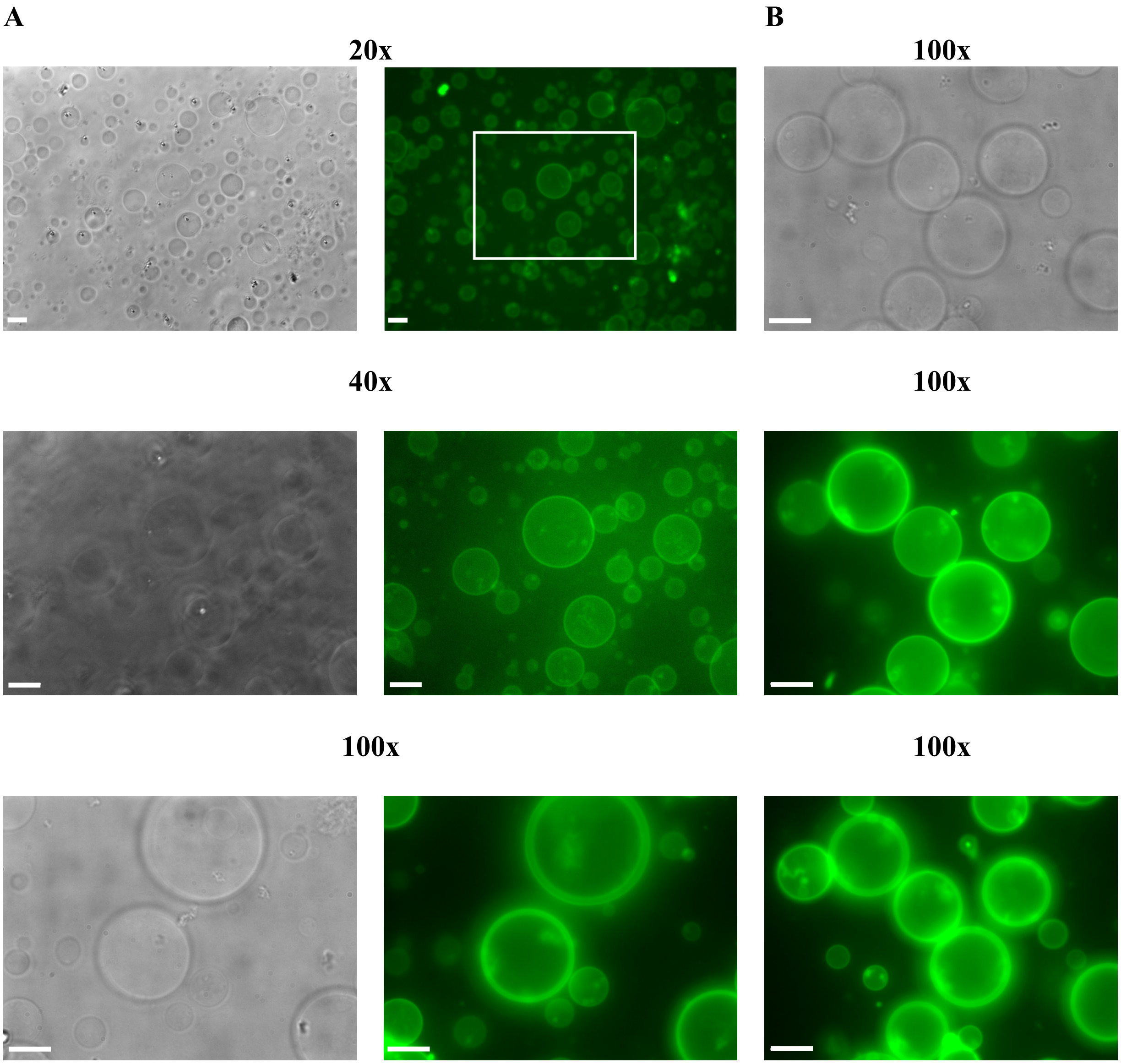


**Fig. S7. RdLPS SCs incubated with GFP-TF*. A.** Zoom in of RdLPS SCs incubated in presence of GFP-TF* added to the outer solution. Images correspond to 20x, 40x and 100x magnification, the white rectangle indicate the magnification area. Scale bars: 20 µm, 20 µm and 10 µm respectively. **B.** 100x magnification of LPS SCs at two different Z positions, focusing on large (top) and smaller (bottom) liposomes of the same viewfield. RdLPS incorporation appears homogeneous. Scale bar: 10 µm.

**A**

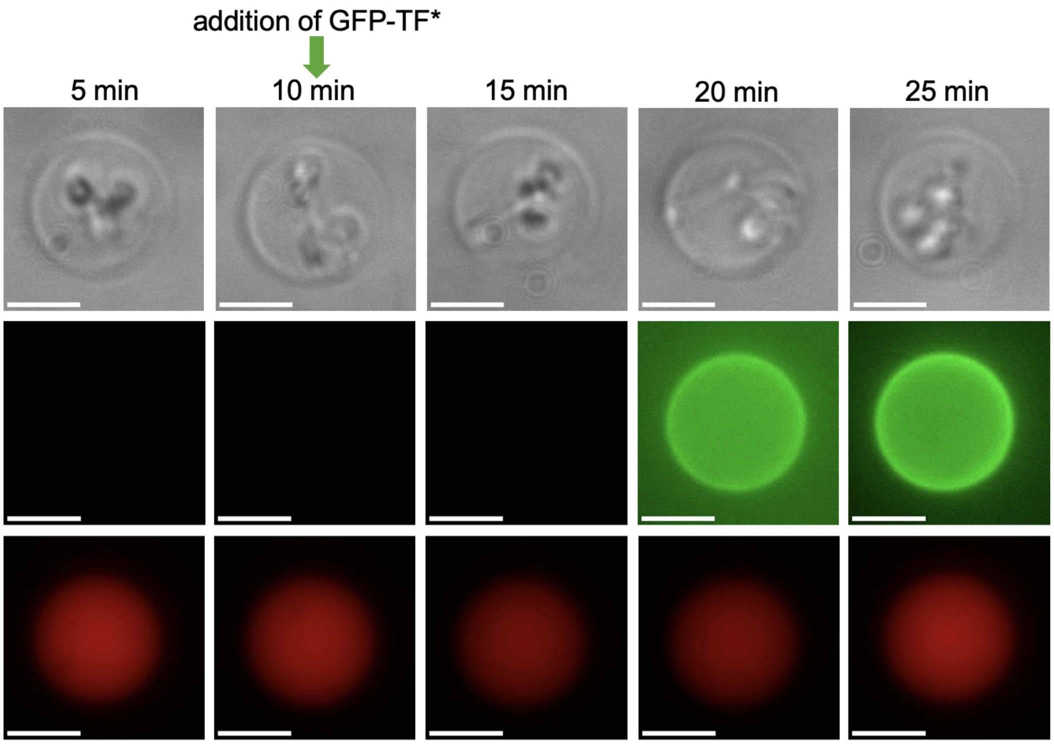


**B**

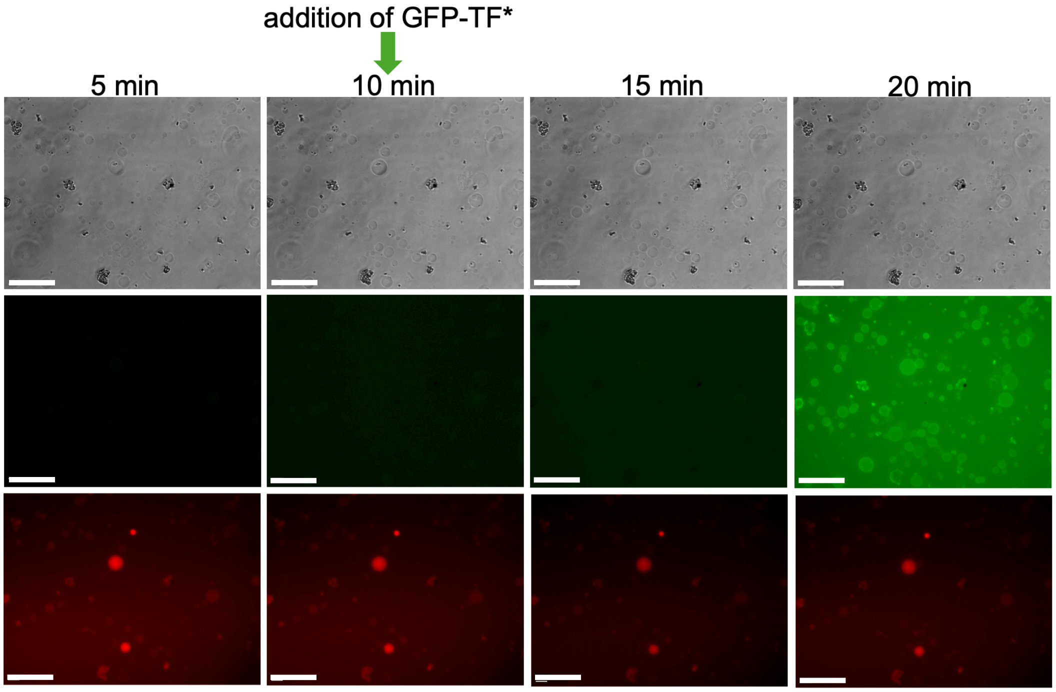


**Fig. S8. Binding kinetics of GFP-TF* to RdLPS SCs.** Time lapse fluomicroscopy of RdLPS SCs following infection by T7-mC-S*. After 15 h of incubation with T7-mC-S*, GFP-TF* is added to the outer solution and bright field, green and red channels are recorded every 5 min. **A.** 100x magnification (scale bar 5 µm) of infected RdLPS SCs. GFP-TFs is added, diffuses and binds to the RdLPS SC in 15 min. **B.** Similar kinetics are observed for 20x magnification. Only a fraction of RdLPS SCs binding GFP-TF* appear red, due to SCs leak, disruption or inactive SCs. This may cause loss of produced virion phages due to rebinding. Scale bar is 50 µm.

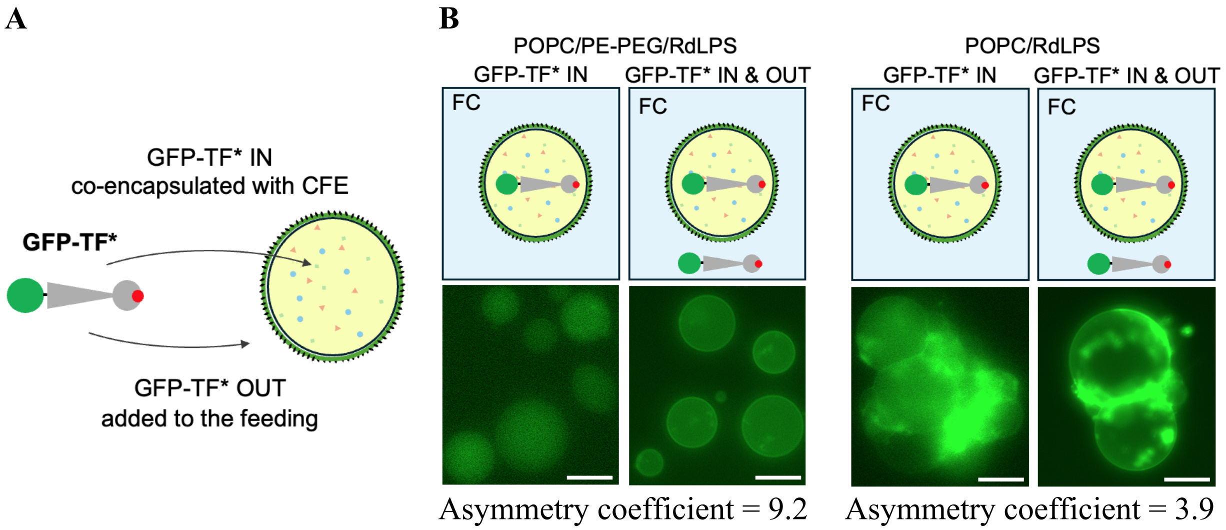


**Fig. S9. Asymmetry of RdLPS distribution in synthetic cells (SCs). A.** Schematic representation of the method used to assess membrane asymmetry of RdLPS. GFP-TF* was either encapsulated with the CFE reaction during the first step of the emulsion transfer protocol ('IN') or added to the outer solution ('OUT'). **B.** Different outer solutions were tested to optimize GFP-TF* binding from the outer leaflet while minimizing internal binding. We quantified GFP-TF* fluorescence localization in RdLPS SCs (POPC/PEGPE/RdLPS, 55:15:30 molar ratio) and SCs without RdLPS (POPC/PE-PEG, 70:30 molar ratio). An asymmetry coefficient was calculated by comparing fluorescence intensity between the inner and outer leaflets (see Supplementary text). The greatest asymmetry coefficient was observed when the cationic solution to outer solution transfer method was used (Fig. S2), indicating enhanced RdLPS localization on the outer leaflet. This suggests that the cationic solution stabilizes RdLPS at the oil-water interface during emulsion formation, promoting its preferential orientation on the outer leaflet. The PE-PEG reduces aggregation between SCs. Scale bar is 10 µm.

**A**

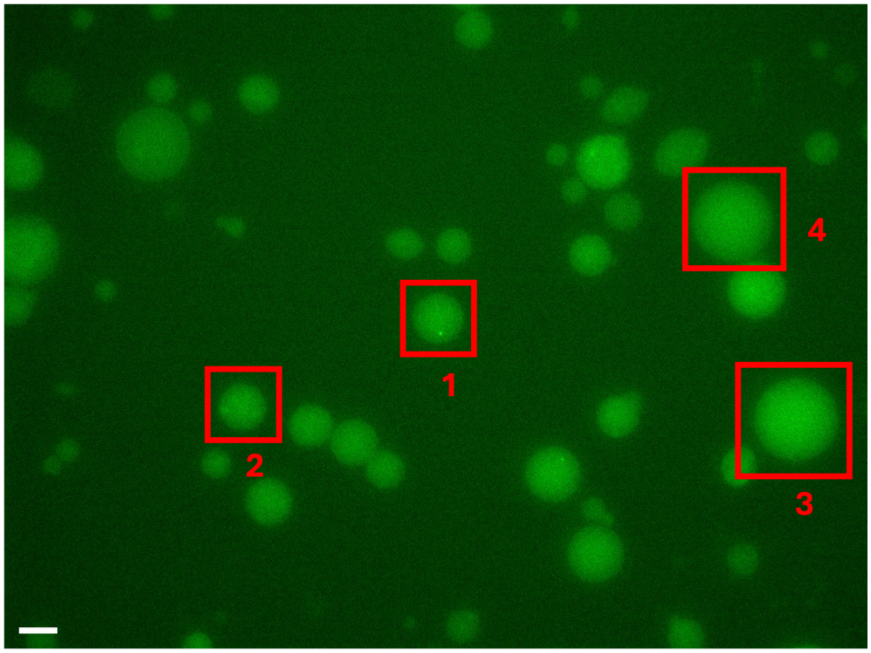

1
2
3
4


**B**

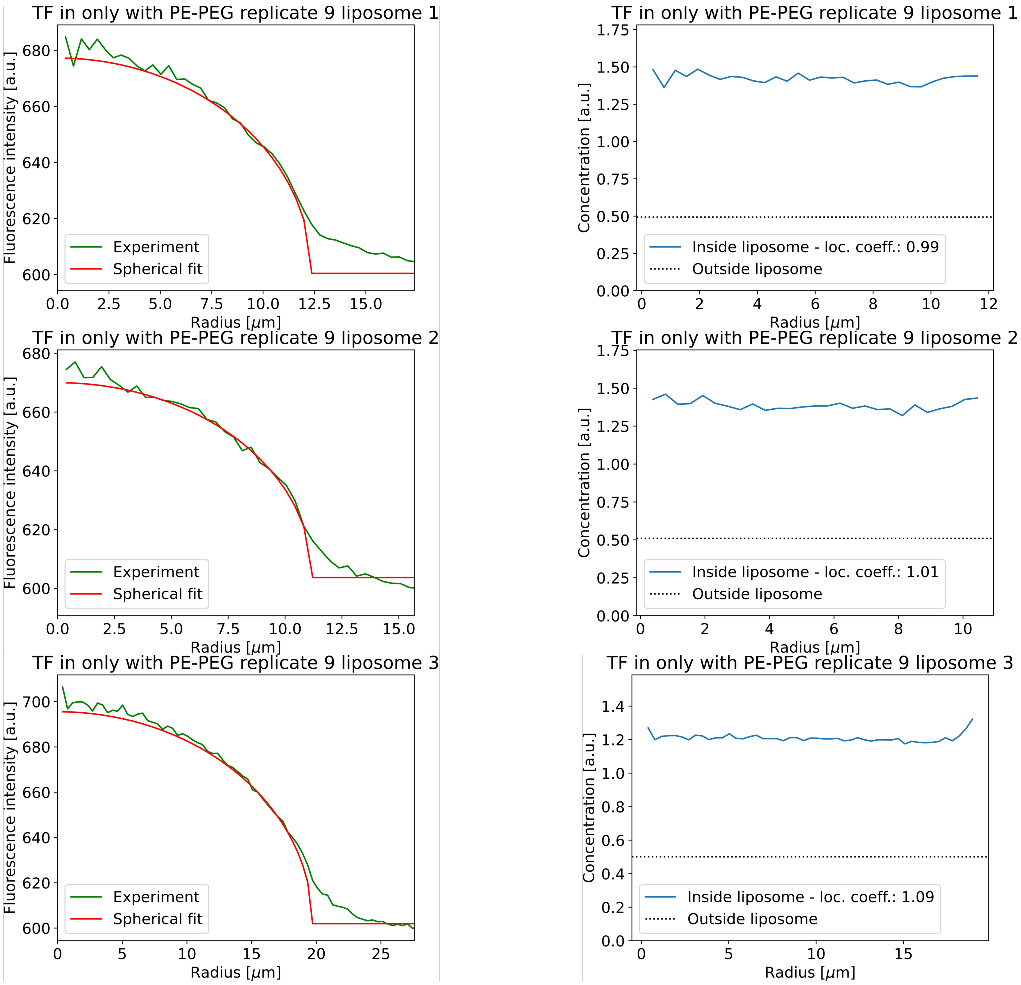

TF in only with PE-PEG replicate 9 liposome 1
Fluorescence intensity [a.u.]
Radius [μm]
Experiment
Spherical fit
TF in only with PE-PEG replicate 9 liposome 1
Concentration [a.u.]
Radius [μm]
Inside liposome - loc. coeff.: 0.99
Outside liposome
TF in only with PE-PEG replicate 9 liposome 2
Fluorescence intensity [a.u.]
Radius [μm]
Experiment
Spherical fit
TF in only with PE-PEG replicate 9 liposome 2
Concentration [a.u.]
Radius [μm]
Inside liposome - loc. coeff.: 1.01
Outside liposome
TF in only with PE-PEG replicate 9 liposome 3
Fluorescence intensity [a.u.]
Radius [μm]
Experiment
Spherical fit
TF in only with PE-PEG replicate 9 liposome 3
Concentration [a.u.]
Radius [μm]
Inside liposome - loc. coeff.: 1.09
Outside liposome

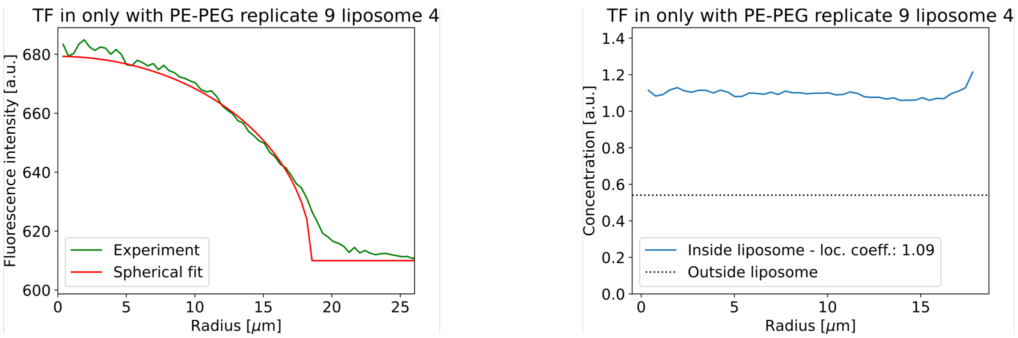


**Fig. S10. Interfacial localization of GFP-TF* on RdLPS SCs with GFP-TF* present only inside liposomes. A.** Fluorescence image corresponding to GFP-TF* fluorescence (green). Four liposomes are presented. A total of N = 47 liposomes from three replicates experiments were processed to calculate the asymmetry coefficient in Fig. S9. **B.** The radial profiles of their green fluorescence intensity (B left, green line) are plotted together with the corresponding best spherical fits (B left, red line) representing the radial profile of green fluorescence intensity of liposomes with the same radius and the same bulk concentration of GFP-TF*. The corresponding radial profiles of the concentration of GFP-TF* are plotted as well (B right). The peak indicates the localization of GFP-TF* at the interface of the liposomes. Scale bar represents 20 μm.

**A**

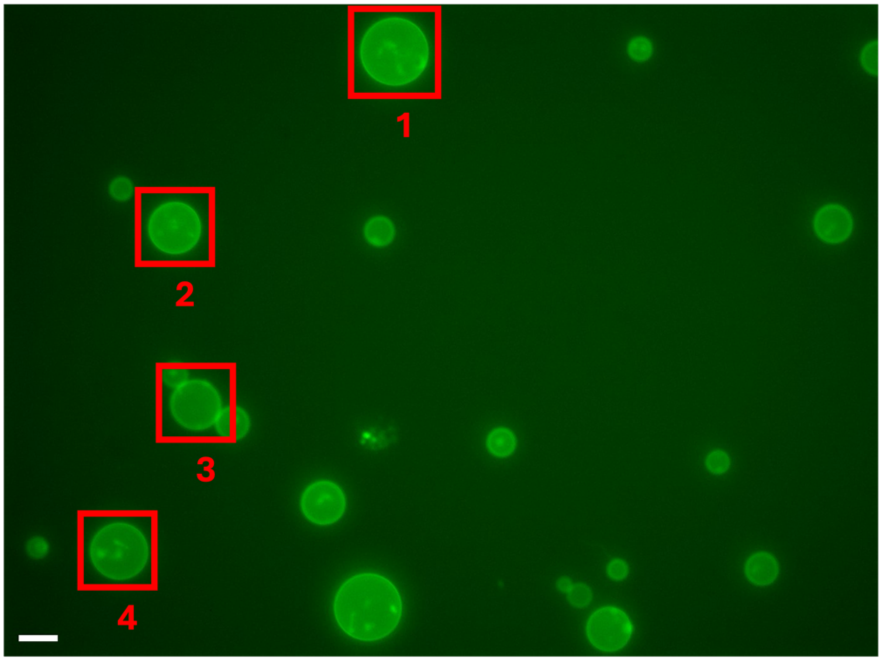
1
2
3
4

**B**

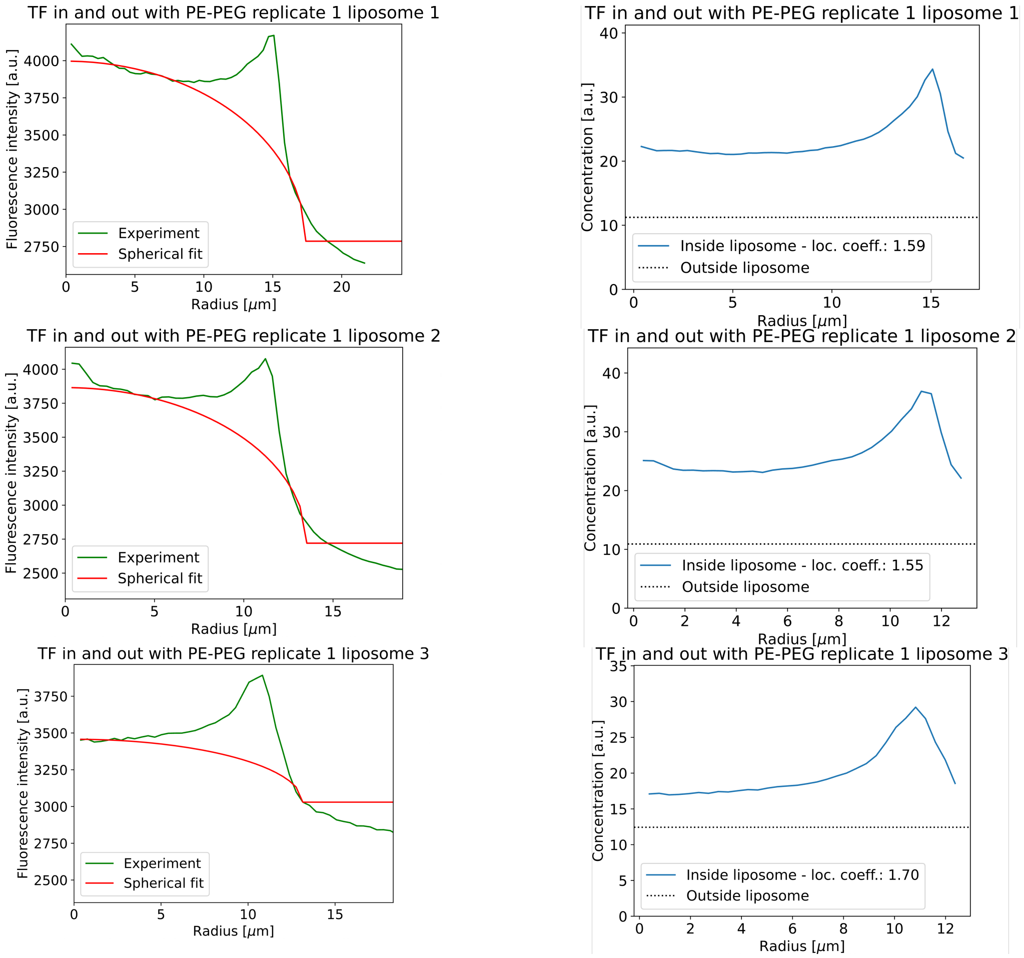
TF in and out with PE-PEG replicate 1 liposome 1
Fluorescence intensity [a.u.]
Radius [$\mu$m]
Experiment
Spherical fit
TF in and out with PE-PEG replicate 1 liposome 1
Concentration [a.u.]
Radius [$\mu$m]
Inside liposome - loc. coeff.: 1.59
Outside liposome
TF in and out with PE-PEG replicate 1 liposome 2
Fluorescence intensity [a.u.]
Radius [$\mu$m]
Experiment
Spherical fit
TF in and out with PE-PEG replicate 1 liposome 2
Concentration [a.u.]
Radius [$\mu$m]
Inside liposome - loc. coeff.: 1.55
Outside liposome
TF in and out with PE-PEG replicate 1 liposome 3
Fluorescence intensity [a.u.]
Radius [$\mu$m]
Experiment
Spherical fit
TF in and out with PE-PEG replicate 1 liposome 3
Concentration [a.u.]
Radius [$\mu$m]
Inside liposome - loc. coeff.: 1.70
Outside liposome

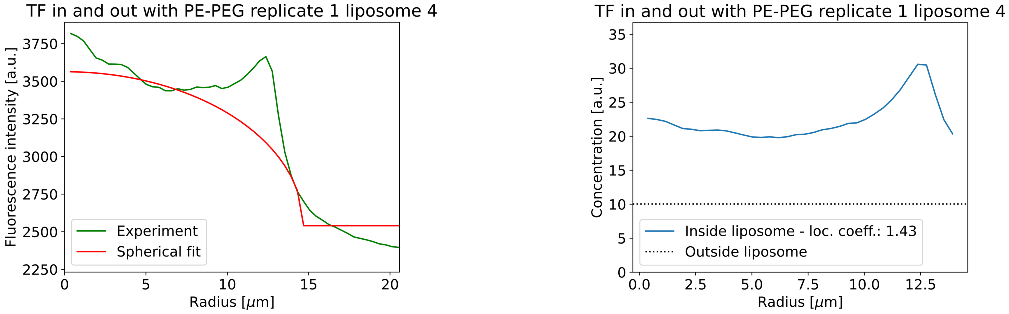


**Fig. S11. Interfacial localization of GFP-TF* on RdLPS SCs with GFP-TF* present both inside and outside the liposomes. A.** Fluorescence image corresponding to GFP-TF* fluorescence (green). Four liposomes characterization are represented. A total of N = 40 liposomes from three replicates experiments were processed to calculate the asymmetry coefficient in Fig. S9. **B.** The radial profiles of their green fluorescence intensity (B left, green line) are plotted together with the corresponding best spherical fits (B left, red line) representing the radial profile of green fluorescence intensity of liposomes with the same radius and the same bulk concentration of GFP-TF*. The corresponding radial profiles of the concentration of GFP-TF* are plotted as well (B right). The peak indicates the localization of GFP-TF* at the interface of the liposomes. Scale bar represents 20 μm.

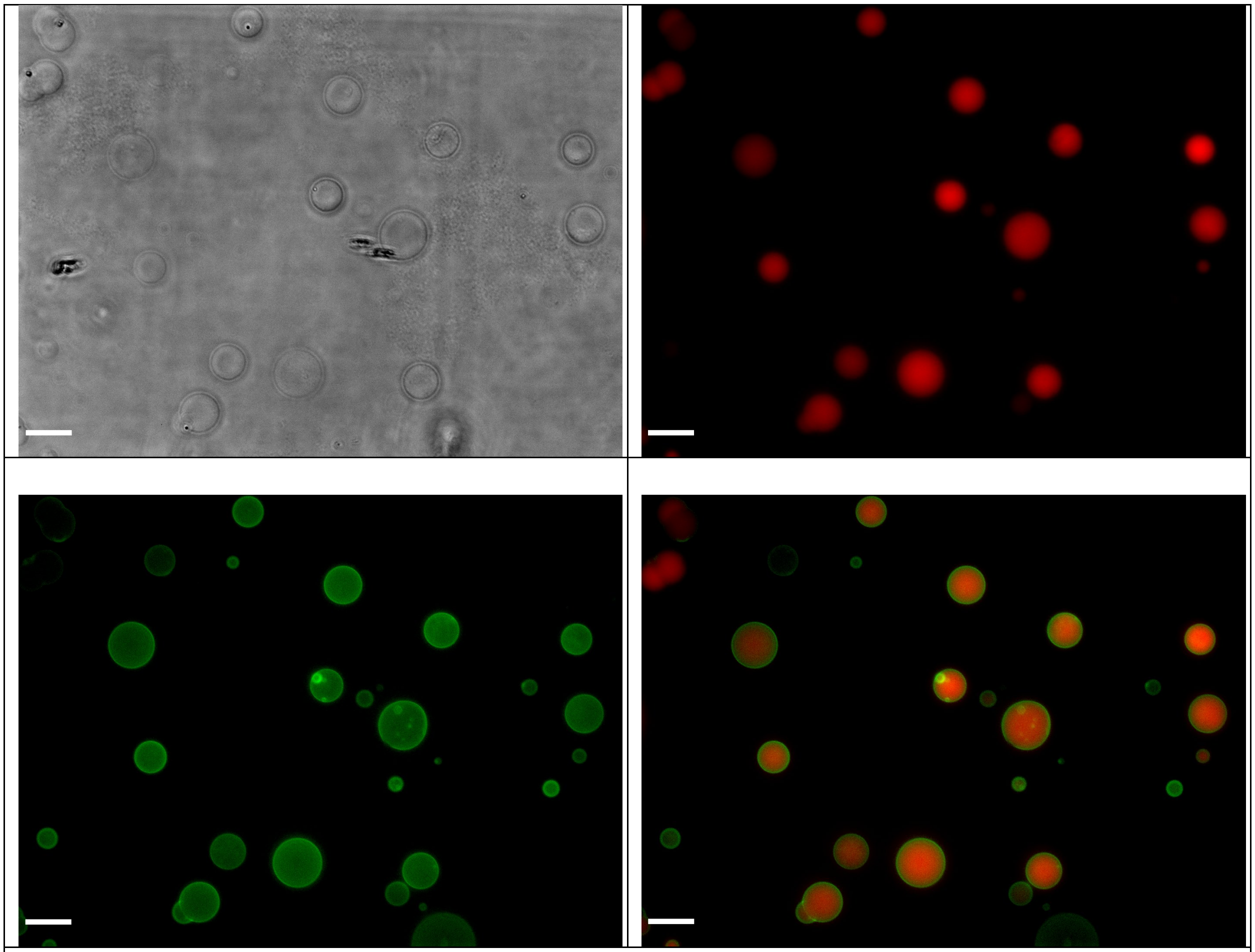

**Fig. S12. Uncropped images on separated channels from Figure 2D.** Images correspond to a mixture of LPS SCs that have expressed *mcherry* and incubated in an outer solution with GFP-TF*. Red signal corresponds to mCherry fluorescence and green signal to GFP-TF*. Images are taken after 15 h of incubation, scale bar is 30 μm.

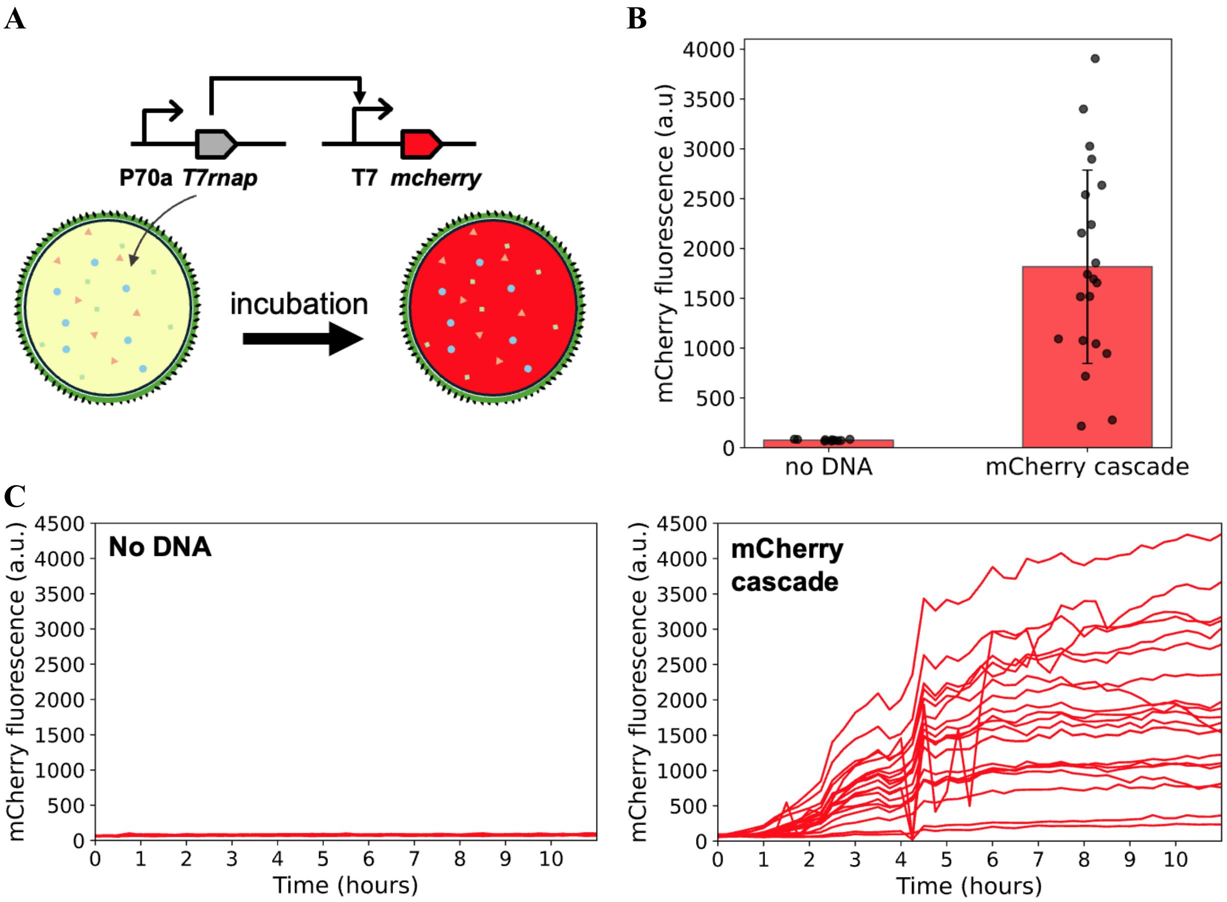


**Fig. S13. Expression of the *mcherry* cascade in RdLPS synthetic cells (SCs). A.** Schematic representation of asymmetric RdLPS SCs encapsulating a *mcherry* expression cascade. The cascade consists of the plasmid P70a-*T7rnap* (constitutive expression of the T7 RNA polymerase) and the plasmid T7-*mcherry* (expression of *mcherry* under a T7 promoter). mCherry fluorescence serves as a readout for active CFE reactions. **B.** Quantification of mCherry fluorescence in RdLPS SCs after 8 h of incubation. Data represent the mean mCherry concentration ± standard deviation from N = 14, 3, and 4 liposomes across three independent experiments for the positive control (*mcherry* cascade) and N = 6 and 4 liposomes for the negative control (no DNA). **C.** Kinetic of *mcherry* expression in RdLPS SCs. Fluorescence intensity was measured every 15 min for both the negative control (left) and *mcherry* cascade (right). Fluorescence drops observed across all liposomes at certain time points result from intermittent microscope lamp intensity drop.

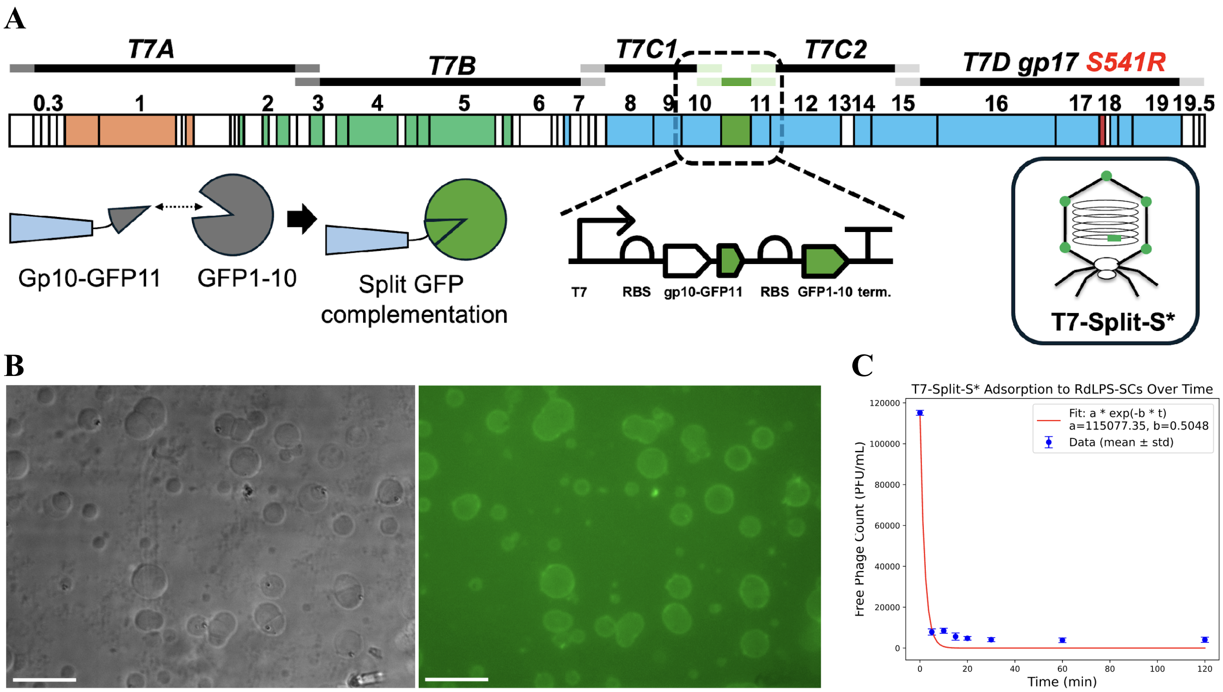


**Fig. S14. Engineering a green fluorescent T7 phage. A.** Schematic of the T7-Split system. Using PHEIGES, a T7 phage with RdLPS-specific tail fibers and fluorescent capsid proteins was assembled. We replaced both versions of the major coat protein gp10A and gp10B in the T7 genome with gp10B fused at the C-terminus to a GFP11 tag (Supplementary text, table S3). The complementary GFP1-10 split GFP fragment was also inserted into the genome, in frame with its own ribosome binding site. This engineered phage, decorated with split GFP, was named T7-Split-S*. The split-GFP system[5] was used to maintain phage capsid self-assembly, as fusion of the full-length sfGFP to gp10B did not produce viable phage when using PHEIGES. **B.** Fluorescence microscopy image of RdLPS SCs incubated for 1 h at 30°C with T7-Split-S* in the outer solution at a titer of $10^{11}$ PFU/mL. T7-Split-S* adsorption led to vesicle deformation and aggregation at this titer, while smaller concentrations produced no detectable signal at the membrane. The green fluorescence observed at the SC membranes corresponds to T7-Split-S* phage adsorption on the RdLPS-rich layer. Scale bar is 30um. **C.** Adsorption kinetics of T7-Split-S* to RdLPS SCs. For this, RdLPS SCs were prepared and diluted to six replicates to 1 x $10^5$ liposomes per tube. T7-Split-S* was added to the SCs at 1.2 x $10^5$ PFU/mL and phage titers in the supernatant were recorded over time. Adsorption kinetics was fitted to exponential decay and adsorption constant, defined as the ratio between the exponential fit factor and the initial liposome concentration (1 x $10^6$ liposome/mL) was determined to be 5 x $10^{-6}$ mL/(min x CFU).

0 h

4 h

8 h

10 h

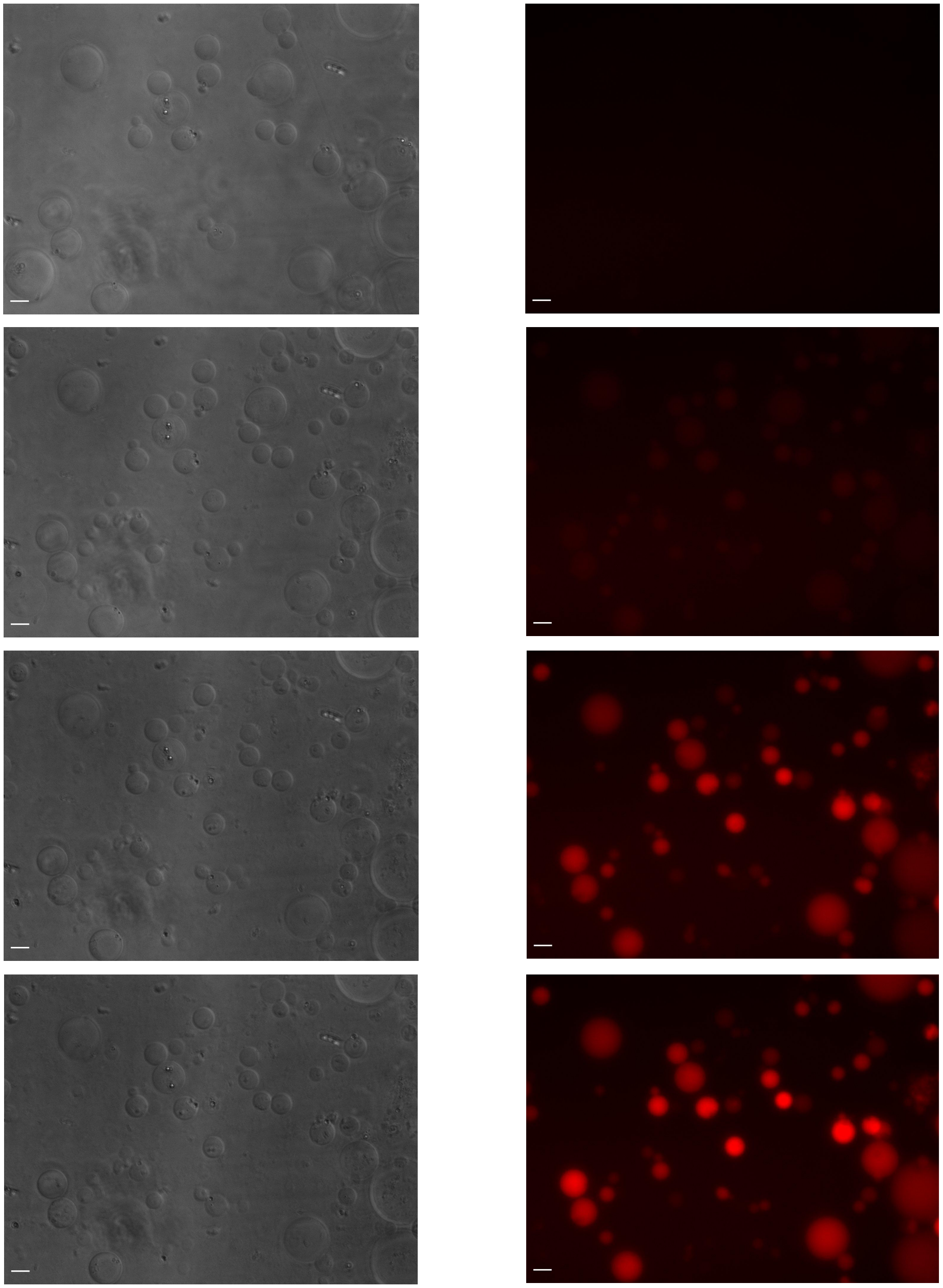

**Fig. S15. Uncropped microscopy images corresponding to Figure 3B, showing separate channels for bright field and red fluorescence.** The red signal represents T7-mC-S* fluorescence. All channels were acquired using identical microscope settings and displayed at the same intensity levels. Scale bar: 20 µm.

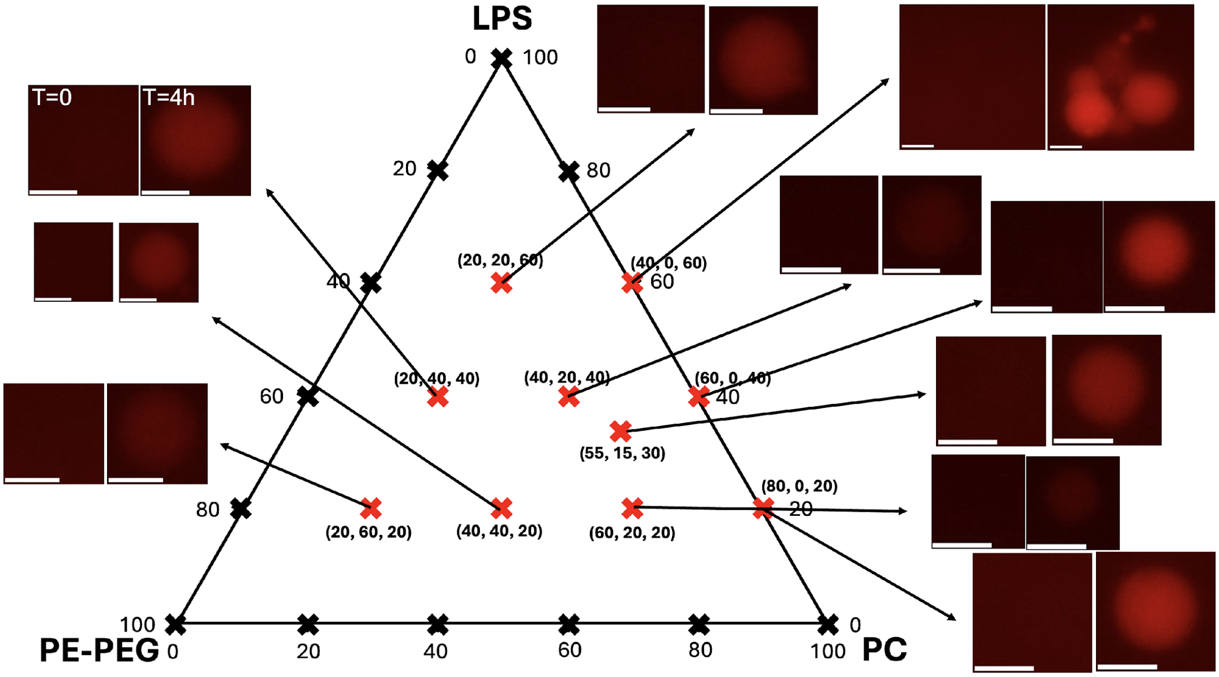


**Fig. S16. Ternary diagram RdLPS SCs infection by T7-mC-S*.** Ternary diagram of POPC/PEGPE/RdLPS at a total lipid concentration of 100 µM in oil for the same compositions as in Fig. S3. Compositions for which liposomes were obtained, T7-mC-S* was added to the outer solution (1x10$^{7}$ PFU/mL). A representative liposome image (red channel) is shown at t = 0 h and after 4 h of incubation. Red fluorescence is observed for all compositions indicating successful liposome infection and *mcherry* expression. Scale bar is 25 µm.

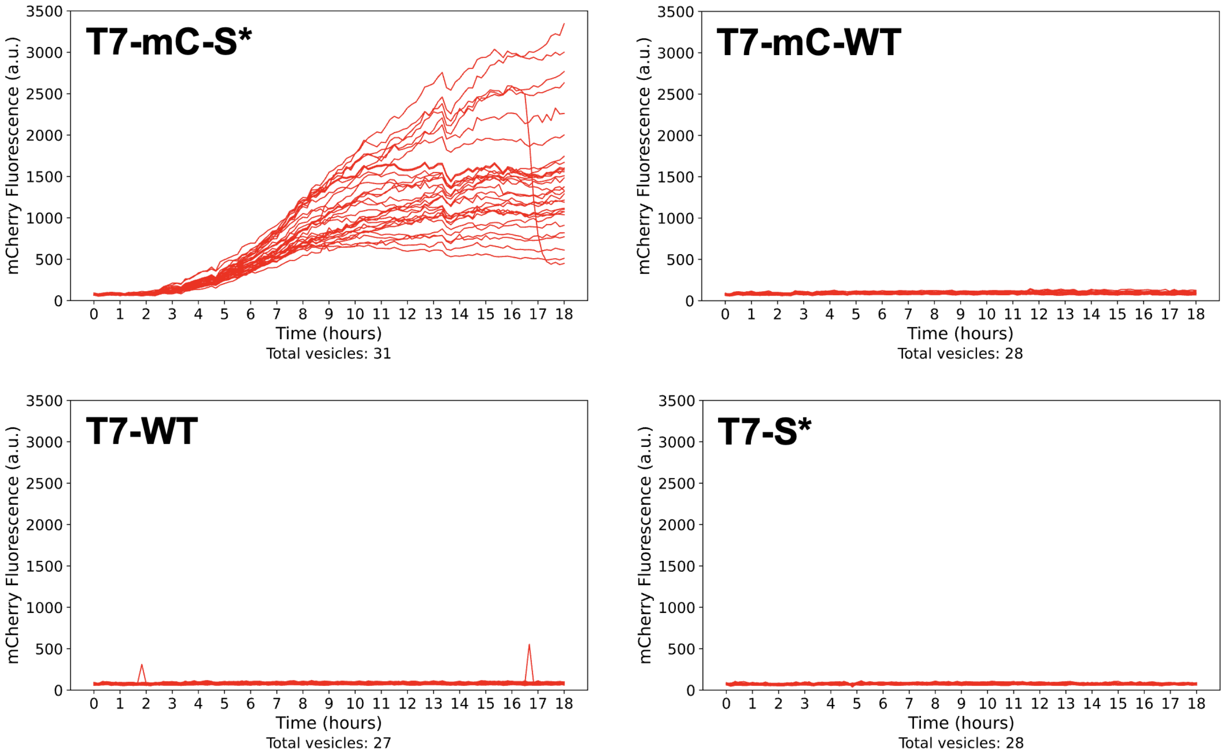


**Fig. S17. mCherry synthesis upon infection of RdLPS SCs by T7-mC-S*, T7-mC-WT, T7-WT, T7-S*.** Red fluorescence kinetics were recorded in single liposomes. Fluorescence intensity was measured every 15 min. Timepoint 640 min was removed due to lamp glitch at this timepoint. Infection by T7-mC-S* corresponds to N = 11, 10, 10 liposomes in replicates 1, 2 and 3, N = 8, 10, 10 liposomes for T7-mC-WT, N = 10, 8, 9 liposomes for T7-WT and N = 10, 8, and 10 liposomes for T7-S*.

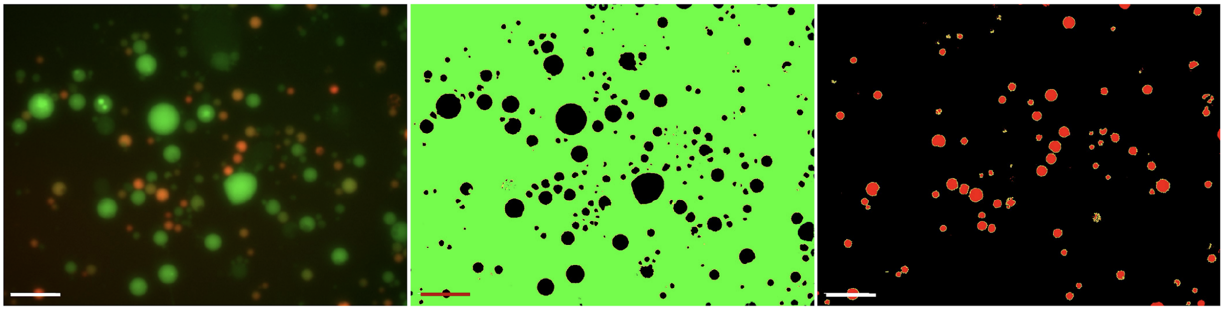

merged image
green: dextran-FITC
red: mCherry from phage infection

Liposomes in viewfield: 243
$N_{tot}$ = 243x12 = 2916
Total PFU in well = 1x10$^5$ PFU.

Infected liposomes: 91

**Fig. S18. Quantification method of the MOI and infected SCs ratio in Figures 3E and 3F.** A green dextran-FITC dye was encapsulated into the SCs to count the total liposomes count per replicate. The MOI was estimated by dividing the PFU/mL added to the outer solution by the total SCs count (green channel). Infected SCs produced mCherry. Red SCs count enabled to estimate the infected SCs ratio. A microscopy image (left) represents the merge green and red channels for a replicate experiment at 1x10$^7$ PFU/mL. Thresholded green channel (middle), MOI is estimated to (1x10$^5$/2916) = 34. Thresholded red channel, infected SCs ratio is estimated to be = 91/243 = 0.37. Scale bar is 50um.

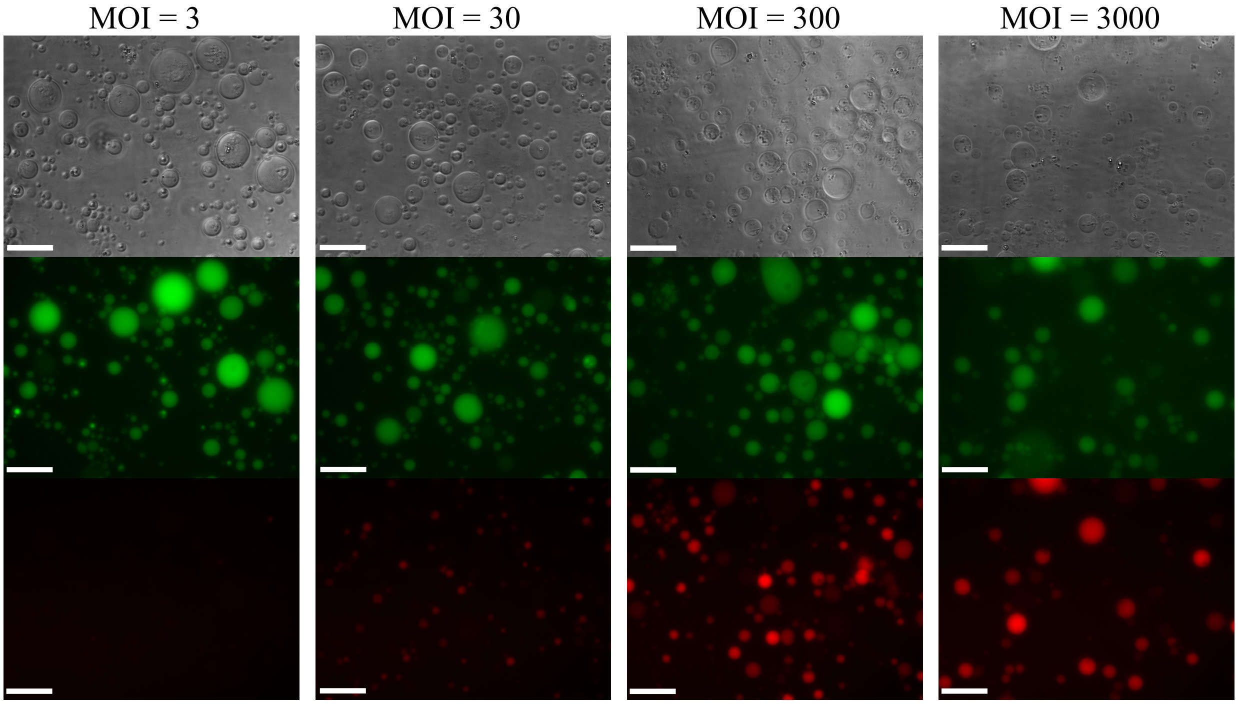


**Fig. S19. Uncropped microscopy images corresponding to Figure 3E.** The red signal represents T7-mC-S* fluorescence, while the green channel shows FITC-dextran (3 kDa) encapsulated within RdLPS SCs. Images were acquired 15 h after adding T7-mC-S* to the outer solution at different concentrations. The PFU/mL added to the well and the estimated MOI are indicated above each condition. All channels were acquired using identical microscope settings and displayed at the same intensity levels. Scale bar: 40 µm.

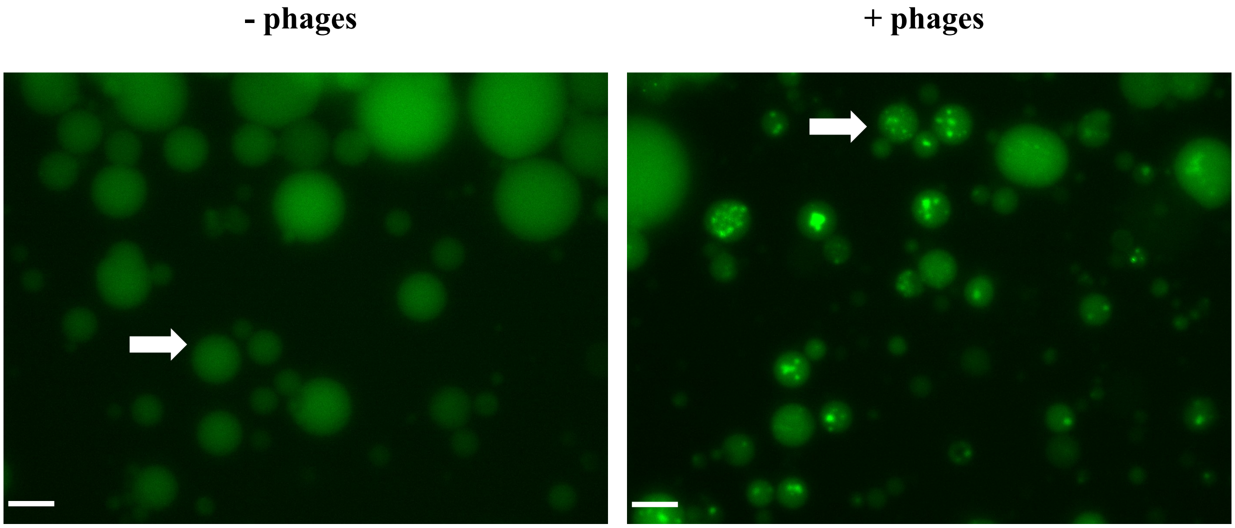


**Fig. S20. Uncropped microscopy images corresponding to Figure 3H (top).** The green channel shows Sybr Gold encapsulated into RdLPS SCs. Images were acquired 8 h after adding T7-mC-S* to the outer solution. The same batch of liposomes was divided into two wells, with (left) or without (right) phage in the outer solution. White arrows indicate the zoomed-in liposomes. Images were acquired using identical microscope settings and displayed at the same intensity levels Scale bar: 30 µm.

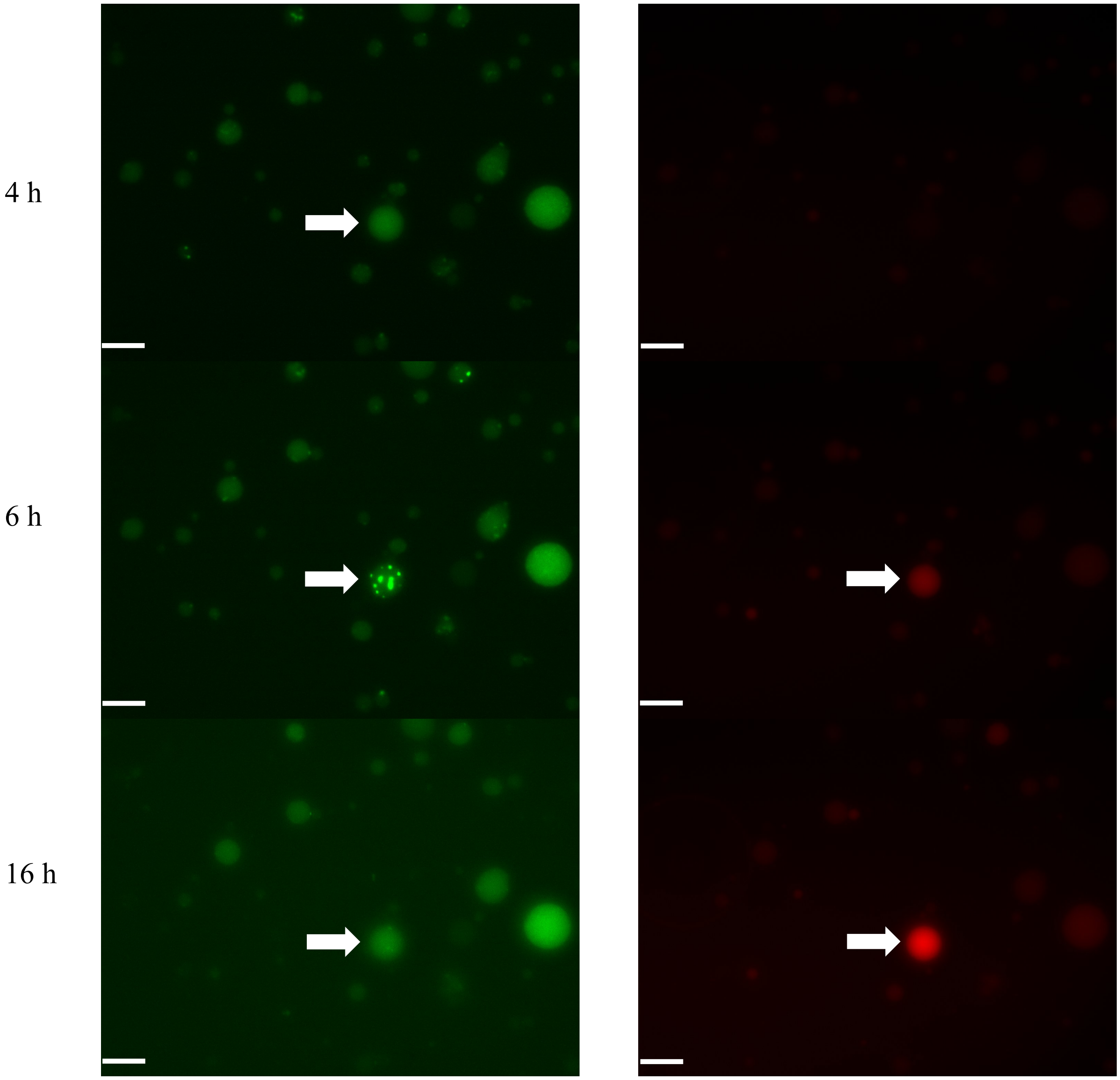


**Fig. S21. Uncropped microscopy images corresponding to Figure 3H (bottom).** The red signal represents T7-mC-S* fluorescence, the green channel shows SYBR Gold encapsulated in RdLPS SCs. Time is indicated in hours after the addition of T7-mC-S* to the outer solution. White arrows indicate the zoomed-in SCs. Red and green channels were acquired using identical microscope settings and displayed at the same intensity levels. Scale bar: 30 µm.

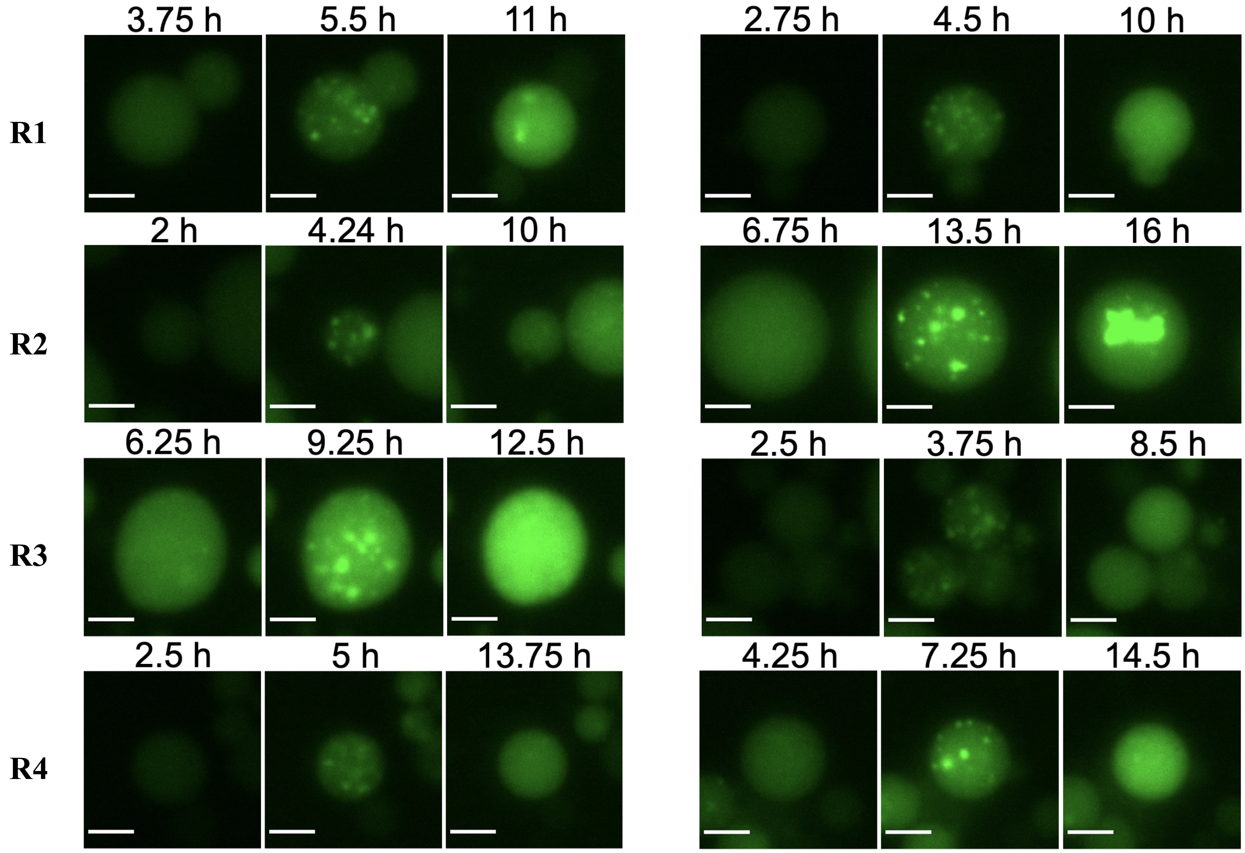


**Fig. S22. Supplement SCs for Figures 3H and 3I, four replicates (R1, R2, R3, R4).** Green channel of representative RdLPS SCs encapsulating Sybr Gold DNA intercalating dye incubated in an outer solution containing T7-mC-S*. Two liposomes are selected from each replicate experiment and three timepoints are shown. The first timepoint corresponds to the liposome before phage infection, the second timepoint to green clusters formation and the third timepoint to cluster disassembly. All images were acquired with the same microscope settings and are displayed at the same intensities. Time is indicated in hours after the start of the experiment. Scale bar is 20 µm.

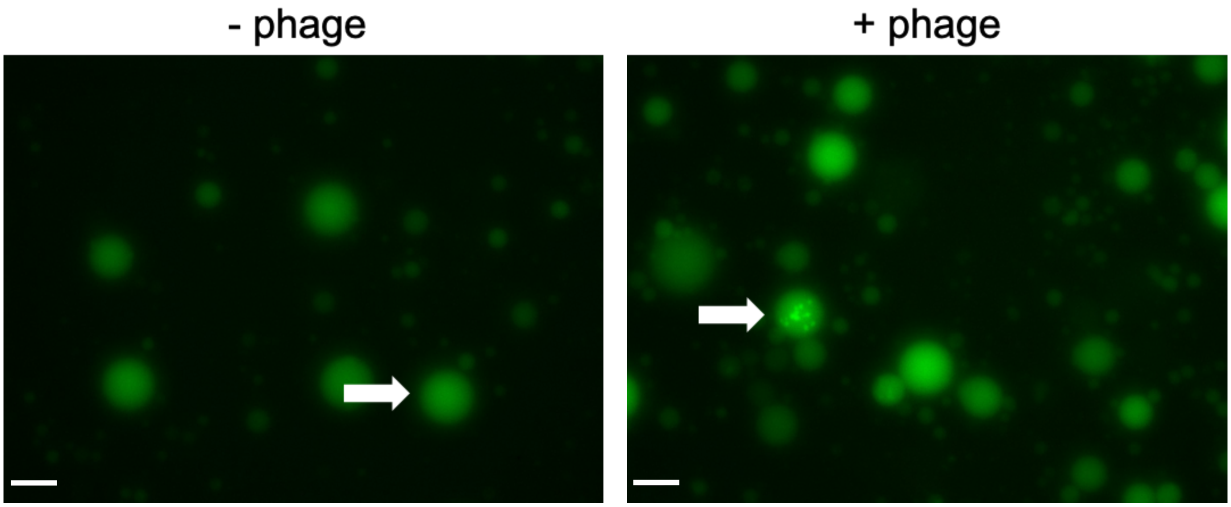


**Fig. S23. Uncropped images (green channel) from Figure 3J.** Picture correspond to the same preparation of RdLPS SCs incubated with or without T7-mC-S* in the outer solution. Images were acquired with the same microscope settings and are displayed at the same intensity levels. A white arrow points at the liposomes at which the zoom was done. Pictures are taken after 11 h of incubation, scale bar is 30 µm.

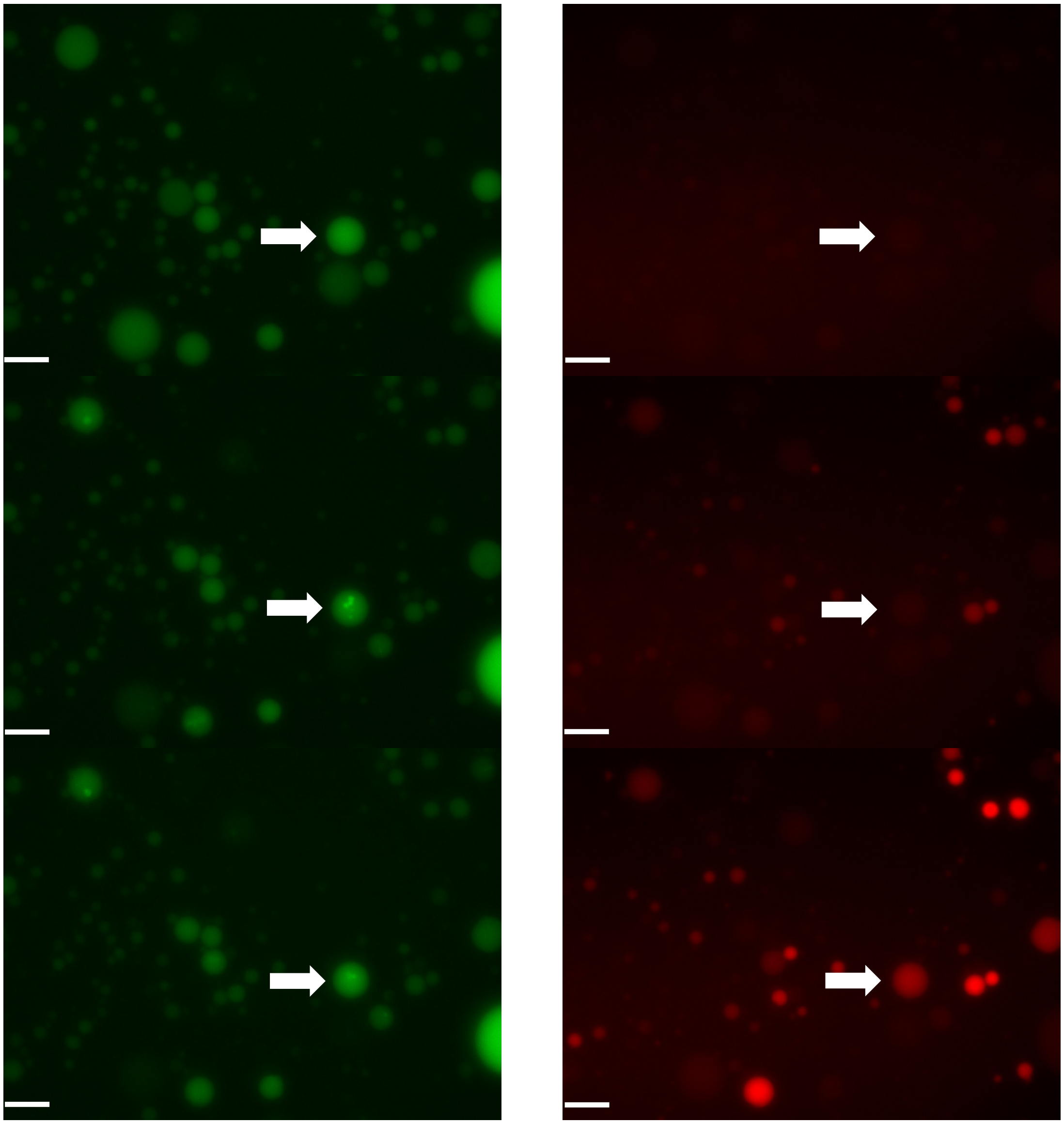


**Fig. S24. Uncropped microscopy Images corresponding to Figure 3J (bottom).** Red signal corresponds to T7-mC-S* fluorescence, green channel to dCTP-FITC encapsulated into RdLPS SCs. Time is indicated in hours after the addition of T7-mC-S* in the outer solution. White arrows indicate the SCs that was zoomed in. Red and green channels were acquired using identical microscope settings and displayed at the same intensity levels. Scale bar is 30 µm.

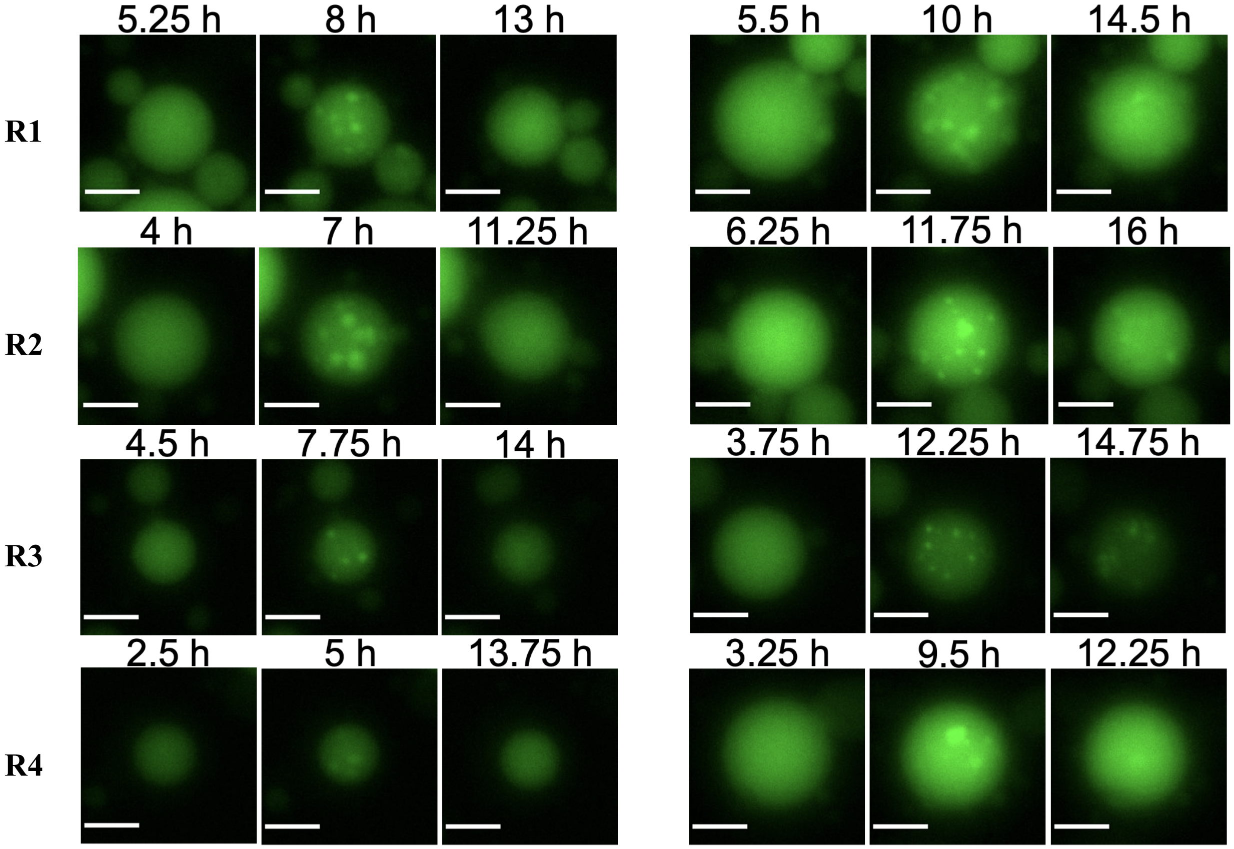


**Fig. S25. Supplement SCs for Figures 3J and 3K, four replicates (R1, R2, R3, R4).** Green channel of representative RdLPS SCs encapsulating dCTP-FITC fluorescent nucleotide incubated in an outer solution containing T7-mC-S*. Two SCs are selected in each replicate experiment and three timepoints are shown. The first timepoint corresponds to the SC before phage infection, the second timepoint to green clusters formation and the third timepoint to cluster disassembly. All images were acquired with the same microscope settings and are displayed at the same intensities. Time is indicated in hours after the start of the experiment. Scale bar is 20 µm.

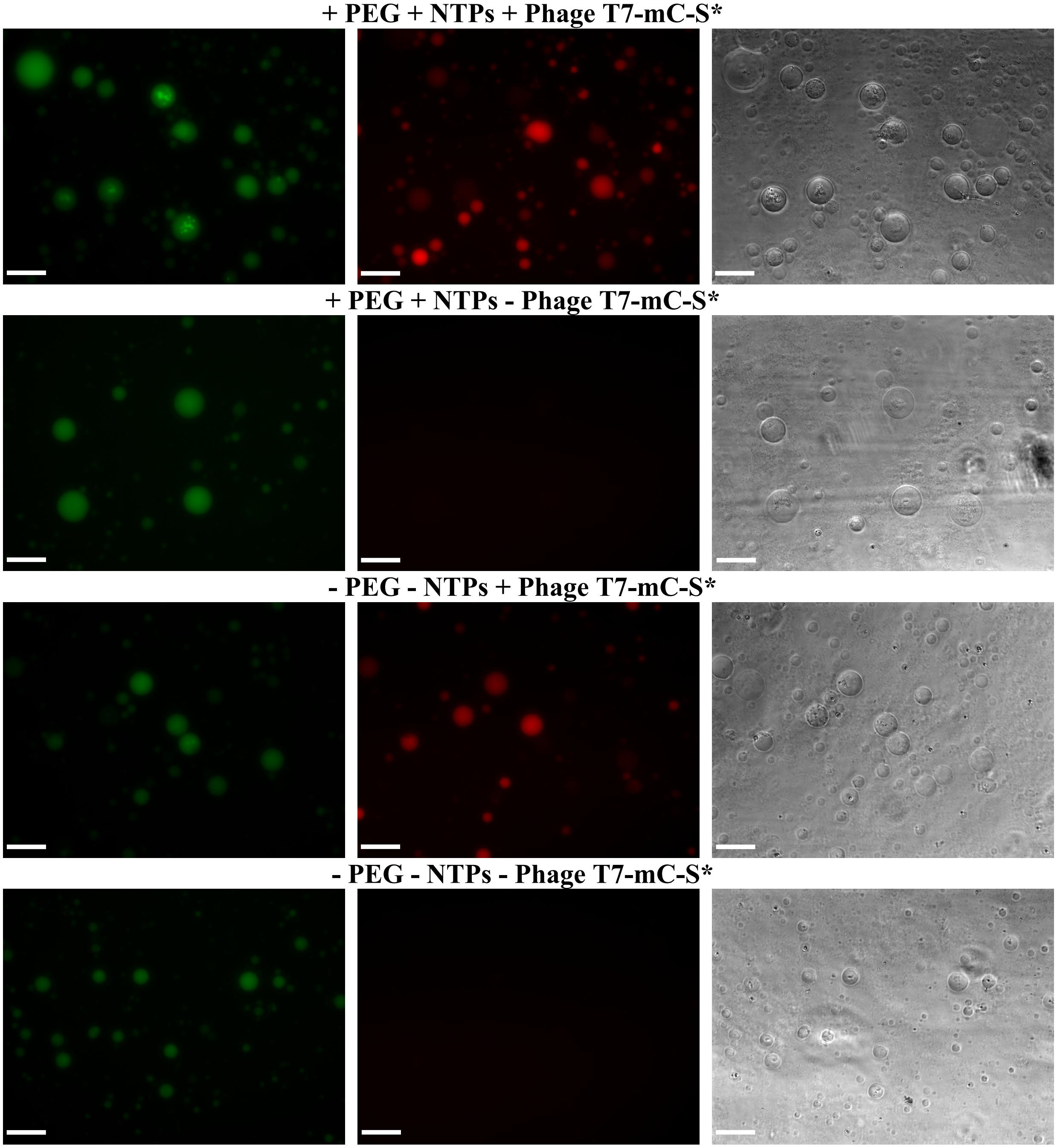


**Fig. S26. Microscopy images illustrating infection of liposomes with or without PEG and dNTPs.** Endpoint images of RdLPS SCs after 15 hours of incubation, red channel corresponds to T7-mC-S* fluorescence and green channel to dCTP-FITC fluorescence. In the absence of phage in the outer solution, no red fluorescence and no green clusters are observed in the liposomes. For each channel, images were acquired with the same microscope settings and are displayed at the same intensities. Scale bar is 40 µm.

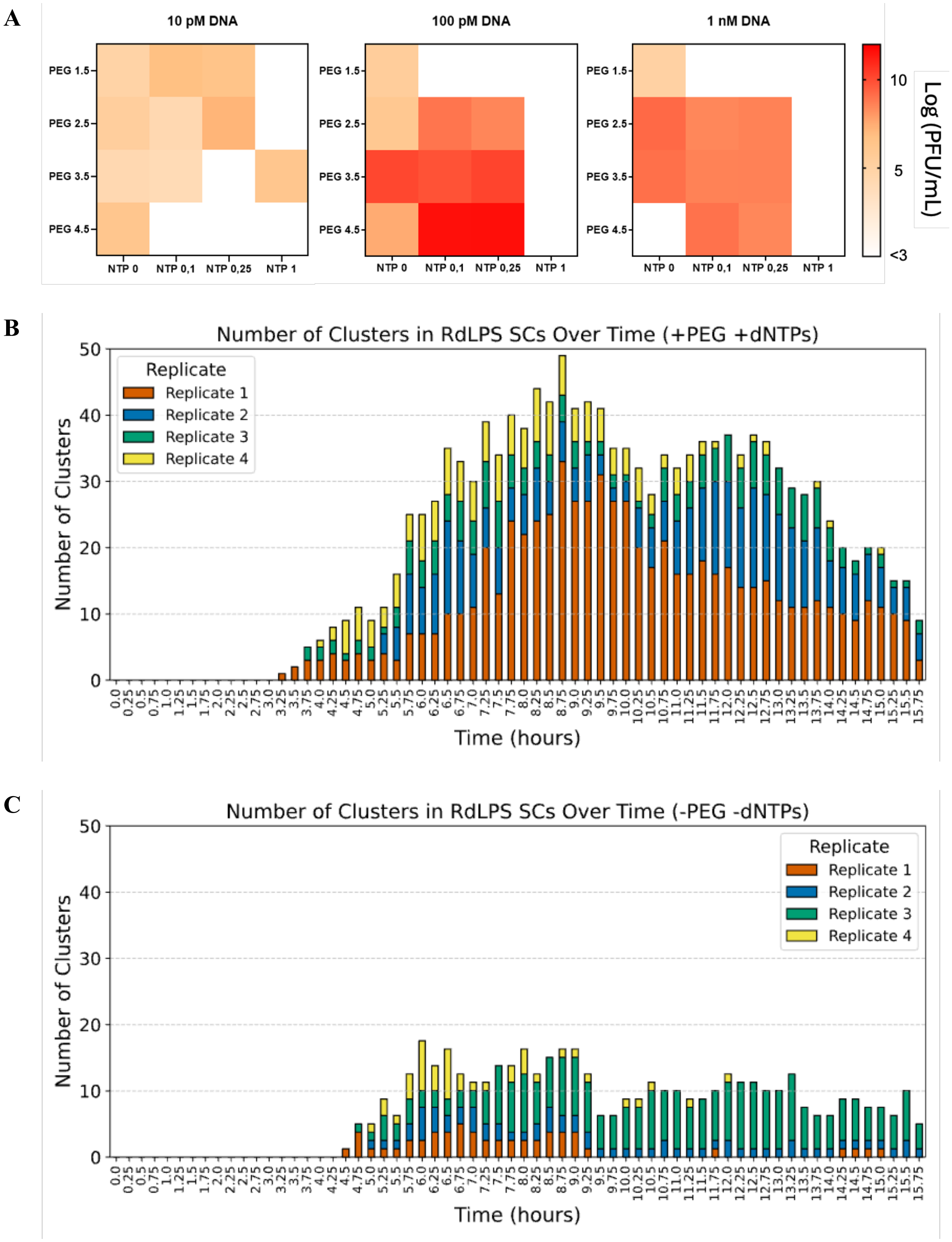
A
10 pM DNA
100 pM DNA
1 nM DNA
PEG 1.5
PEG 2.5
PEG 3.5
PEG 4.5
NTP 0
NTP 0,1
NTP 0,25
NTP 1
10
5
<3
Log (PFU/mL)
B
Number of Clusters in RdLPS SCs Over Time (+PEG +dNTPs)
Replicate
Replicate 1
Replicate 2
Replicate 3
Replicate 4
Number of Clusters
Time (hours)
C
Number of Clusters in RdLPS SCs Over Time (-PEG -dNTPs)
Replicate
Replicate 1
Replicate 2
Replicate 3
Replicate 4
Number of Clusters
Time (hours)

**Fig. S27. Effect of PEG and dNTPs on CFE phage yield. A.** Bulk CFE production of T7 phage at three initial concentrations for varying concentrations of PEG and dNTPs. CFE reactions were incubated for 12 h and titer were recorded by serial dilution spotting. We observed that presence of both dNTPs and PEG increase phage yield and that 0.1 nM maximize phage yield. **B.** Plot from Figure 2K, data from the four different replicates are represented. This data corresponds to average cluster formation in presence of 3.5% PEG and 0.1 mM dNTPs for four replicate experiments. **C.** Plot for cluster formation in absence of PEG and dNTPs. We observe that less cluster are formed in absence of PEG and dNTPs. Without phage in the outer solution, no cluster is observed in the liposomes. Cluster detection analysis is described in methods section.

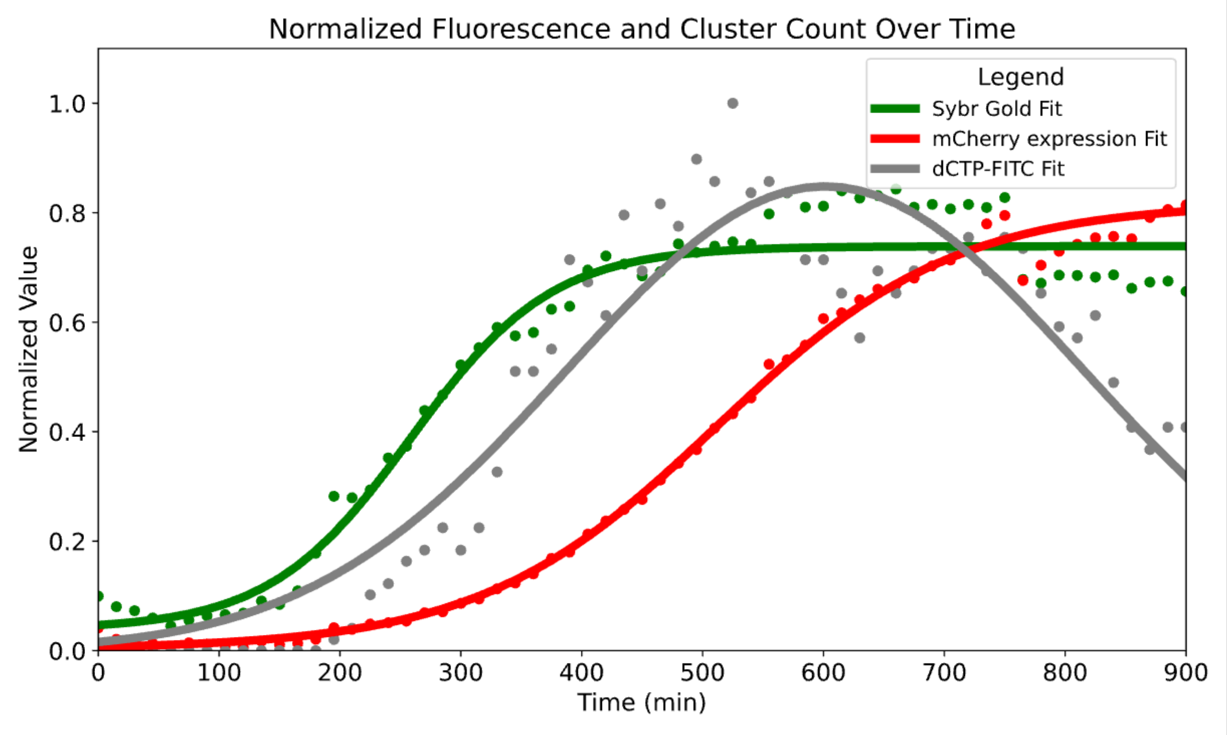


**Fig. S28. Supplement for Figure 4A.** Same plot as in Figure 4A including all data points. Green and red data points correspond to normalized mean fluorescence for Sybr Gold and mCherry fluorescence signals in liposomes. Grey Data points correspond to normalize cluster counts for dCTP-FITC signal. Liposome analysis is described in the method section. Sigmoidal fits were selected for green and red signal and a gaussian fit was selected for cluster formation.

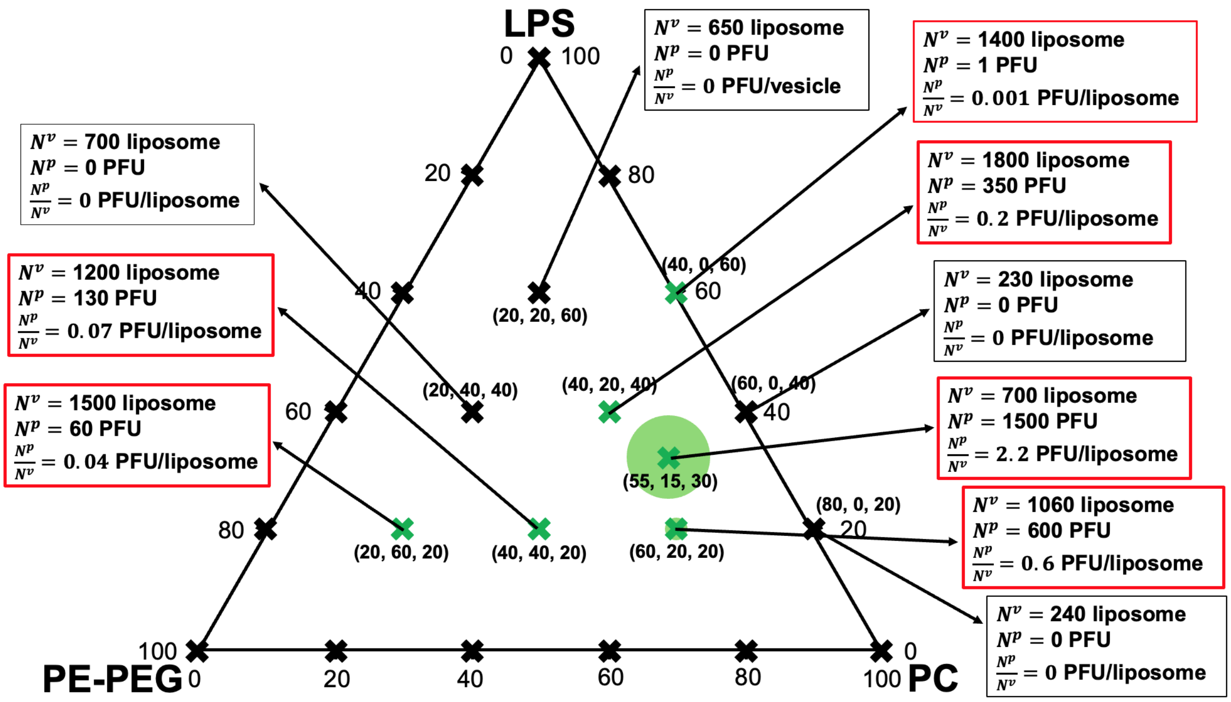


**Fig. S29. Ternary diagram of T7-mC-S* production from encapsulated genomes in RdLPS SCs.** The ternary diagram represents POPC/PEG-PE/RdLPS at a total lipid concentration of 100 µM in oil, using the same compositions as in Fig. S3. The T7-mC-S* genome was encapsulated at 0.1 nM in RdLPS SCs and incubated at 30°C for 15 h. SC count was estimated for each condition, along with PFU measurements obtained by SC lysis in water followed by spotting on an *E. coli* B lawn. The composition 55/15/30 (POPC/PEG-PE/RdLPS) yielded the highest PFU per liposome.

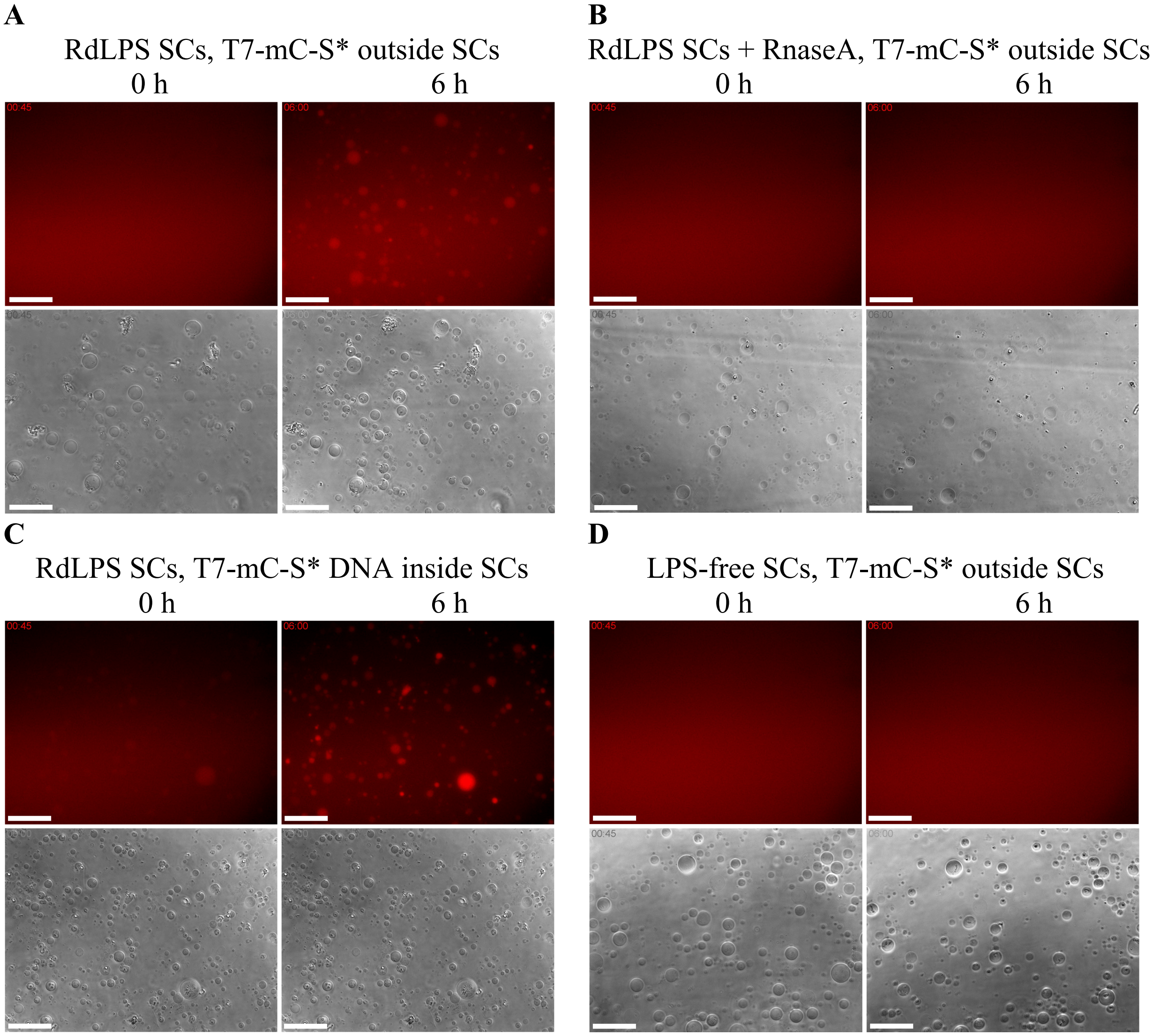


**Fig. S30. RdLPS SCs infection controls.** Four populations of liposomes bright field and red channel microscopy images acquired with the same microscope settings and displayed at the same red intensity after 45 min and 6 h of incubation. **A.** RdLPS SCs encapsulating an active CFE reaction incubated with phage T7-mC-S* added to the outer solution. A red fluorescence signal is observed in liposomes indicating infection. **B.** RdLPS SCs encapsulating a dead CFE reaction. RNAseA was added to the CFE reaction to inactivate the phage production. Phage T7-mC-S* was added to the outer solution. No red signal is observed after 6 h in the liposomes. **C.** RdLPS SCs encapsulating a CFE reaction and T7-mC-S* genome at 0.1 nM, incubated in an outer solution devoid of phages. A red fluorescence signal is observed in liposomes indicating T7-mC-S* genome expression. **D.** RdLPS-free SCs encapsulating an CFE reaction incubated with phage T7-mC-S* added to the outer solution. No red fluorescence signal is observed in liposomes indicating no infection. Scale bar is 50 μm.

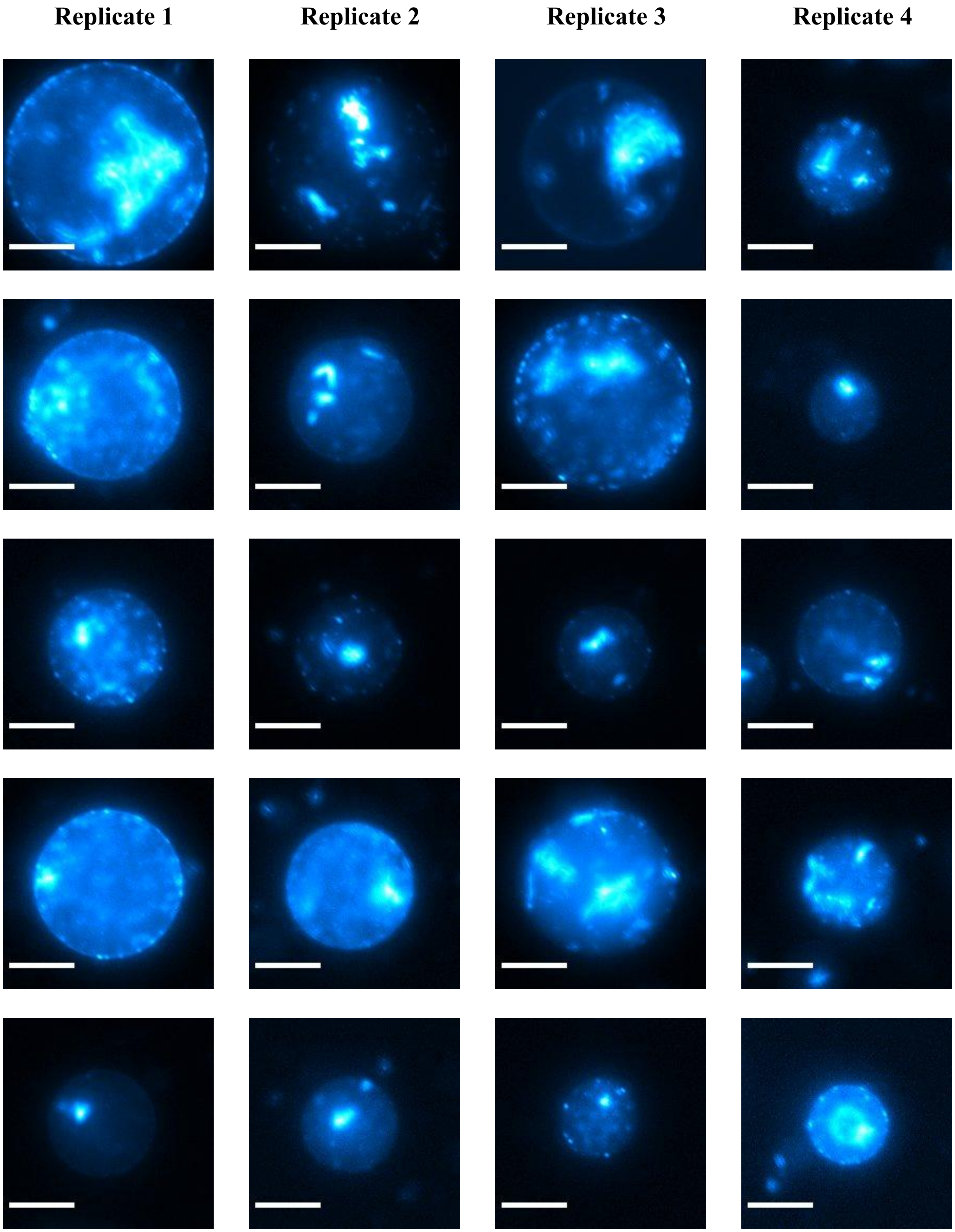
Replicate 1
Replicate 2
Replicate 3
Replicate 4

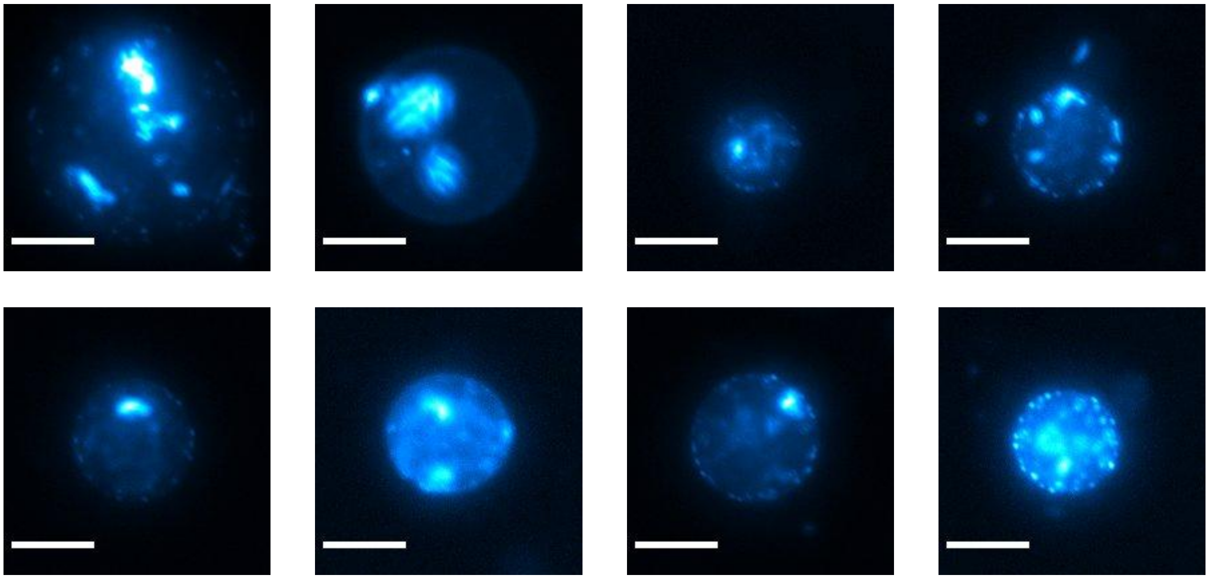

**Fig. S31. Supplementary SCs corresponding to Figure 4E.** Green fluorescence signal of RdLPS SCs infected by T7-Split-S* at an MOI of 100 after 16 h of incubation. The signal corresponds to Split-GFP expression. Seven representative SCs from four independent experiments are shown. Fluorescence signal appears as large clusters at concatemer loci, along with smaller dispersed clusters, as observed in Figure 3E. Liposome images were each adjusted for brightness and contrast and are artificially colored using pseudo-colors from ImageJ (Cyan Hot). All images were acquired using identical microscope settings. Scale bar: 20 µm.

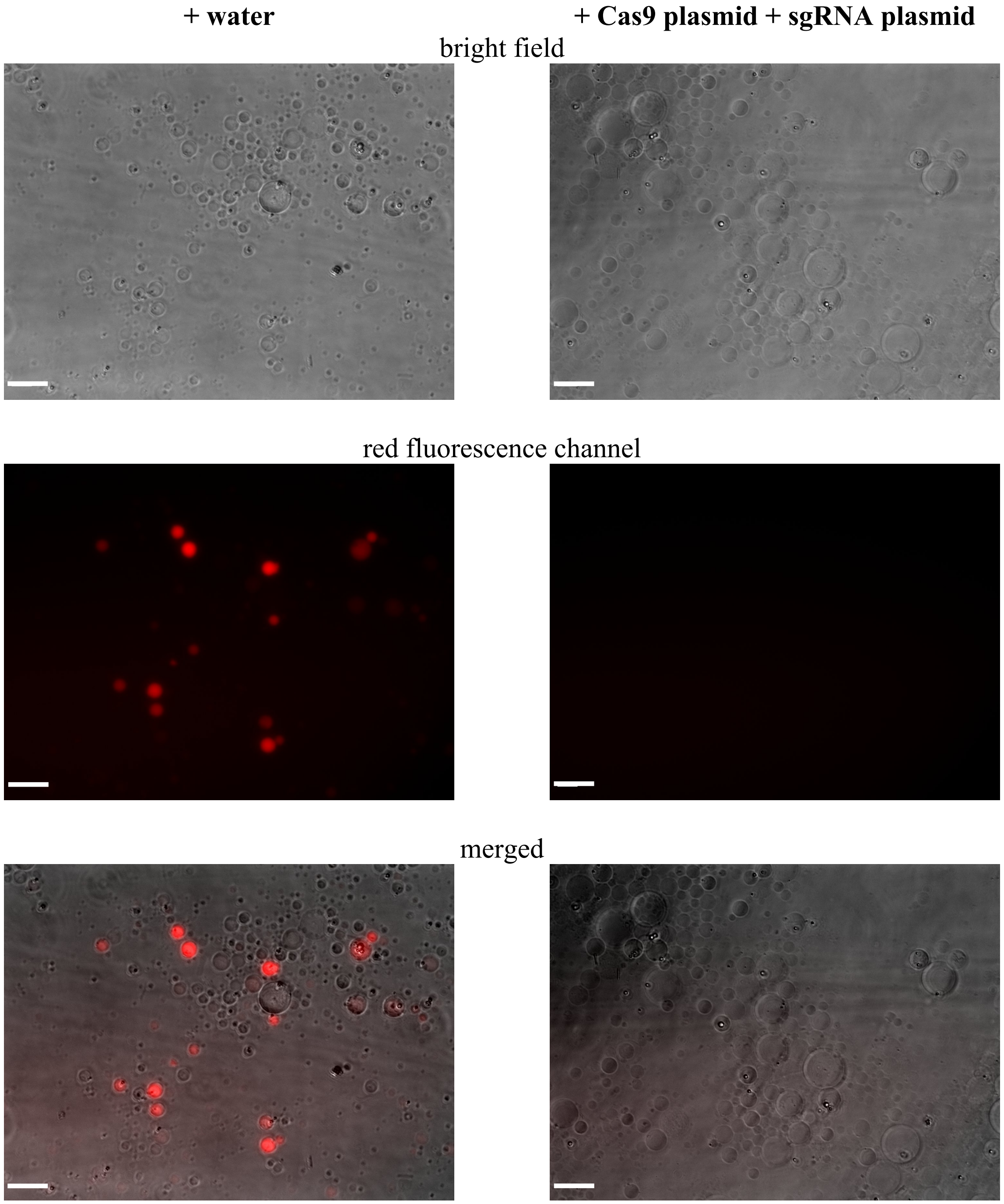


**Fig. S32. CRISPR-mediated abortion of T7-mC-S* infection in RdLPS SCs.** Merged and separated channel images of SCs encapsulating CFE with either water or a CRISPR plasmid and guide RNA targeting the *mcherry* gene in T7-mC-S*. Images show liposomes after 16 h of incubation in the presence of T7-mC-S* in the outer solution at an MOI of 100. No red fluorescence is observed in CRISPR-inhibited liposomes, indicating aborted infection. Red signals were acquired with the same microscope settings and displayed at the same intensity. Scale bar: 20 µm.

| Figures | SCs | Encapsulated Solution | Outer solution |
|---|---|---|---|
| Figure 2A | RdLPS SCs | CFE+ Rhodamine-dextran (10 μM) | + nothing |
| Figure 2C, S5 | RdLPS SCs | CFE+ Rhodamine-dextran (10 μM) | + GFP-TF* (200 nM) |
| | no LPS SCs | CFE + water | + GFP-TF* (200 nM) |
| Figure 2D, S12, S13 | RdLPS SCs | CFE + P70a-*T7rnap* (0.1 nM) + T7-*mcherry* (2 nM) | + GFP-TF* (200 nM) |
| Figure 2E, S14 | RdLPS SCs | CFE + water | + T7-SPLIT-S* ($10^{11}$ PFU/mL) |
| Figure 3B, S15 | RdLPS SCs | CFE + water | + T7-mC-S* ($10^{7}$ PFU/mL) |
| Figure 3C | RdLPS SCs | CFE + water | + T7-mC-S* ($10^{7}$ PFU/mL) + GFP-TF* (200 nM) |
| Figure 3D, S17, S18 | RdLPS SCs | CFE + water | + T7-mC-WT/T7-mC-S*/T7-WT/T7-S* ($10^{7}$ PFU/mL) |
| Figure 3E-F, S19 | RdLPS SCs | CFE + FITC-dextran (10 μM) | + T7-mC-S* ($10^{8}$, $10^{7}$, $10^{6}$, $10^{5}$, $10^{4}$, $10^{3}$ PFU/mL) |
| Figure 3G | RdLPS SCs | CFE + P70a-*T7rnap* (0.1 nM) + T7-*mcherry* (2 nM) | + nothing |
| | RdLPS SCs | CFE + water | + T7-mC-S*($10^{7}$ PFU/mL) |
| Figure 3H-I, S20-22 | RdLPS SCs | CFE + Sybr Gold (1X) | + T7-mC-S*($10^{7}$ PFU/mL) |
| Figure 3J-K, S23-25 | RdLPS SCs | CFE + dCTP-FITC (10 μM) + 2% PEG 8k + dNTPs (0.1 mM each) | + T7-mC-S*($10^{7}$ PFU/mL) |
| Figure 4C | RdLPS SCs | CFE + T7 WT DNA (0.01, 0.1, 1 nM) + Rhodamine-dextran (10 μM) | + RNAse A (10 μg/mL ) |
| Figure 4D | RdLPS SCs | CFE + T7 WT DNA or T7-mC-S* DNA (0.01 nM) + Rhodamine-dextran (10 μM) | + RNAse A (10 μg/mL ) |
| | no LPS SCs | CFE + T7 WT DNA or T7-mC-S* DNA (0.05 nM) + Rhodamine-dextran (10 μM) | + RNAse A (10 μg/mL ) |
| Figure 4E, S31 | RdLPS SCs | CFE + Water | + T7-SPLIT-S* ($10^{7}$ PFU/mL) |
| Figure 4F | RdLPS SCs | CFE + P70a-*T7rnap* (0.1 nM) + T7-holin (5 nM) + Rhodamine-dextran (10 μM) | + nothing |

| | | | |
|---|---|---|---|
| | no LPS SCs | CFE + P70a-*T7rnap* (0.1 nM) + T7-holin (5 nM) + Rhodamine-dextran (10 μM) | + nothing |

**Table S1.**
Experimental conditions – Main Figures

| Figures | SCs | Encapsulated Solution | Outer solution |
|---|---|---|---|
| S3 | variable RdLPS SCs | CFE+ water | + GFP-TF* (200 nM) |
| S4 | RdLPS SCs | CFE+ water | + GFP-TF* or GFP-TF (200 nM) |
| | no LPS SCs | CFE+ water | + GFP-TF* or GFP-TF (200 nM) |
| S6 | RdLPS SCs | CFE+ water | + GFP-TF* or GFP-TF or sfGFP (200 nM) |
| | no LPS SCs | CFE+ water | + GFP-TF* or GFP-TF or sfGFP (200 nM) |
| S7 | RdLPS SCs | CFE+ water | + GFP-TF* (200 nM) |
| S8 | RdLPS SCs | CFE+ water | + T7-mC-S* ($10^7$ PFU/mL) + GFP-TF* (200 nM) |
| S9, S10, S11 | RdLPS SCs | CFE+GFP-TF* (200 nM) | + nothing |
| | RdLPS SCs | CFE+GFP-TF* (200 nM) | + GFP-TF* (200 nM) |
| | no PE-PEG RdLPS SCs | CFE+GFP-TF* (200 nM) | + nothing |
| | no PE-PEG RdLPS SCs | CFE+GFP-TF* (200 nM) | + GFP-TF* (200 nM) |
| S16 | variable RdLPS SCs | CFE+ water | + T7-mC-S* ($10^7$ PFU/mL) |
| S26-27 | RdLPS SCs | CFE+ 2% PEG 8k + 0.1 mM dNTPs + 10 µM FITC-dCTP | + T7-mC-S* ($10^7$ PFU/mL) |
| | RdLPS SCs | CFE+ 2% PEG 8k + 0.1 mM dNTPs + 10 µM FITC-dCTP | + nothing |
| | RdLPS SCs | CFE + 10 µM FITC-dCTP | + T7-mC-S* ($10^7$ PFU/mL) |
| | RdLPS SCs | CFE + 10 µM FITC-dCTP | + nothing |
| S29 | variable RdLPS SCs | CFE + T7-mC-S* DNA (0.1 nM) + FITC-dextran (10 µM) | + RNAse A (10 µg/mL ) |
| S30 | RdLPS SCs | CFE + water | + T7-mC-S* ($10^7$ PFU/mL) |
| | RdLPS SCs | CFE + RNAse A (10 µg/mL) | + T7-mC-S* ($10^7$ PFU/mL) |
| | RdLPS SCs | CFE + T7-mC-S* DNA (0.1 nM) | + nothing |
| | no LPS SCs | CFE +water | + T7-mC-S* ($10^7$ PFU/mL) |
| S32 | RdLPS SCs | CFE +water | + T7-mC-S* ($10^7$ PFU/mL) |

| | | | |
|---|---|---|---|
| | RdLPS SCs | CFE + Cas9 plasmid (1 nM) + *mcherry* sgRNA plasmid (1 nM) | + T7-mC-S* ($10^7$ PFU/mL) |

**Table S2.**

Experimental conditions – Supplementary Figures

| fragments | size (bp) | primer forward | primer reverse | template | accession number |
|---|---|---|---|---|---|
| T7A | 10680 | T7A-s | T7A-as | T7 WT | V01146.1 |
| T7B | 9864 | T7B-s | T7B-as | T7 WT | V01146.1 |
| T7C1 | 2778 | T7C1-s | T7C1-as | T7 WT | V01146.1 |
| gp10-GFP11-GFP1-10 | 1722 | gp10-SPLIT-s | gp10-SPLIT-as | gp10b-GFP11-RBS-GFP1-10 | synthetic DNA |
| T7C2 | 6345 | T7C2-s | T7C2-as | T7 WT | V01146.1 |
| T7D-RdLPS | 9429 | T7D-s | T7D-as | rfaC_2 | PP384400 |

**Table S3.**
T7-Split-S* genome assembly map.

| fragments | size (bp) | primer forward | primer reverse | template | accession number |
|---|---|---|---|---|---|
| T7A | 10680 | T7A-s | T7A-as | T7 WT | V01146.1 |
| T7B | 9864 | T7B-s | T7B-as | T7 WT | V01146.1 |
| T7C-mC | 10114 | T7C-s | T7C-as | T7_mC-g10-o | PP384393 |
| T7D-RdLPS | 9429 | T7D-s | T7D-as | rfaC_2 | PP384400 |

**Table S4.**
T7-mC-S* genome assembly map.

**Movie S1.**

Four representative timelapse microscopy of RdLPS SCs in presence of T7-mC-WT, T7-mC-S*, T7-WT or T7S* in the outer medium. Images correspond to the superposition of the red fluorescence channel and bright field.

**Movie S2.**

Four representative timelapse microscopy of RdLPS SCs in presence of T7-mC-S* at an MOI of 3, 30, 300 or 3000 in the outer medium. Images correspond to the superposition of the red fluorescence channel and bright field.

**Movie S3.**

Two representative green fluorescence timelapse microscopy of RdLPS SCs containing Sybr Gold with or without T7-mC-S* in the outer medium.

**Movie S4.**

Two representative green fluorescence timelapse microscopy of RdLPS SCs containing FITC-dCTP with or without T7-mC-S* in the outer medium.

**Movie S5.**

Two representative timelapse microscopy of RdLPS SCs incubated with T7-Split-S* in the outer medium. Green channel and bright field are represented side by side.

**Movie S6.**

Four representative red fluorescence timelapse microscopy of liposomes with or without RdLPS containing a red fluorescent dye loaded CFE reaction with or without a T7 holin gene circuit.

**Data S1. (separate file)**

DNA sequences (primers, gblocks, plasmids and T7 genomes) used in this work.
